# Bacterial pathogen deploys iminosugar galactosyrin to manipulate plant glycobiology

**DOI:** 10.1101/2025.02.13.638044

**Authors:** Nattapong Sanguankiattichai, Balakumaran Chandrasekar, Yuewen Sheng, Nathan Hardenbrook, Werner W. A. Tabak, Daniel Krahn, Margit Drapal, Pierre Buscaill, Suzuka Yamamoto, Atsushi Kato, Robert Nash, George Fleet, Paul Fraser, Markus Kaiser, Peijun Zhang, Gail M. Preston, Renier A. L. van der Hoorn

**Author notes:** Present address: Department of Biological Sciences, Birla Institute of Technology and Science, Pilani; Pilani, India (Balakumaran Chandrasekar).

## Abstract

The extracellular space (apoplast) of plants is an important molecular battleground during infection by many pathogens. We previously found that a plant-secreted β-galactosidase BGAL1 acts in immunity by facilitating the release of immunogenic peptides from bacterial flagellin and that *Pseudomonas syringae* suppresses this enzyme by producing a small molecule inhibitor called galactosyrin. Here, we elucidated the structure and biosynthesis of galactosyrin and uncovered its multifunctional roles during infection. Structural elucidation by cryo-EM and chemical synthesis revealed that galactosyrin is an iminosugar featuring a unique geminal diol attached to the pyrrolidine moiety that mimics galactose binding to the β-galactosidase active site. Galactosyrin biosynthesis branches off from purine biosynthesis and involves three enzymes of which the first is a reductase that is unique in iminosugar biosynthesis. Besides inhibiting BGAL1 to avoid detection, galactosyrin also changes the glycoproteome and metabolome of the apoplast. The manipulation of host glycobiology may be common to plant-associated bacteria that carry putative iminosugar biosynthesis clusters.

## INTRODUCTION

The extracellular space in plant tissues (the apoplast) is an important molecular battleground during plant-pathogen interactions (*1*). This microenvironment is colonized by bacteria, fungi and oomycetes that must have evolved various strategies to avoid recognition, suppress immune responses and manipulate host physiology. Yet most of these apoplastic plant-pathogen interactions remain to be elucidated. Our previous work on the interaction between *Nicotiana benthamiana* plants and the model bacterial pathogen *Pseudomonas syringae* revealed the role of plant apoplastic β-galactosidase BGAL1 in plant immunity (*2*). BGAL1 initiates the hydrolytic release of immunogenic peptides from glycosylated flagella of *P. syringae* that activate plant defences (*2*). Interestingly, we also found that during infection, *P. syringae* pv. *tomato* DC3000 produces a small molecule inhibitor of BGAL1, which we named galactosyrin (*2*). In this work, we report the molecular structure of galactosyrin and its full biosynthesis pathway. This molecule represents a novel iminosugar class and has multifunctional roles in manipulating the extracellular glycoproteome and metabolome during infection.

## RESULTS

### Galactosyrin biosynthesis gene cluster expression is controlled by virulence gene regulators

To identify genes required for galactosyrin biosynthesis, we transformed *P. syringae* pv. *tomato* DC3000 Δ*hopQ1-1* (*3*) (called wild-type for galactosyrin (WT) in this work) with *lacZ* encoding the β-galactosidase from *Escherichia coli*, which is routinely used for blue staining with X-gal (5-bromo-4-chloro-3-indoyl-β-D-galactopyranoside). We then performed Tn5-transposon mutagenesis and selection on virulence-inducing medium containing X-gal to identify darker blue colonies of mutants that cannot produce galactosyrin to inhibit LacZ (**Fig. 1A**). The loss of galactosyrin was confirmed in activity assays with purified LacZ and a fluorogenic substrate (**Fig. S1**) and transposon insertion sites were identified for 140 galactosyrin-deficient mutants (**Table S1**). These Tn5 insertion sites concentrated in four virulence gene regulators (*hrpR, hrpS, hrpL* and *rhpS*) and one putative galactosyrin biosynthesis gene cluster (*gsn*, locus tags PSPTO_0834-8, new locus tags PSPTO_RS04425-RS04445, **Fig. 1B**).

**Fig. 1.**
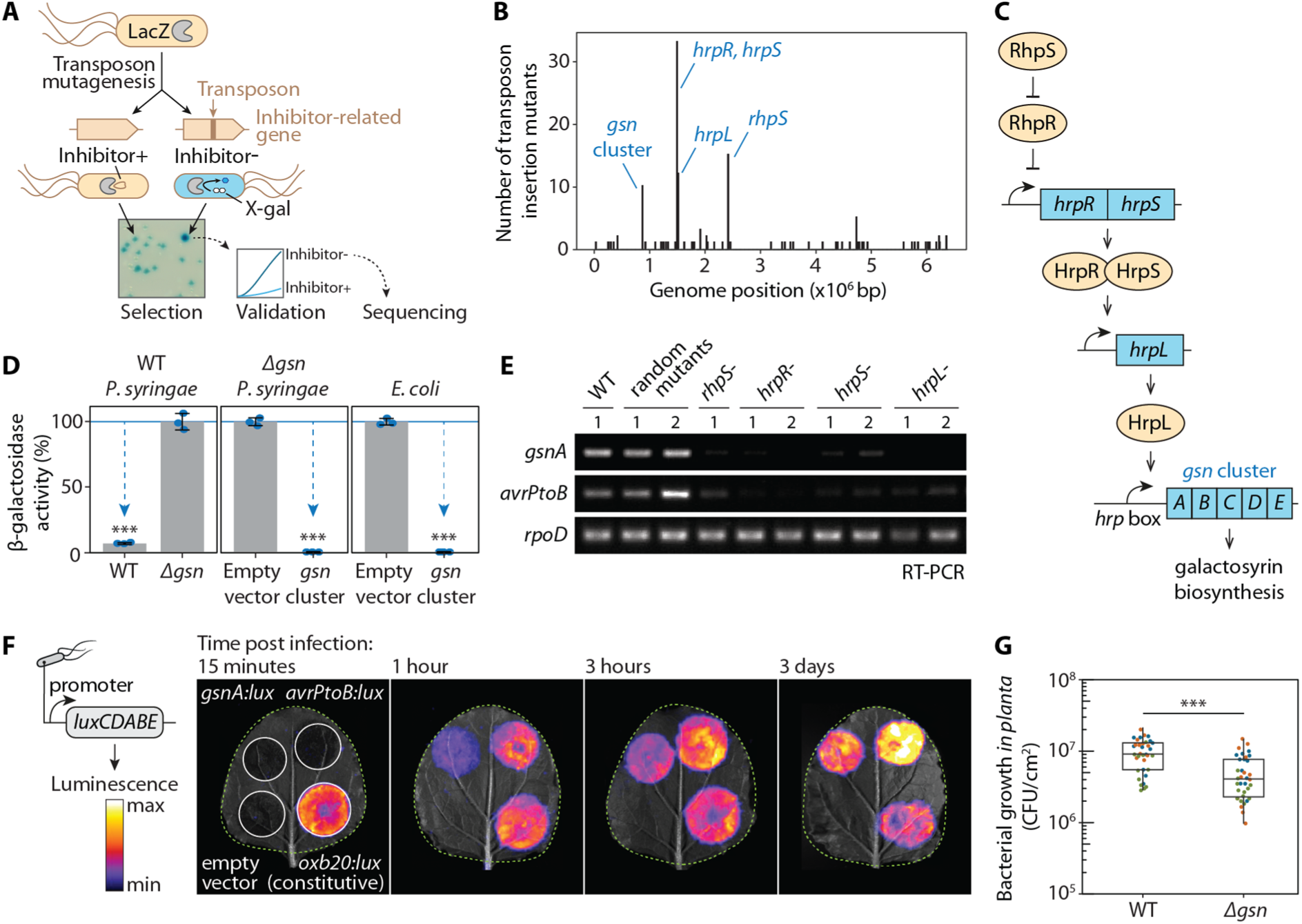
Galactosyrin biosynthesis gene cluster and its regulators identified by forward genetics. **(A)** Genetic screen for galactosyrin-deficient mutants. *P. syringae* expressing LacZ β-galactosidase was used to create a random transposon insertion mutant library. When plated onto a virulence-inducing medium supplemented with X-gal, galactosyrin-deficient mutants cannot inhibit LacZ, resulting in a darker blue colour. These candidate mutants were validated in an enzymatic assay for the inability to produce galactosyrin. Confirmed mutants were sequenced to identify transposon insertion sites. **(B)** Histogram with number of transposon insertion sites identified from galactosyrin mutants along the position within the genome, showing four hotspots corresponding to the *gsn* gene cluster and virulence regulators *hrpR*, *hrpS*, *hrpL* and *rhpS*. **(C)** Summary of the roles of genes required for galactosyrin production. The *gsn* cluster (containing five genes *gsnABCDE,* PSPTO0834-8) confers galactosyrin biosynthesis. The expression of the *gsn* cluster is controlled by a regulatory cascade of type III secretion system regulators (RhpS, HrpR, HrpS and HrpL), which controls virulence gene induction during infection. The promotor of the *gsn* cluster contains the binding site of the HrpL transcriptional activator (*hrp* box). **(D)** The *gsn* cluster confers galactosyrin biosynthesis in *P. syringae* and *E. coli*. Bacterial strains were grown in virulence-inducing medium and the supernatant was tested for LacZ inhibition using purified LacZ and substrate FDG (Fluorescein di(-β-D-Galactopyranoside). β-galactosidase activity is reported as a percentage of the activity relative to the mean of the no-inhibitor-control (*`Δ`gsn* or empty vector). Arrows highlight significant inhibition. Error bars represent standard deviation from 3 replicates. Asterisks indicate statistically significant difference compared to no-inhibitor-control (P < 0.001) using Welch’s t-test. **(E)** Expression of the *gsn* cluster is dependent on *hrpR*, *hrpS*, *hrpL* and *rhpS*. Bacterial strains were grown in virulence-inducing medium, then total RNA was extracted for reverse transcription polymerase chain reaction (RT-PCR) to monitor transcript levels of *gsnA, avrPtoB* (type III secreted effector gene) and *rpoD* (reference gene). **(F)** The *gsn* cluster is transcribed during infection. Bacteria carrying various promoter:*luxCDABE* reporter fusion constructs were infiltrated into *N. benthamiana* leaves and luminescence was imaged at different time points after infection. Signals displayed are scaled to the maximum and minimum within each image. Leaves are outlined with dashed lines. **(G)** *gsn* cluster contributes to virulence. Bacterial strains were spray-inoculated on *N. benthamiana* leaves then bacterial growth (number of bacterial colony forming units (CFU) per cm^2^ of leaf) was quantified at 3 days post infection. Results from 3 independent experiments with 12 replicates each are plotted in different colours. Asterisks indicate statistically significant difference between strains (P < 0.001) using two-way ANOVA with experiments as blocks.

The *gsn* cluster contains five genes encoding three biosynthesis enzymes (GsnA/B/C), a protein of unknown function (GsnD), and a transporter (GsnE) (**Fig. 1C**). The deletion mutant lacking the *gsn* cluster (*`Δ`gsn*) is unable to produce the inhibitor, and transformation of this mutant with a plasmid carrying the *gsn* cluster restores inhibitor production (**Fig. 1D**). Galactosyrin production was also established in *E. coli* upon transformation with the plasmid carrying the *gsn* cluster (**Fig. 1D**). These results confirm that the *gsn* gene cluster is necessary and sufficient for galactosyrin production in bacteria.

The promoter of the *gsn* gene cluster contains the *hrp* box, a conserved binding site for transcription factor HrpL (*4*), which is transcriptionally regulated by HrpR/S and RhpS (*5*) (**Fig. 1C**), RhpS, HrpR/S and HrpL are master regulators of virulence genes including type-III effectors such as *avrPtoB* (*5*). Indeed, expression of the *gsn* cluster is impaired in *rhpS, hrpR/S* and *hrpL* mutants, like *avrPtoB* (**Fig. 1E**), clarifying why these mutants are galactosyrin deficient. Consequently, as demonstrated with a *gsn:lux* reporter strain, the *gsn* cluster is transcribed from the initial to late stages of infection (**Fig. 1F**), consistent with inhibitor production during infection (*2*). When compared to WT bacteria, the *`Δ`gsn* mutant has reduced growth in *N. benthamiana* (**Fig. 1G**) but not *in vitro* (**Fig. S2**), indicating that the *gsn* cluster produces a virulence factor during infection. This is also consistent with reduced virulence described earlier for a *gsnA* mutant in *Arabidopsis thaliana* (*4*).

The *gsn* cluster is present in various strains across the major phylogroups of *P. syringae* (**Fig. S3A**), but the phylogeny of *gsnA* is incongruent with that of *P. syringae* (**Fig. S3B**). The *gsn* cluster is also flanked by transposable elements located downstream of tRNA^Lys^ loci (**Fig. S3A, C**), which are typical for integrase sites (*6*). Together with the fact that the *gsn* cluster has lower GC content than its neighboring regions and the genomic average (**Fig. S3D**), these data indicate that the *gsn* cluster has been distributed in *P. syringae* through horizontal gene transfer. Furthermore, GsnA homologs (aldehyde dehydrogenases, ADHs) are present in diverse bacterial species in different gene clusters with similar gene functions (**Fig. S4**), some of which are known to produce distinct iminosugars, potent glycosidase inhibitors with sugar-like structures containing a nitrogen instead of oxygen in the ring (**Fig. S4**) (*7–9*). However, unlike previously characterised gene clusters, the *gsn* cluster forms a distinct clade that also encodes GsnB (homolog of reductase RibD) (**Fig. S4**), suggesting that galactosyrin could be a novel iminosugar produced by a yet unknown metabolic pathway.

### Galactosyrin structure and inhibition mechanism resolved by cryo-EM

To elucidate the molecular structure of galactosyrin, we used His-tagged LacZ immobilised on a metal affinity resin to capture galactosyrin from the crude secretome of the WT strain until LacZ saturation. Subsequent washing and elution with imidazole yielded a LacZ-galactosyrin complex with a high degree of inhibitor saturation (**Fig 2A, B**). Using cryo-electron microscopy (cryo-EM), we resolved the structure of the LacZ-galactosyrin complex at 1.9 Å resolution and detected an electron density in the active site that was absent in the negative control generated using the secretome of the *`Δ`gsn* mutant (**Fig. 2C, D, S5**). This density revealed that galactosyrin consists of a five-membered ring with three chiral centers: two with putative hydroxyls and one with a putative branching geminal diol group, which likely forms by hydration of an aldehyde group (**Fig. 2D**). We next chemically synthesised this molecule **(Fig. S13)** and obtained an unprecedented 1.4 Å resolution structure of its complex with LacZ, which is identical to the native galactosyrin (**Fig. 2D, S5**), confirming the structure of galactosyrin. To the best of our knowledge, this iminosugar has not been observed and characterized before and illustrates that cryo-EM can be used to elucidate structures of novel natural products bound to their targets at atomic resolution.

**Fig. 2.**
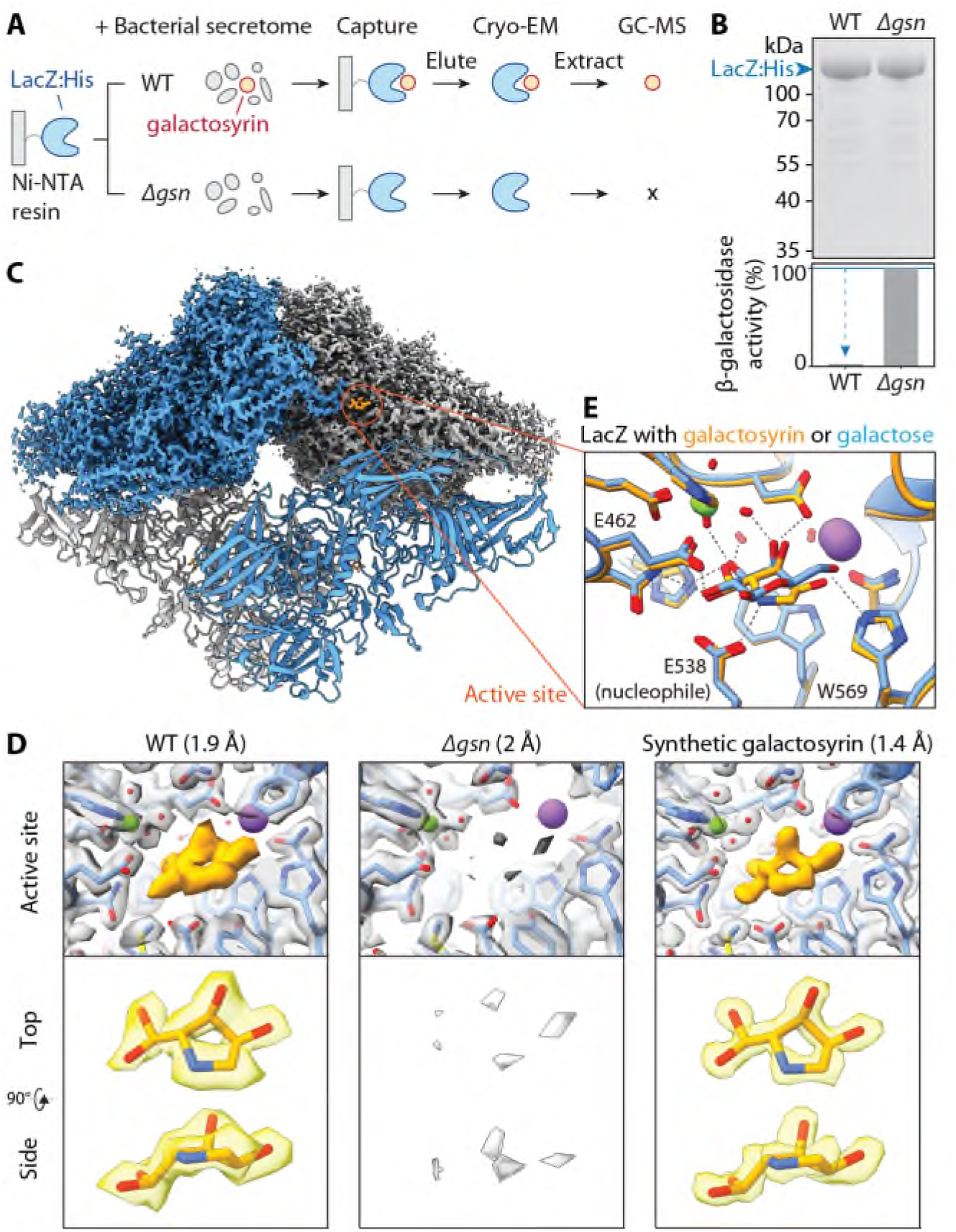
Galactosyrin is a hydrated pyrrolidine of a novel iminosugar class. **(A)** LacZ-galactosyrin complex capture and downstream analyses. A Histidine-tagged β-galactosidase enzyme from *E. coli* (LacZ:His) immobilised on Ni-NTA beads was used to capture galactosyrin inhibitor from crude bacterial secretome of galactosyrin-producing *P. syringae* (WT) or the galactosyrin-deficient mutant (*`Δ`gsn*, negative control). After washing, the complex was eluted and used for cryo-electron microscopy (Cryo-EM), and soluble metabolites were extracted for analysis by gas chromatography-mass spectrometry (GC-MS). **(B)** Captured LacZ is saturated with galactosyrin. (Top) Total protein stain of eluted samples separated on SDS-PAGE. (Bottom) β-galactosidase activity of each sample measured by FDG assay showing inhibition of WT sample compared to *`Δ`gsn*. **(C)** Structure of LacZ-galactosyrin complex from Cryo-EM. The density map is shown for the top half of the structure and a fitted model is shown for the bottom half. Each monomer of LacZ tetramer is coloured differently. **(D)** Structure of galactosyrin revealed by Cryo-EM. Top: structures of LacZ-galactosyrin complex capture from WT or *`Δ`gsn* strains and of LacZ incubated with synthetic galactosyrin. Density maps with fitted protein structures show the enzyme active site with the presence and absence of galactosyrin (orange). The resolution of each structure is shown in brackets. Bottom: extracted density map with fitted structure of galactosyrin from top and side view. **(E)** Galactosyrin mimics galactose binding in the active site. Overlay of structures of the LacZ active site and interacting residues in complex with galactosyrin (orange) or galactose (blue) showing similarity of overall binding pose and positioning of hydroxyl groups. The positive charge on the likely protonated amine nitrogen of galactosyrin can introduce extra electrostatic interaction with the negatively charged catalytic glutamic acid (E538) and cation-pi interaction with the aromatic tryptophan (W569). The stick representation of the molecular structure is coloured by heteroatoms (red:oxygen, blue:nitrogen) while hydrogen is not shown. The green sphere represents Mg2^+^ and the purple sphere represents Na^+^. Dashed lines represent hydrogen bonds.

To further validate the structure, we analysed soluble metabolites extracted from the captured LacZ-galactosyrin complex with gas chromatography-mass spectrometry (GC-MS) after chemical modifications to enable carbohydrate analysis. The peaks of the synthetic galactosyrin standard are identical to those detected in native galactosyrin and absent in the *`Δ`gsn*-derived sample (**Fig. S6A**). These mass spectra are consistent with the identified structure (**Fig. S6A, S9**). We also detected the same MS signals in apoplastic fluid extracted from *N. benthamiana* leaves infected with WT but not *`Δ`gsn* mutant *P. syringae* (**Fig. S6B**).

The LacZ-galactosyrin complex structure also revealed the inhibition mechanism. Galactosyrin binds to the enzyme active site and closely mimics the orientation of hydroxyl groups of galactose, the natural target of LacZ (**Fig. 2E**) (*10*). Remarkably, the branching geminal diol group allows the five-membered galactosyrin ring to mimic the conformation of the six-membered galactose ring. Additionally, the nitrogen of galactosyrin is likely protonated, resulting in a positive charge that electrostatically interacts with the catalytic glutamic acid (E538), and establishes a cation-pi interaction with the aromatic tryptophan (W569) (**Fig. 2E**). Indeed, the synthetic galactosyrin is a potent inhibitor of both LacZ from *E. coli* and BGAL1 from *N. benthamiana*, with an IC_50_ below that of 1-deoxy-galactonojirimycin and similar to galactostatin, two well-known iminosugars with 6-membered rings (**Fig. S7**).

### Galactosyrin is produced from a purine pathway intermediate through three enzymes and chemical conversion

To resolve the biosynthesis pathway of galactosyrin, we considered its structure and the putative functions of the three biosynthesis enzymes encoded by the *gsn* cluster. We first focused on GsnB as it is unique to the *gsn* cluster in our comparative genomics analysis (**Fig. S4**). GsnB is homologous to RibD reductase, which functions in the riboflavin synthesis pathway (**Fig. S8A**). The Alphafold2-predicted structure of GsnB contains conserved active site pockets similar to those in the crystal structures of RibD in complex with the substrate analog ribose-5-phosphate (R5P) and cofactor NADPH (*11*) (**Fig. S8B**), suggesting that GsnB could act on a similar substrate. This GsnB substrate likely also contains an amine group since the *gsn* cluster lacks an aminotransferase, unlike other clusters containing GsnA homologs (**Fig. S4**).

Considering that galactosyrin is a 5-carbon sugar-like molecule, we hypothesised that 5-phosphoribosyl-1-amine (PRA) might be a substrate of GsnB (**Fig. 3A, S8C**). PRA is produced by PurF from 5-phosphoribosyl-1-pyrophosphate (PRPP) and is used by PurD in purine synthesis (*12*). Indeed, when grown on purines to complement for purine deficiency, the *`Δ`purF* mutant is unable to produce galactosyrin, unlike the *`Δ`purD* mutant (**Fig. 3B**). The *`Δ`purD* mutant possibly produces even more galactosyrin than WT bacteria because this mutation prevents PRA conversion through PurD (**Fig. 3A, B**). These findings establish PRA as the precursor for galactosyrin biosynthesis.

**Fig. 3.**
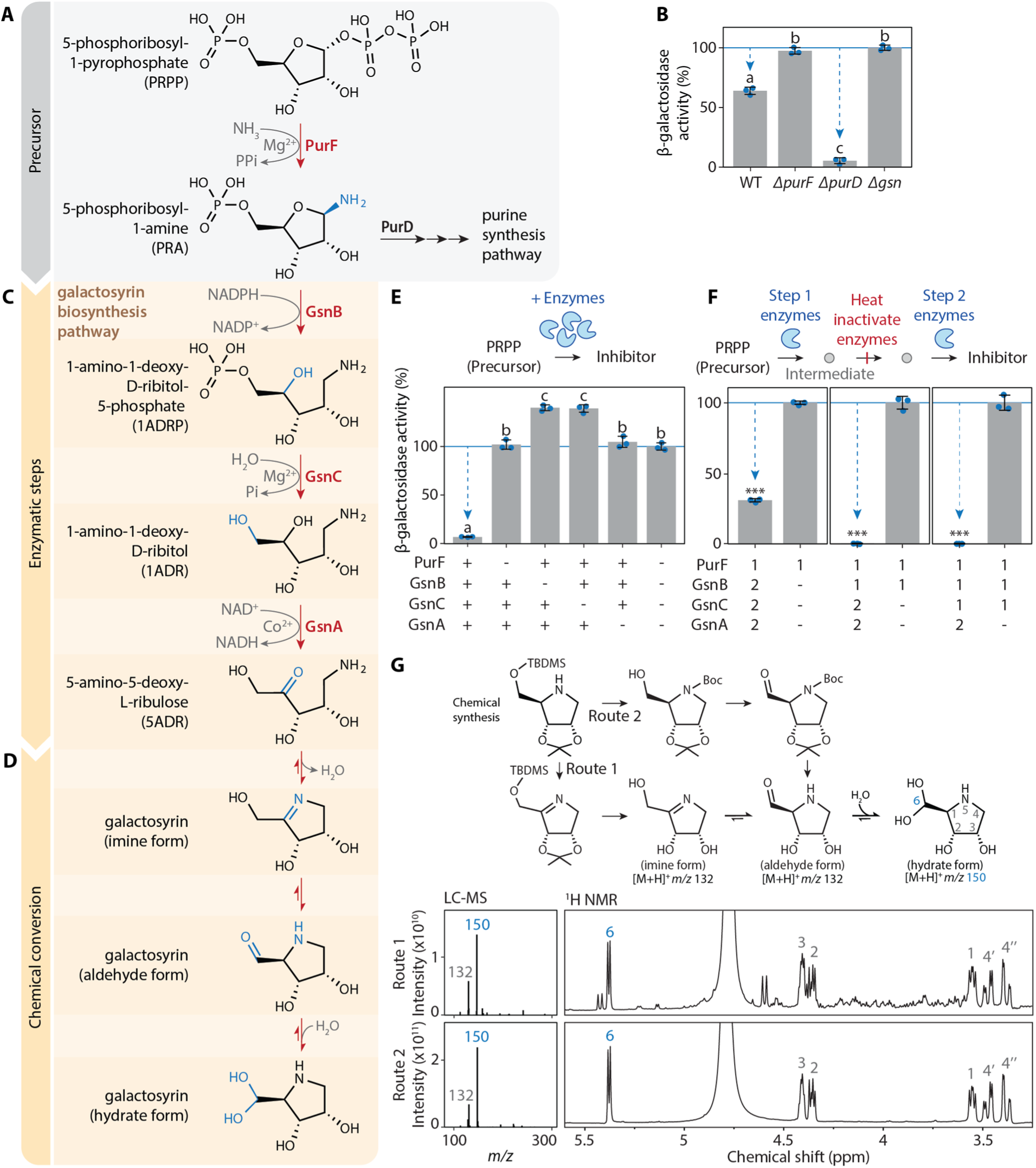
Galactosyrin biosynthesis branches off from purine biosynthesis by enzymatic and chemical conversion. **(A)** Galactosyrin biosynthesis branches off from the purine biosynthesis pathway. **(B)** *purF* but not *purD* is required for galactosyrin biosynthesis. Bacterial strains (WT or knockout mutants *`Δ`purF*, *`Δ`purD*, *`Δ`gsn*) were grown in virulence-inducing MG medium containing purines overnight then the supernatant was tested for inhibitor production. **(C)** Three *gsn*-encoded enzymes convert PRA into 5ADR. **(D)** A plausible 3-step chemical conversion pathway of 5ADR into the final hydrate form of galactosyrin. **(E)** PurF, GsnB, GsnC and GsnA are required and sufficient for the biosynthesis of galactosyrin from PRPP *in vitro*. Galactosyrin biosynthesis was reconstructed by mixing PRPP precursor with purified enzymes and their cofactors. For different mixtures, + indicates added enzymes, while - indicates omitted enzymes. **(F)** PurF, GsnB, GsnC and GsnA act consecutively in galactosyrin biosynthesis. Biosynthesis of galactosyrin was reconstructed by mixing PRPP precursor with purified enzymes and their cofactors in 2 separate steps: [1] enzymes added in the first step to produce an intermediate before heat inactivation of the enzymes, then [2] enzymes added in the second step to complete galactosyrin biosynthesis. **(B, E, F)** Inhibitor production was tested in an enzyme activity assay with FDG substrate and LacZ enzyme. β-galactosidase activity is reported as a percentage of the activity relative to the mean of no-inhibitor-control (*`Δ`gsn* for B, all enzymes omitted for E, enzyme 2 omitted for F). Arrows highlight inhibition. Error bars represent standard deviation from 3 replicates. Different letters indicate different groups with statistically significant difference (P < 0.001) using one-way ANOVA and post-hoc Tukey HSD test (for B, E). Asterisks indicate statistically significant difference (P < 0.001) using Welch’s t-test (for F). **(G)** Both imine and aldehyde forms of galactosyrin spontaneously convert into the hydrate form in water. (Top) Chemical synthesis of galactosyrin using two routes, via imine or aldehyde forms. (Bottom) Products were analysed with liquid chromatography-mass spectrometry (LC-MS) (left) and H^1^-NMR (right), showing the spectra that correspond to the spontaneously formed hydrate form. Positions within the structure of the hydrate form are numbered and labelled on the corresponding signals in NMR spectra.

Given the reductase activity of RibD, we speculated that GsnB could similarly reduce PRA into 1-amino-1-deoxy-D-ribitol-5-phosphate (1ADRP) using cofactor NADPH (**Fig. 3C**). Subsequently, the putative phosphatase GsnC might remove the phosphate of 1ADRP to produce 1-amino-1-deoxy-D-ribitol (1ADR). Finally, the putative oxidase GsnA might oxidise the secondary hydroxyl in 1ADR to produce the ketose 5-amino-5-deoxy-L-ribulose (5ADR). 5ADR can spontaneously convert into the detected hydrated galactosyrin (**Fig. 3D**), as explained below.

To confirm enzymatic steps of galactosyrin biosynthesis, we produced purified enzymes (PurF, GsnB, GsnC and GsnA) and incubated them with PRPP precursor and cofactors. Galactosyrin was produced from the mixture with all four enzymes, demonstrating that these components are sufficient to produce galactosyrin *in vitro* (**Fig. 3E**). Omission of any of the four enzymes blocked galactosyrin production (**Fig. 3E**), demonstrating that each enzyme is required for galactosyrin biosynthesis.

To verify the order of these reactions, we first produced intermediates from each enzymatic step *in vitro* and heat-inactivated the enzymes. We were then able to produce galactosyrin from these intermediates by adding the subsequent enzymes and cofactors in the expected order (**Fig. 3F**). Furthermore, the intermediates (PRA, 1ADRP and 1ADR) were detected after each step by GC-MS and these intermediates were depleted upon the addition of subsequent enzymes (**Fig. S10**). To confirm the final enzymatic step, we also detected galactosyrin formation from synthetic 1ADR by GsnA (**Fig. S11, S12B**), an NAD^+^-dependent oxidase (**Fig. S12C**). Notably, the pink color of purified GsnA indicated that cobalt ion is a preferred cofactor, confirmed *in vitro* (**Fig. S12D**), unlike other alcohol dehydrogenases of Pfam family PF00107, which are zinc-dependent (*13*). Taken together, these results confirmed the biosynthesis pathway of galactosyrin (**Fig. 3A**).

Finally, we propose a plausible chemical conversion pathway for the final steps of galactosyrin formation (**Fig. 3D**). First, imine formation by the amine and ketone in 5ADR produced the imine form of galactosyrin. This imine substructure is known to be unstable in aqueous environment and we propose that it will undergo spontaneous chemical conversions, including Heyn’s rearrangement (*14*) that results in the aldehyde form of galactosyrin. This aldehyde can then be hydrated to yield the hydrate form (**Fig. 3D**). To confirm that this pathway occurs spontaneously, we chemically synthesised both the imine and aldehyde derivatives of galactosyrin (**Fig. S13**) and found that, upon protecting group cleavage in water, they both spontaneously convert into the hydrate form detectable by both LC-MS and NMR as the major product (**Fig. 3G, S14**). In addition, GC-MS analysis of galactosyrin produced *in vitro* by GsnA from 1ADR detected both imine and aldehyde forms of galactosyrin because GC-MS was performed in anhydrous conditions (**Fig. S11**). These results demonstrate that 5ADR undergoes spontaneous chemical conversions that ultimately yield the hydrate form of galactosyrin discovered in LacZ by Cryo-EM.

### Galactosyrin triggers the accumulation of extracellular glycoproteins and glycosides

The originally identified target of galactosyrin is BGAL1, which acts in plant defence by facilitating the release of immunogenic peptides from glycosylated flagellin protein (*2*). We recently found that BGAL1 also removes terminal β-D-galactose from *N-* and *O-*glycans of transiently expressed recombinant proteins (*15*). Here, we found that BGAL1 can also process endogenous plant glycoproteins using RCAI, a terminal β-D-galactose-specific lectin, to probe the apoplastic proteome (**Fig. 4A**). Consequently, these RCAI-positive glycoproteins accumulated upon infection with WT *P. syringae* but not the *Δgsn* mutant in WT plants, while no differential accumulation of glycoproteins was observed in *bgal1-1* mutant plants (**Fig. 4A**), indicating that this modification of the host glycoproteome occurs through BGAL1 inhibition by galactosyrin. Moreover, untargeted metabolomics of apoplastic fluid from plants infected with WT and *Δgsn P. syringae* revealed that galactosyrin also triggers the accumulation of 873 ± 150 µM galactosylglycerol and 46 ± 12 µM trehalose in the apoplast of WT *P. syringae*-infected leaves (**Fig. 4B, S15**). The accumulation of these metabolites was independent of BGAL1 (**Fig. S15A**), suggesting that galactosyrin also targets other glycosidases that are involved in glycoside processing. Indeed, galactosyrin can inhibit several other glycosidases, including other β-galactosidases in *N. benthamiana* (**Fig. S16**) and α- and β-glucosidases (**Fig. S17**). Taken together, these findings demonstrate that galactosyrin is a multifunctional novel iminosugar produced by *P. syringae* to manipulate different plant glycosidases to influence various aspects of glycobiology in the plant apoplast during infection (**Fig. 4C**).

**Fig. 4.**
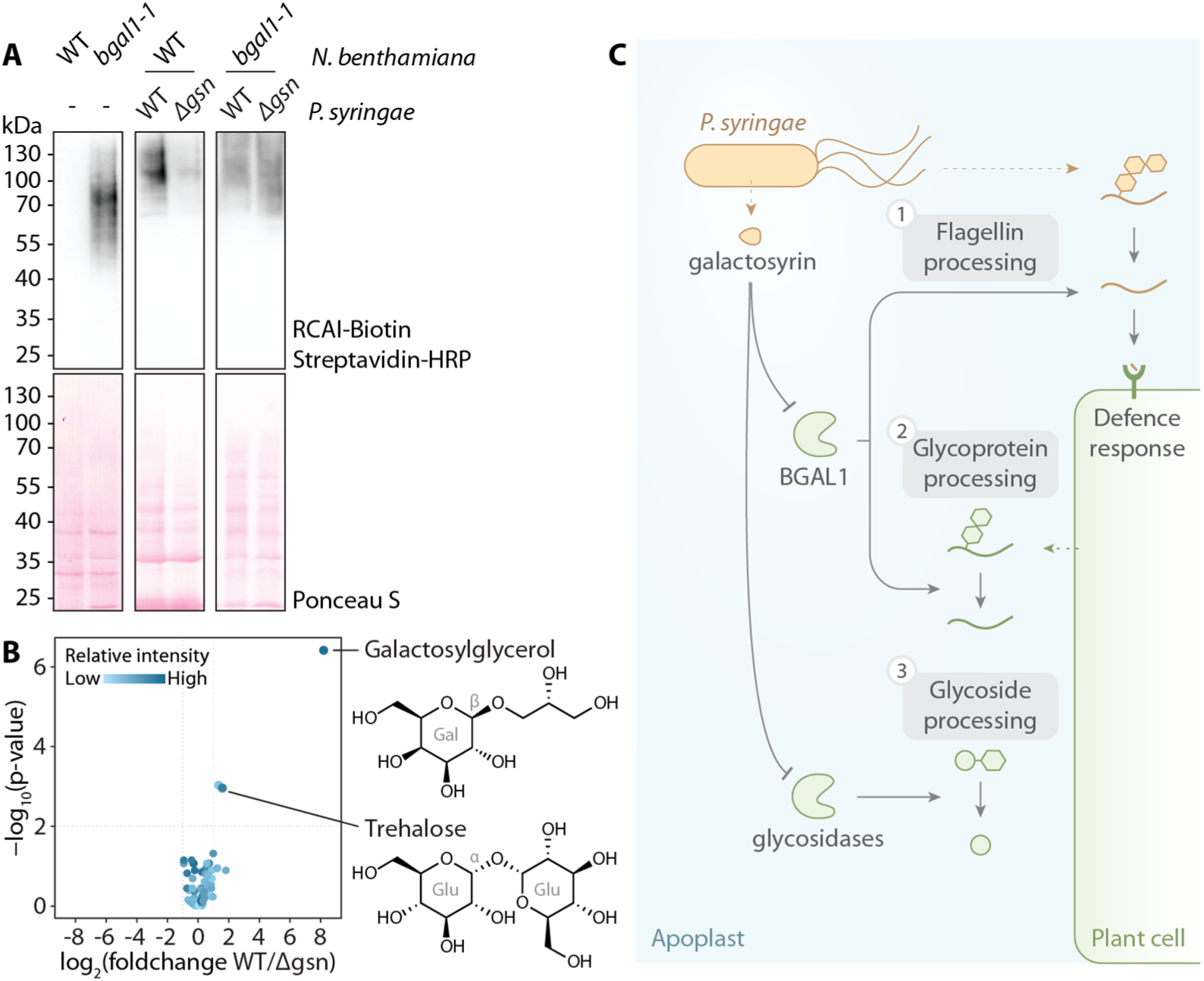
Galactosyrin manipulates multiple aspects of plant apoplast glycobiology. **(A)** Accumulation of RCAI-positive glycoproteins in the apoplast upon infection is dependent on BGAL1 and galactosyrin production. Proteins were extracted by acetone precipitation of apoplastic fluids from *N. benthamiana* (wild-type (WT) or BGAL1 knockout mutant (*bgal1-1*) with or without infection by *P. syringae* (WT or *`Δ`gsn*), then separated on SDS-PAGE, blotted and probed with RCAI lectin targeting galactose. **(B)** Galactosylglycerol and trehalose accumulate in the apoplast during infection dependent on galactosyrin production. Volcano plot of soluble metabolites detected by GC-MS of apoplastic fluids from leaves infected with WT or *`Δ`gsn* mutant (**Table S2**). Glucoside components are shown with galactose (Gal), Glucose (Glu) and bond configuration (α or β). **(C)** *P.* syringae produces galactosyrin to manipulate multiple aspects of glycobiology inside host plants by inhibiting plant glycosidases in the apoplast. [1] One major target of galactosyrin is the β-galactosidase BGAL1 previously shown to play a role in the processing of glycosylated flagellin to release plant defence elicitor (*2*). [2] Inhibition of BGAL1 by galactosyrin also interrupts apoplastic glycoprotein processing resulting in the accumulation of galactose-containing glycoproteins. [3] Galactosyrin also disrupts processing of glycosides by glycosidases other than BGAL1, resulting in the accumulation of galactosylglycerol and trehalose in the apoplast.

## DISCUSSION

We have elucidated the structure and biosynthesis pathway of galactosyrin, a novel iminosugar secreted by the model plant pathogen *P. syringae.* We discovered galactosyrin structure and its mode of action by solving the cryo-EM structure of inhibitor-bound β-galactosidase complex at an unprecedented atomic resolution (**Fig. 2**). Besides inhibiting BGAL1 to avoid the release of immunogenic flagellin fragments, we discovered that galactosyrin also manipulates the glycoproteome and metabolome in the apoplast of infected plants.

The biosynthesis pathway of galactosyrin is unique among iminosugars because it does not involve an aminotransferase but instead coopts an intermediate from the purine biosynthesis pathway using the NADPH-dependent reductase GsnB. Biosynthesis of 1-deoxynojirimycin (DNJ), nectrisine and 1,4-dideoxy-1,4-imino-arabinitol (DAB-1) all start with an aminotransferase acting on a sugar-phosphate precursor (*8*, *9*, *16*, *17*). However, all known iminosugar biosynthesis pathways involve an oxidase (GsnA homolog) to convert a hydroxyl group into a carbonyl group that then reacts with an amine group to form the iminosugar ring. Comparative genomic analysis of GsnA (**Fig. S4**) identified several biosynthesis gene clusters in bacterial species that are likely to produce novel iminosugars that remain to be characterized. Although hundreds of iminosugars isolated as natural products and thousands of synthetic analogues have been studied (*7*), galactosyrin is the first iminosugar with an aldehyde substituent attached to the heterocyclic ring. The hydrated form of the aldehyde is stable and constitutes a new class of iminosugars. We propose that this unique structure of galactosyrin is formed from an imine bond, which is then rearranged into an aldehyde form, followed by its hydration, thereby forming a branching geminal diol group, which is uniquely stable, even in water (**Fig. 3**). This property allows galactosyrin to accomodate a configuration that efficiently mimics galactose when bound to the active site of β-galactosidase. This discovery expands the diversity of iminosugars and initiates the exploration of a new class of galactosyrin-like iminosugars with different specificity or affinity. These new iminosugars might include important future pharmaceuticals because of their affinity to a wide range of carbohydrate active enzymes. Miglitol and DGJ, for instance, are iminosugars used to treat type-II diabetes and Fabry disease, respectively (*18*, *19*).

Galactosyrin is a multifunctional virulence factor produced by *P. syringae* during infection. First, galactosyrin inhibits BGAL1, which was previously shown to function in plant immunity by promoting the hydrolytic release of immunogenic fragments from glycosylated bacterial flagellin (*2*). Second, we discovered that the inhibition of BGAL1 also results in an accumulation of galactose-containing glycoproteins during infection with *P. syringae.* Alterations of glycoproteomes during infection by *P. syringae* have also been reported (*20*, *21*). We previously found that BGAL1 removes the terminal β-D-galactose residues of both *N*- and *O*-glycans on recombinant glycoproteins (*15*). Galactose is also a common monosaccharide in endogenous glycans of *N*- and *O*-glycosylated proteins such as cell surface receptor kinases and defence-related arabinogalactan proteins (AGPs) (*22*). At this stage, it is unclear how the manipulated glycoproteome could affect the bacterial colonization but the loss of apoplastic β-galactosidase activity was also reported to impact cell wall glycan processing and functions, such as interactions and hydration properties (*23–25*).

We also discovered that galactosyrin induces an accumulation of galactosylglycerol and trehalose in the apoplast of infected plants, which is independent from BGAL1 and possibly results from galactosyrin inhibiting other glycosidases. Degradation of glycosides by glycosidases could therefore be another mechanism to modulate solute levels in the apoplast. Although pathogens often induce accumulation of sugars and metabolites in the apoplast through production or efflux to provide nutrient sources (*26*, *27*), trehalose and galactosylglycerol are not consumed by *P. syringae in vitro* (**Fig. S15D**). On the other hand, these glycosides are well-known osmolytes (*28*, *29*) and may therefore contribute to the establishment of aqueous apoplast conditions that promotes virulence (*30*, *31*). Accumulation of glycosides might also influence bacterial colonisation in different ways. Elevated trehalose levels, for instance, dampens plant defence responses and promotes *P. syringae* infection (*32*). In addition, by inhibiting β-glucosidases, galactosyrin may also prevent the activation of glucosides that act in plant defence and signaling (*33*).

The presence of homologous iminosugar biosynthesis gene clusters in plant pathogens such as *Acidovorax* and *Erwinia*, and plant-associated bacteria such as *Kosakonia*, *Bacillus* and *Paenibacillus* indicates that the use of iminosugars to manipulate the glycobiology of the host plant might be a common strategy used by plant-associated bacterial pathogens and symbionts.

## ACKNOWLEDGEMENTS

We thank Pedro Bota for GC-MS maintenance and training; Urszula Pyzio for plant care; Sarah Rodgers, Caroline O’Brien and Patricia Bowman for technical support; Brian Mooney and Jie Huang for feedback on the manuscript. We thank Diamond Light source for access and support of the cryoEM facilities at the UK National Electron Bio-Imaging Centre (eBIC), proposal NT21004, NT29812, and BI28713.

Computation was performed at the Diamond Light Source and the Oxford Biomedical Research Computing (BMRC) facility, a joint development between the Wellcome Centre for Human Genetics and the Big Data Institute (BDI) supported by Health Data Research UK and the NIHR Oxford Biomedical Research Centre.

## Author contributions

Conceptualization: NS, BC, GMP, RALvdH

Methodology: NS, BC, DK, GMP, RALvdH, YS, NH, PZ, WWAT, MK, MD, PF, SY, AK

Investigation: NS, BC, DK, PB, YS, NH, WWAT, MD, SY, AK, RN, GF

Funding acquisition: AK, PF, MK, PZ, GMP, RALvdH

Supervision: AK, PF, MK, PZ, GMP, RALvdH

Writing – original draft: NS, RALvdH

Writing – review & editing: all authors

## Data and materials availability

All data are available in the manuscript, supplementary materials and cited references. The cryoEM density maps and corresponding atomic models have been deposited in the EMDB and PDB, respectively. The accession codes are: for LacZ with native inhibitor (WT), EMDB-19182 and PDB 8RI7; for LacZ with *Δgsn* negative control, EMDB-19181 and PDB 8RI6; for LacZ with synthetic galactosyrin EMDB-19183 and PDB 8RI8.

## Materials and methods

### Bacterial strains, culture conditions and growth media

Bacterial strains and plasmids used in this study are listed in **Table S3** and **Table S4**, respectively. Generally, *P. syringae* and *A. tumefaciens* were grown at 28°C and *E. coli* were grown at 37°C. Liquid cultures were incubated with shaking at 220 rpm. Growth media were made as follows. LB medium: 10g/L tryptone, 5 g/L yeast extract, 10 g/L NaCl. Mannitol glutamate (MG) virulence-inducing minimal medium (*38*): 10 g/L mannitol, 2 g/L L-glutamic acid, 0.5 g/L KH_2_PO_4_, 0.2 g/L NaCl, 0.2 g/L MgSO_4_.7H_2_O, 50 µM iron citrate, adjusted to pH 5.5. M9 fructose medium: 6 g/L Na_2_HPO_4_, 3 g/L KH_2_PO_4_, 0.5 g/L NaCl, 1 g/L NH_4_Cl, 0.1 mM CaCl_2_, 2 mM MgSO_4_, 22 mM fructose, adjusted to pH 5.5. Solid medium for plating contains 1.5% (w/v) agar. Antibiotics were generally used at the concentrations of 25 µg/ml rifampicin, 12.5 µg/ml gentamicin, 50 µg/ml kanamycin, 10 µg/ml tetracycline, 100 µg/ml carbenicillin, 25 µg/ml chloramphenicol. For purine auxotrophs (Δ*purF* and Δ*purD* mutants), cultures were supplemented with 0.2 mM adenine, guanine and thiamine.

### Bacterial conjugation

Overnight bacterial cultures in LB medium were collected by centrifuging at 4,000 × g for 5 minutes, washed and resuspended in LB medium with no antibiotic. The recipient strain (*P. syringae* to be transformed), donor strain (*E. coli* containing plasmid constructs) and helper strain (*E. coli* pRK2013) were mixed in a ratio of 7:3:1 and centrifuged at 4,000 × g for 5 minutes to form a pellet which was spotted on to LB agar without antibiotics and incubated at 28°C overnight. The bacterial mixture was then resuspended and plated on LB agar with 25 µg/ml nitrofurantoin to select against *E. coli* and appropriate antibiotics to select for the desired transformants.

### Forward genetic screen for galactosyrin mutants

A random transposon insertion mutant library was created by delivering mini-Tn5 transposon into the recipient *P. syringae* strain carrying a plasmid with constitutively expressed *lacZ* β-galactosidase gene (WT *lacZ*) through triparental mating conjugation with the *E. coli* donor strain (mini-Tn5 transposon) and helper strain (pRK2013). The resulting mutant library was plated on MG agar supplemented with 12.5 µg/ml each of rifampicin, gentamicin and kanamycin, 0.1 mg/ml X-gal (5-bromo-4-chloro-3-indolyl-beta-D-galacto-pyranoside) and 0.1 mM IPTG. Candidate galactosyrin inhibitor mutant colonies were identified by their darker blue colour from the higher activity of LacZ cleaving X-gal in the absence of galactosyrin. To validate the mutants, these candidate colonies were grown in MG medium overnight and the supernatant was heat inactivated at 95°C for 5 minutes and used in FDG assay to confirm the lack of inhibitor production. Then, inverse PCR was performed to amplify the transposon insertion site. Briefly, a small amount of bacteria was resuspended in water and heated at 95°C for 5 minutes to release DNA, which was restriction digested with Fast Digest PaeI (Thermo Scientific) at 37°C for 1 hour, heat inactivated at 65°C for 20 minutes, then self-ligated by adding ATP and T4 DNA ligase (Thermo Scientific) at 16°C for 18 hour and heat inactivated at 65°C for 20 minutes. The ligated DNA was used in a PCR with Q5 high-fidelity DNA polymerase (New England Biolabs) with GC enhancer using primer pair oNS22/oNS23 (**Table S5**). PCR products were treated with Illustra ExoProStar (Cytiva) and Sanger sequenced (Eurofins, Luxembourg) using primer oNS15 (**Table S5**). To identify transposon insertion sites in the genome, the sequence adjacent to the transposon boundary was BLAST searched against the *P. syringae* pv. *tomato* DC3000 genome using The Pseudomonas Genome Database (pseudomonas.com) (*39*).

### Galactosyrin production from bacterial culture

Bacterial cells were collected from an overnight culture in LB medium by centrifugation at 4,000 × g for 5 minutes, washed with sterile water, resuspended in MG medium to an OD_600_ of 0.5 and grown overnight at 28 °C (for both *P. syringae* and *E. coli*). Then the culture was centrifuged again and the supernatant containing the inhibitor was collected.

### FDG assay for β-galactosidase activity and inhibition

For LacZ assays, the reaction was set up in Z buffer (60 mM Na_2_HPO_4_, 40 mM NaH_2_PO_4_, 10 mM KCl, 1 mM MgSO_4_) with 0.2 µM FDG (Fluorescein di-β-D-galactopyranoside, Marker Gene Technologies, M0250) and 0.1 mU/µl LacZ (β-galactosidase enzyme from *E. coli*, Sigma, G6008). For inhibition tests, inhibitor samples made up half the reaction volume. For apoplastic fluid assays, apoplastic fluid from *N. benthamiana* was incubated with 0.2 µM FDG and 50 mM MES pH 5.5. Fluorescence signal from fluorescein, a product of FDG cleavage by β-galactosidase, was measured with 485 nm excitation and 535 nm emission every minute at 25°C using the plate reader Infinite M200 (Tecan). β-galactosidase activity was calculated from the rate of fluorescence increase over time. For *in planta* assays, 10 µM FDG was infiltrated into *N. benthamiana* leaves as spots then the leaves were imaged using Amersham Typhoon 5 (Cytiva) with 488 nm excitation laser and Cy2 emission filter (525BP20) at 300 PMT. Images were analysed using Fiji (*40*). β-galactosidase activity was calculated from the integrated density of fluorescence signal from each spot subtracted by a background from an uninfiltrated spot on each leaf.

### Generation of gene deletion mutants

Deletion of the gene of interest in *P. syringae* genome was generated by the two-step allelic exchange method (*41*). For plasmid construct generation, fragments of around 1 kb flanking each side of the target gene were PCR amplified from genomic DNA using Q5 high fidelity DNA polymerase (New England Biolabs) with primer pairs L_F/L_R (left flanking fragment) and R_F/R_R (right flanking fragment) (**Table S5**). The plasmid backbone was PCR amplified from the suicide vector pK18mobsacB (*42*) using Q5 high fidelity DNA polymerase (New England Biolabs) with primer pair oNS34/oNS35 (**Table S5**) then assembled with the flanking fragments by Gibson assembly and transformed into *E. coli* TOP10. The resulting constructs were checked for correct assembly by Sanger sequencing using primers oNS50 and oNS51 (**Table S5**). The validated plasmids were transformed into *P. syringae* by triparental mating conjugation with the helper strain (pRK2013). The resulting merodiploid transformants were counter-selected on LB agar plates containing 10% (w/v) sucrose. Then, colonies were screened by colony PCR using primer pairs L_F/R_R (**Table S5**) to confirm the deletion mutant with the expected size of the deleted genomic region.

### Generation of *gsn* cluster expression constructs

The sequence of the *gsn* cluster was PCR amplified from *P. syringae* WT genomic DNA using Q5 high fidelity DNA polymerase (New England Biolabs) with primer pairs oNS68/oNS63 for the *gsn* cluster with native promoter, or oNS206/oNS63 for the *gsn* cluster with no promoter (**Table S5**). The pBBR1MCS plasmid backbones (*43*) were linearised by restriction digest with Fast Digest SmaI and NsiI (Thermo Scientific) for pBBR1MCS2 (*l*ac promoter removed for insertion of *gsn* cluster with native promoter), or SmaI for pBBR1MCS3 (for insertion of *gsn* cluster with no promoter downstream of *lac* promoter) then assembled with the gene fragment by Gibson assembly and transformed into *E. coli* TOP10. The resulting constructs were checked for correct assembly by Sanger sequencing using primers seq_F and seq_R (**Table S5**). The validated plasmids were transformed into *P. syringae* by triparental mating conjugation with the helper strain (pRK2013).

### Reverse transcriptase polymerase chain reaction (RT-PCR)

*P. syringae* cells were collected from LB agar plates grown at 28°C for 2 days, washed with sterile water, resuspended at an OD600 of 1 in MG medium and incubated at 28°C with shaking for 6 hours to induce virulence gene expression. Then, the cells were collected by centrifuging at 12,000 × g for 5 min, frozen with liquid nitrogen and lysed with lysozyme (5 mg/ml, Sigma) in TE buffer (10 mM Tris-HCL pH 7.4, 1 mM EDTA) at 25°C for 10 minutes while shaking. Total RNA was extracted from the sample using Monarch Total RNA miniprep kit (New England Biolabs), treated with Turbo DNA-free kit (Invitrogen) to remove genomic DNA followed by cDNA synthesis with GoScript reverse transcriptase master mix with random primers (Promega). PCR was then performed using GoTaq green master mix (Promega) with primer pairs F/R listed in **Table S5**. The PCR product was detected upon separation in 4% (w/v) agarose gel electrophoresis and staining with ethidium bromide. The no RT controls were performed on equivalent samples without reverse transcriptase to ensure that no amplification was detected from genomic DNA contamination.

### Generation of Tn7 lux reporter constructs

The Tn7 lux reporter construct (pNS161) was derived from pRS-pOXB20:lux (*44*) by removing the OXB20 promoter by restriction digest with Ecl136II and Eco105I (Thermo Scientific) and replacing with T0T1 terminators amplified from pRS-pOXB20:lux using primer pair oNS386/oNS387 (**Table S5**). To generate the promoter:lux reporter fusion, the sequence of the promoter of interest was PCR amplified from *P. syringae* WT genomic DNA using Q5 high fidelity DNA polymerase (New England Biolabs) with primer pairs F/R (**Table S5**) and assembled into the Ecl136II restriction site in pNS161 by Gibson assembly and transformed into *E. coli* TOP10. The resulting constructs were checked for correct assembly by Sanger sequencing using primers oNS384 and oNS385 (**Table S5**). The validated plasmids were used in a four-parental mating conjugation with helper strains (pRK2013 and pUXBF13) to integrate the Tn7 transposon construct into the genome of *P. syringae*.

### Luminescence reporter activity in planta

*P. syringae* strains with Tn7 lux reporter constructs were collected from an overnight culture in LB medium by centrifuging at 4,000 × g for 5 minutes, washed with 10 mM MgCl_2_ and resuspended in 10 mM MgCl_2_ at the OD_600_ of 1 (for imaging the same day) or 0.001 (for imaging after 3 days). Bacterial suspensions were infiltrated into *N. benthamiana* leaves using a needleless syringe. Then, leaves were detached from the plant, placed in a plastic Petri dish with wet tissue paper, left in the dark for 5 minutes and imaged for luminescence signal using Imagequant LAS-4000 imager (Cytiva).

### Infection assay

*P. syringae* cells were collected from an overnight culture in LB medium by centrifuging at 4,000 × g for 5 minutes, washed with sterile water and resuspended in sterile water at the OD_600_ of 0.2 (∼10^8^ CFU/ml) with 0.04% (v/v) Silwet L-77. The bacterial suspension was then sprayed on both surfaces of 4-5 weeks-old *N. benthamiana* plant leaves until runoff. The infected plants were placed in a transparent plastic box to keep high humidity and maintained in a growth chamber at 21°C with 12-hour photoperiod and light intensity of 100 μmol/m^2^/s. At 3 days post infection, leaf discs were then collected from infected tissue, surface sterilised with 70% (v/v) ethanol, washed with sterile water and homogenised in 1 ml of sterile water for serial dilution plating. Colonies were counted to calculate the number of bacterial colony forming units (CFU) per cm^2^ area of infected tissue. The bacterial suspension used for inoculation was also plated to confirm the bacterial inoculum.

### Bacterial growth curve *in vitro*

*P. syringae* cells were collected from an overnight culture in LB medium by centrifuging at 4,000 × g for 5 minutes, washed with sterile water and resuspended in sterile water at the OD_600_ of 0.05 in 150 µl of media in each well of 96 well plate. Bacterial growth was monitored with optical density at 600 nm (OD600) taken every 30 minutes using the plate reader Infinite M200 (Tecan) maintained at 28°C with shaking.

### Generation of recombinant protein production constructs

The sequence of the gene of interest was PCR amplified from *P. syringae* WT genomic DNA using Q5 high fidelity DNA polymerase (New England Biolabs) with primer pairs F/R (**Table S5**). The pET-28b plasmid backbone was PCR amplified using Q5 high fidelity DNA polymerase (New England Biolabs) with primer pair oNS207/oNS208 (**Table S5**) then assembled with the gene fragments by Gibson assembly and transformed into *E. coli* TOP10. The resulting constructs were checked for correct assembly by Sanger sequencing using primers oNS209 and oNS210 (**Table S5**). The validated plasmids were then transformed into *E. coli* expression strains.

### Enzyme purification

*E. coli* expression strains were grown in LB medium with appropriate antibiotics at 37°C overnight, then diluted in a fresh LB medium to OD600 of 0.04 and grown until OD600 of 0.6, then IPTG was added (0.4 mM) to induce protein expression at 20°C overnight. For GsnB, 500 uM L-rhamnose was also added. Then, bacterial cells were collected by centrifugation and lysed with CelLytic Express (Sigma). Strep-tagged enzymes from the lysate were isolated with Strep-Tactin XT 4Flow resin (IBA Life Sciences), washed with TBS buffer (50 mM Tris, 150 mM NaCl, pH 7.5) and eluted with TBS buffer with 50 mM biotin. Protein concentration was measured using Bradford assay (Sigma) with BSA (bovine serum albumin) standards.

### Galactosyrin biosynthesis pathway reconstruction *in vitro*

The reaction was set up in TBS buffer (50 mM Tris, 150 mM NaCl, pH 7.5) with 100 mM NH_4_Cl, 1 mM MgCl_2_, 1 mM PRPP (5-Phospho-D-ribose 1-diphosphate), 1 mM NAD^+^ (nicotinamide adenine dinucleotide), 1 mM NADPH (nicotinamide adenine dinucleotide phosphate) and biosynthesis enzymes (0.2 µM PurF, 0.2 µM GsnB, 0.4 µM GsnC and 2.6 µM GsnA). For each enzyme omission, an equivalent amount of GFP was used instead. The reaction mixture was incubated at 25°C for 20 hours to produce galactosyrin. For stepwise reaction, the first reaction was heat inactivated at 95°C for 5 minutes before the components of the second reaction was added.

### GsnA assay

The reaction was set up in TBS buffer (50 mM Tris, 150 mM NaCl, pH 7.5) with 1 mM NAD^+^, 1 mM 1ADR (1-amino-1-deoxy-ribitol) and 10 µM GsnA. Enzyme activity was monitored by measuring the formation of NADH with 340 nm absorbance every minute at 25°C using the plate reader Infinite M200 (Tecan). For metal cofactor preference test, GsnA was pretreated with 100 µM EDTA to chelate divalent metal ions. Then, the assay was supplemented with or without 1 mM CoCl_2_ or ZnCl_2_.

### Isolation of 1ADR

Bacterial culture supernatant prepared the same way as galactosyrin production was mixed with 50% volume methanol and 20% volume chloroform, centrifuged at 10,000 × g for 5 minutes, then the top aqueous fraction was collected. 1ADR from the fraction was isolated with Amberlyst 15 hydrogen form cation exchange resin (Supelco), washed with 70% (v/v) ethanol followed by water, and eluted with 1M NH_4_OH. The eluate was freeze dried then resuspended in water.

### Protein structure modeling

GsnB (PSPTO_0835) structure was predicted with Alphafold2 through ColabFold (*63*) with default settings. The first ranked model was used. ChimeraX was used to visualise protein structures and matchmaker was used for structure comparison (*64*).

### Inhibitor dose response curve

β-galactosidase activity was measured with the FDG assay as described above in the presence of different concentrations of each inhibitor. The activity was normalised with a no inhibitor control set as 100%. Then, drc package (*65*) was used to fit a four-parameter logistic model with a lower limit of 0 and upper limit of 100 to calculate the half maximal inhibitory concentration (IC_50_).

### N. benthamiana infection

*P. syringae* cells were collected from an overnight culture in LB medium by centrifuging at 4,000 × g for 5 minutes, washed and resuspended in sterile water at the OD_600_ of 0.0002 (∼10^5^ CFU/ml). The bacterial suspension was then infiltrated into leaves of 4-5 weeks-old *N. benthamiana* plants. Infected plants were maintained in a growth chamber at 21°C with 12-hour photoperiod and light intensity of 100 μmol/m^2^/s. Samples were collected after 3 days for GC-MS metabolomics or 5 days for lectin blot.

### Apoplastic fluid extraction

*N. benthamiana* leaves were rolled into a 50 ml syringe filled with ice-cold water. The leaves were infiltrated by pulling the plunger to apply negative pressure drawing air out of the leaf tissue then pushing the plunger to apply positive pressure forcing water into the leaf tissue, while keeping the syringe tip sealed. The infiltrated leaves were blotted dry and rolled into a 20 ml syringe barrel placed inside a 50 ml conical centrifuge tube. Apoplastic fluid was collected from the leaves by centrifugation at 900 × g for 15 minutes at 4°C.

### Lectin blot

Proteins were collected by mixing apoplastic fluids with 4 volumes of acetone and 20 mM NaCl, incubating at room temperature for 15 minutes, then centrifuging at 12,000 × g for 10 minutes. The protein pellet was briefly air dried and resuspended in gel loading buffer (100 mM Tris-Cl (pH 6.8), 200 mM DTT, 4% (w/v) SDS, 0.2% (w/v) bromophenol blue, 20% (v/v) glycerol) then heated at 95°C for 5 minutes before being separated in 12% (w/v) polyacrylamide gel electrophoresis and transferred onto PVDF membrane using Trans-Blot Turbo system (Bio-Rad). The membrane was blocked with Carbo-Free blocking solution (Vector Laboratories) with 0.05% (v/v) Tween-20, then incubated with 5 µg/ml biotinylated RCA I lectin (Ricinus Communis Agglutinin I, RCA120, Vector Laboratories) in PBS buffer (10 mM sodium phosphate buffer, 3 mM KCl, 140 mM NaCl, pH 7.4) with 0.05% (v/v) Tween-20 (PBST), then washed with PBST, then incubated with Streptavidin-HRP (Sigma) (1:5,000x dilution in 5% BSA in PBST), and then washed with PBST. Chemiluminescence was detected using Clarity ECL western blotting substrates (Bio-Rad) with an Imagequant LAS-4000 imager (Cytiva). The membrane was then stained with Ponceau S for total protein detection.

### Untargeted metabolomics

Apoplastic fluid samples (200 µl) were spiked with 20 µl of 2 µg/ml ribitol as an internal standard and analysed with GC-MS as described in GC-MS section. GC-MS data were processed using MassHunter Unknown Analysis (Agilent). Metabolite peaks were detected using TIC analysis with default settings, then mass spectra were searched against NIST11 (*66*) and Golm (*67*) library to putatively assign compound identifications to hits with match factors of at least 90. Peaks of the same metabolite feature were aligned using hierarchical clustering with a retention time window of 0.06. Peaks also detected in solvent blank samples were excluded. Metabolite features not robustly detected in all samples were removed. Peak area of each metabolite feature was log2 transformed and normalised with the peak of ribitol internal standard in each sample. Differentially accumulating metabolites were identified using Welch’s t-test with p-values adjusted using Benjamini-Hochberg method to correct for multiple comparisons.

### Estimation of apoplastic metabolite concentrations

Apoplastic fluid samples and metabolite standards at different concentrations (50 µl) were spiked with 5 µl of 0.1 mg/ml ribitol as an internal standard and analysed with GC-MS as described in GC-MS section. The peak area of each metabolite was normalised with the peak area of ribitol in each sample. A standard curve was constructed from the normalized peak areas and the corresponding concentrations of standards then linear regression was used to calculate the unknown concentration of the metabolite in apoplastic fluid samples based on their normalized peak area. Since apoplastic fluids extracted from leaves were diluted compared to the original plant apoplast in the leaf tissue, metabolite concentrations in the plant apoplast was estimated by multiplying the concentration in the apoplastic fluid samples by a dilution factor of 2.7 ± 0.3, determined based on the method from (*68*).

### Glycosidase inhibition assay

The enzymes β-glucosidase (from almond and bovine liver), α-galactosidase (from coffee beans), β-galactosidase (from *E. coli* and bovine liver), β-glucuronidases (from *E.coli* and bovine liver), β-xylosidase (*T. longibrachiatum*), *p*-nitrophenyl glycosides, and various disaccharides were purchased from Sigma-Aldrich Co. Brush border membranes were prepared from the rat small intestine according to the method of Kessler et al. (*69*) and were assayed at pH 6.8 for rat intestinal maltase using maltose. The inhibitory activity toward human lysosomal acid glycosidases was measured with Myozyme, Cerezyme, and Fabrazyme (Genzyme) as the enzyme source and an appropriate 4-methylumbelliferyl-glycopyranoside (Sigma-Aldrich) as substrate. The reaction mixture consisted 100 mM McIlvaine buffer (pH 5.2), 0.25% sodium taurocholate and 0.1% Triton X-100 (Nacalai Tesque Inc), and the appropriate amount of enzyme. The reaction mixture was pre-incubated at 0°C for 45 min, and the reaction was started by the using 3 mM substrate solution, followed by incubation at 37°C for 30 min. The reaction was stopped by the addition of 1.6 mL of the solution of 400 mM Glycine-NaOH solution (pH 10.6). The released 4-methylumbelliferone was measured (excitation 362 nm, emission 450 nm) with a F-4500 fluorescence spectrophotometer (Hitachi). For rat intestinal maltase activity, the reaction mixture contained 25 mM maltose, and the appropriate amount of enzyme, and the incubations were performed for 10 min at 37 °C. The reaction was stopped by heating at 100 °C for 3 min. After centrifugation (600 g; 10 min), the resulting reaction mixture was added to the Glucose CII-test Wako (Wako Pure Chemical Ind.). The absorbance at 505 nm was measured to determine the amount of the released D-glucose. Other glycosidase activities were determined using an appropriate *p*-nitrophenyl glycoside as substrate at the optimum pH of each enzyme. The reaction mixture contained 2 mM of the substrate and the appropriate amount of enzyme. The reaction was stopped by the addition of 400 mM Na_2_CO_3_. The released *p*-nitrophenol was measured spectrometrically at 400 nm.

### Phylogenetic analysis

For each tree, the sequences described below were aligned using MAFFT 7 with L-INS-i algorithm (*45*). The resulting multiple sequence alignment was used to construct a maximum likelihood phylogenetic tree using IQ-TREE 2 (*46*) with the best-fit amino acid substitution model from ModelFinder (*47*). Branch support values were calculated using ultrafast bootstrap with 1,000 replications. Phylogenetic trees were visualised using iTOL (*48*).

### Phylogeny of *P. syringae*

The *P. syringae* tree was constructed using concatenated coding DNA sequences of 4 conserved housekeeping genes (*rpoD* encoding sigma factor 70, *gyrB* encoding DNA gyrase B, *gltA* encoding citrate synthase and *gapA* encoding glyceraldehyde-3-phosphate dehydrogenase), identified through a local BLAST+ search (*49*) against selected genomes obtained from publicly available NCBI (*50*) and pseudomonas.com databases (*39*). *P. syringae* genomes used in the analysis were selected to include all strains that have the *gsn* gene cluster identified from a BLAST search of GsnA amino acid sequence from *P. syringae* pv. *tomato* DC3000 (on www.blast.ncbi.nlm.nih.gov) with default settings, 62 type and pathotype strains (*51*) and additional strains to represent different phylogroups. Strains that have incomplete assembly or annotation at the site of the 4 housekeeping genes were discarded. *P. fluorescens* SBW25 was included as an outgroup. Phylogroups were assigned according to (*52*). For strains that contain the *gsn* cluster, the *gsn* cluster together with 20 kb of genomic region flanking each side was displayed. Gene model annotation of the genome was acquired from the RefSeq database (*53*) and gene family assignment was based on conserved domain database entry (CDD) linked to each gene (*54*). Many available genomes were not fully assembled so fragmented genomic regions and incompletely annotated genes are common.

### Phylogeny of gsnA

The phylogenetic tree was constructed using coding DNA sequences of homologs of *gsnA* identified in *P. syringae* strains from above. A homolog of *gsnA* from *P. fluorescens* Pf275 was included as an outgroup since *P. fluorescens* SBW25 does not contain the *gsn* cluster.

### Phylogeny of GsnA homologs

The amino acid sequence of GsnA from *P. syringae* pv. *tomato* DC3000 was used as a query for a PSI-BLAST search (on www.blast.ncbi.nlm.nih.gov) (55) on RefSeq database with default settings over 3 iterations to obtain 1,000 proteins, which were then filtered to include only hits with a length of 300-400 amino acids and at least 80% query coverage. For brevity, only a maximum of 5 homologs from each species were used as representatives. The sequence of threonine dehydrogenase from *E. coli* (WP_000646007.1) was included as an outgroup. Putative gene clusters were assigned by identifying a stretch of consecutive genes with the same orientation on the same strand of the genome as the gene encoding the GsnA homolog, both up and downstream until there was a gene with an opposite orientation. Gene model annotation of the genome was acquired from the RefSeq database (*53*) and gene family assignment was based on conserved domain database entry (CDD) linked to each gene (*54*). The association with transposable elements was defined as the presence of transposable elements within five genes distance from the first and last gene in the cluster. Many available genomes were not fully assembled so fragmented genomic regions and incompletely annotated genes are common.

## Capture of enzyme-inhibitor complex

### Galactosyrin production

*P. syringae* WT cells were collected from an overnight culture in LB medium by centrifugation at 4,000 × g for 5 minutes, washed and resuspended to the OD_600_ of 1 in M9 fructose medium to induce inhibitor production overnight at 28 °C. Then the culture was centrifuged again and the supernatant containing the inhibitor was collected, filtered through a 0.4 μm syringe filter and adjusted to pH 7.2 with NaOH. As a negative control, the equivalent sample from the galactosyrin-deficient Δ*gsn* mutant was also produced.

### LacZ production

The *E. coli* expression strain (pNS141) was grown in LB medium with 50 ug/ml kanamycin at 37°C overnight, then diluted in fresh LB medium to OD600 of 0.04 and grown until OD600 of 0.6, then IPTG was added (0.4 mM) to induce protein expression at 28°C overnight. The bacterial cells were collected by centrifugation and lysed with CelLytic Express (Sigma). The lysate was mixed with imidazole (10 mM) before adding HisPur Ni-NTA resin (Thermo Scientific) and incubating at 4°C for 30 minutes to immobilize the His-tagged LacZ protein on the resin, which was then washed with 2xPBS (20 mM sodium phosphate buffer, 6 mM KCl, 280 mM NaCl, pH 7.4) with 25 mM imidazole.

### Complex capture

The immobilized LacZ was loaded into a column then at least 10-fold volume of *P. syringae* supernatant from the first step was passed through the column to allow LacZ to capture galactosyrin to saturation. The column was then washed with 2xPBS with 25 mM imidazole before the complex was eluted with 1xPBS with 250 mM imidazole. For quality control, the eluted complex was run on SDS-PAGE to check the purity and quantity. To assess inhibitor saturation, the eluted complex was diluted 100 times and used in FDG assay to measure its enzyme activity compared to a negative control sample with no inhibitor.

### Metabolite extraction

LacZ-galactosyrin complex capture was performed as above except the elution step. Instead, the immobilized complex was washed with water then with 17 mM acetic acid to denature the enzyme and release the metabolite.

## Cryo-electron microscopy (cryo-EM)

### Sample preparation

Purified LacZ β-galactosidase samples were refined using the Akta Pure chromatography system (Cytiva) with SuperDex 200 size exclusion chromatography column (Cytiva) and elution buffer with 25 mM Tris, 50 mM NaCl, 2 mM MgCl_2_, 2 mM EDTA, 1 mM TCEP, pH 8. Protein concentration was measured with Nanodrop (Thermo Scientific) and adjusted to 0.7 mg/ml. Samples of native galactosyrin (WT) and negative control (Δ*gsn*) were prepared from enzyme-inhibitor complex capture. The sample with synthetic galactosyrin was prepared by adding 5 mM galactosyrin into a purified LacZ aliquot and incubating on ice for 30 minutes before applying onto grids.

### Loading

An aliquot of 3 μl of control sample (Δ*gsn*) was applied onto a glow-discharged holey carbon copper grid (300 mesh, QUANTIFOIL® R 2/1). 3.2 µL of native inhibitor sample (WT) was applied onto a glow-discharged holey carbon copper grid (200 mesh, QUANTIFOIL® R 2/1). The grid was blotted and flash-frozen in liquid ethane with an FEI Mark IV Vitrobot. 3-3.5 µL of synthetic inhibitor sample was applied to a glow-discharged holey carbon copper grid (300 mesh, QUANTIFOIL® R 2/1). A Glacios Cryo-TEM was used to screen grids. CryoEM grids with optimal particle distribution and ice thickness were obtained by varying the blotting time/force. The Vitrobot chamber was set to 22 °C temperature and 100% humidity.

### Imaging

Optimized cryo-EM grids for inhibitor-bound β-galactosidase were loaded onto a Titan Krios (Thermo Fisher Scientific Inc) operated at 300 keV in eBIC (electron Bio-Imaging Centre, Diamond). The inhibitor-bound dataset was collected with a Gatan Quantum post-column energy filter (Gatan Inc) operated in zero-loss mode with 20 eV slit width, paired with a Gatan K3 direct electron detector, using EPU by electron counting in super-resolution mode (bin 2) at a physical pixel size of 0.829 per pixel. Optimized cryo-EM grids for control β-galactosidase were loaded onto another Titan Krios (Thermo Fisher Scientific Inc) operated at 300 keV in eBIC (electron Bio-Imaging Centre, Diamond). The control dataset was collected with a Gatan Quantum post-column energy filter (Gatan Inc) operated in zero-loss mode with 20 eV slit width, paired with a Gatan K3 direct electron detector, using EPU by electron counting in super-resolution mode (bin 2) at a physical pixel size of 0.831 Å per pixel. Optimized cryo-EM grids for synthetic inhibitor-bound β-galactosidase were loaded onto a third Titan Krios (Thermo Fisher Scientific Inc) operated at 300 keV in eBIC (electron Bio-Imaging Centre, Diamond). The synthetic inhibitor-bound dataset was acquired with a Thermo Scientific Selectris X imaging filter (Thermo Fisher Scientific Inc) with 10 eV slit width, paired with the latest generation Thermo Scientific Falcon 4i direct electron detector, using EPU by super-resolution counting mode in EER format at a physical pixel size of 0.921 Å per pixel. For all 3 datasets, a total number of 40 frames were acquired for each exposure, giving a total dose of ∼40 e−/Å^2^/micrographs, with a defocus range between −0.5 and −2.1 µm. Details of data collection parameters are listed in **Table S6**.

### Structure determination

The raw data was first corrected using a gain reference recorded immediately before the data acquisition, followed by generating a single micrograph that was corrected for overall drift using the MotionCor2 (*56*).The defocus value of each drift-corrected micrograph was determined by CTFFIND-4.1 (*57*) generating values ranging from −0.5 to −5.0 μm. Particle picking was done using the crYOLO program (*58*). Initially, a total of 3061973 particles were automatically picked from 15180 averaged images for native inhibitor-bound β-galactosidase, 1977809 particles from 11944 images for control β-galactosidase, and 2705966 particles from 10166 images for synthetic inhibitor-bound β-galactosidase, respectively.

All the 2D classification and 3D auto-refinement were done in RELION4.0 (*59*). The particles in 4k (super-resolution mode, bin 2) were boxed out in dimensions of 276 × 276 square pixels square for inhibitor-bound and control β-galactosidase, 250 × 250 square pixels square for synthetic inhibitor-bound β-galactosidase before further processing by the GPU accelerated RELION4.0. The first round 2D classification was done at a pixel size of around 2.39 Å for all 3 β-galactosidase samples. Several iterations of reference-free 2D classification were subsequently performed to remove bad particles (i.e., classes with fuzzy or un-interpretable features). 1406757, 1161378 and 2414813 good particles were re-extracted, and the second round 2D classification was done on the un-binned particles at pixel sizes of 0.829, 0.831 and 0.921 Å, yielding 659661, 593967 and 2353968 good particles for inhibitor-bound, control, and synthetic inhibitor-bound β-galactosidase respectively. 588492 particles from synthetic inhibitor-bound β-galactosidase were used for 3D auto-refinement, for better comparison with the other 2 datasets. D2 symmetry was imposed throughout processing. The two half-maps of each dataset from this auto-refinement step were subjected to RELION’s standard post-processing procedure. After auto-refinement and CTF refinement, the final maps of inhibitor-bound, control, and synthetic inhibitor-bound β-galactosidase achieved an averaged resolution of 1.93, 2.06, and 1.85 Å, respectively, based on RELION’s gold-standard FSC with cutoff at 0.143. The synthetic inhibitor-bound β-galactosidase had reached its Nyquist resolution, so we re-extracted the 588492 particles using RELION reconstruct and further processed the synthetic inhibitor-bound β-galactosidase dataset in 8k (super-resolution mode, bin 1) at a pixel size of 0.4605 Å. After several iterations of 3D auto-refinement and CTF refinement, the final overall resolution for the synthetic inhibitor-bound β-galactosidase is 1.42 Å. Prior to visualization, all density maps were sharpened by applying a negative B-factor calculated by RELION post-process (*59*).

### Model building and refinement

Atomic model building of the β-galactosidase protein chains were accomplished using ModelAngelo to build the initial β-galactosidase into our refined maps (*60*). For the inhibitor-bound structures, galactosyrin was manually fit into the map using COOT (*61*). Ions were then added to the models in COOT. Models refined via an iterative process using COOT and PHENIX real-space refinement (*61*, *62*). Water molecules were added using PHENIX douse function before final real-space refinement (*62*).

The cryoEM maps and corresponding PDB models for control (Δ*gsn*), native inhibitor-bound (WT), and synthetic inhibitor-bound have been deposited under accession number 8RI6, 8RI7, and 8RI8, and EMDB accession numbers EMD-19181, EMD-19182, and EMD-19183, respectively.

## Gas chromatography-mass spectrometry (GC-MS)

### Sample preparation

To extract soluble metabolites, sample was mixed with an equal volume of a mixture of methanol:chloroform (2.5:1 ratio), centrifuged at 12,000 × g for 5 minutes then the top aqueous fraction was collected and freeze dried. The dried sample was dissolved in pyridine with 20 mg/ml methoxyamine HCl (for carbonyl-containing compounds such as galactosyrin aldehyde and untargeted metabolomics) or pyridine only (for all other samples) and incubated at 37°C for 30 minutes. Then the sample was derivatised with MSTFA (N-Methyl-N-trimethylsilyl trifluoroacetamide) at 37°C for 30 minutes. The resulting mixture was centrifuged at 12,000 × g for 5 minutes and the supernatant was transferred into vials for GC-MS.

### GC-MS

GC-MS was conducted using Intuvo 9000 GC system with 5977B single quadrupole MSD system and 7693 autosampler (Agilent). The GC system was equipped with a guard chip and two Agilent 122-5512UI-INT columns connected sequentially. The sample (0.5 µl) was injected in splitless mode at 230°C. Helium was used as carrier gas and the flow was set constant at 1 ml/min through the first column and 1.2 ml/min through the second column. The oven temperature was initially held at 70°C for 5 min then increased to 330°C at the rate of 7.5°C/min. The transfer line temperature was 250 °C. The sample was ionised by electron impact ionisation at 70 eV. The MS source temperature was 230°C and the MS quadrupole temperature was 150°C. The mass spectrometer was autotuned before running each batch of samples. The acquisition of mass spectra was in scan mode for ions from 50 to 600 mass units with the scan speed of n=2. Solvent delay of 7.5 min was set before acquisition. Data were analysed using MassHunter qualitative analysis software (Agilent).

### Chemical standards

1ADR (1-Amino-1-deoxy-D-ribitol hydrochloride, Supelco 72679, CAS no. 22566-17-2), galactosylglycerol ((2R)-Glycerol-O-β-D-galactopyranoside, Cayman Chemical CAY17218, CAS no. 16232-91-0), trehalose (Sigma T9531, CAS no. 6138-23-4)

### Chemical synthesis

The synthesis scheme is outlined in **Fig. S13**.

### Chemicals and other materials

All reactions were carried out under an argon atmosphere. Reagents and (dry) solvents were commercially obtained from Acros Organics, Merck, VWR Chemicals, Fisher Chemical and Carbolution Chemicals, compound **1** was obtained from AstaTech (P10778, CAS no. 153172-31-7). Flash chromatography was performed on silica gel from Thermo Scientific with a particle size of 60-200 μm and an average pore size of 60 Å. Thin layer chromatography (TLC) analysis was carried out on Merck 60F_254_ silica gel plates.

### HPLC and mass spectrometry

Preparative HPLC reversed-phase chromatography was performed on a Shimadzu Prominence system with a LC-20AP pump equipped with a Phenomenex Luna^®^ C18(2) 100Å column (5 μm, 100 x 21.2 mm). Analytical LC-MS (ESI) analysis was performed on a Thermo Scientific UltiMate 3000 HPLC system with a Thermo Finnigan LCQ Fleet ion trap mass spectrometer, using a gradient program with eluent A = MQ-water with 0.1% formic acid and eluent B = acetonitrile with 0.1% formic acid, 0.5 min 10% B, then 10% to 100% B in 5.5 min, then 3.2 min 100% B, a flow rate of 1.0 mL min^-1^ on a Macherey-Nagel NUCLEODUR C18 Pyramid column (particle size 5 μm, 250 mm x 4.0 mm) at 25 °C. HRMS (ESI) analyses were performed on a Thermo Scientific Exactive Plus EMR mass spectrometer.

### Nuclear magnetic resonance

NMR spectra were recorded with a Bruker 400 MHz Avance II spectrometer, equipped with a PATXI probe, with a ^1^H frequency of 399.99 MHz and ^13^C frequency of 100.59 MHz. Residual ^1^H resonance from the deuterated solvents was used to reference the ^1^H spectra with the methyl resonance of tetramethylsilane (δ = 0.00 ppm), the ^13^C spectra were referenced through the solvent ^13^C resonance.

### *tert*-Butyl (3a*R*,4*R*,6a*S*)-4-(hydroxymethyl)-2,2-dimethyl-tetrahydro-[1,3]dioxolo[4,5-*c*]pyrrole-5-carboxylate (2)

A solution of compound **1** (67.4 mg, 234 µmol, 1.0 equiv.) in a mixture of dry MeOH (600 µL) and triethylamine (200 µL) was treated with di-*tert*-butyl dicarbonate (76.6 mg, 351 µmol, 1.5 equiv.) and stirred for 2 hours at room temperature. The resulting mixture was quenched by the addition of EtOAc (20 mL) and saturated aqueous NH_4_Cl (10 mL). The organic layer was separated, washed with 1 M hydrochloric acid (10 mL) and brine (10 mL), dried over Na_2_SO_4_, filtered, and concentrated *in vacuo*. The obtained crude residue was dissolved in a 1.0 M solution of *N*, *N*, *N*-tributylbutan-1-aminium fluoride (1.0 mL, 1.0 mmol, 4.3 equiv.) and stirred for 2.5 hours at room temperature. The resulting mixture was concentrated *in vacuo*, re-dissolved in DCM (10 mL), washed with saturated aqueous NH_4_Cl (10 mL) and brine (10 mL), and concentrated *in vacuo*. The obtained crude residue was purified by flash column chromatography (6 g SiO_2_, EtOAc/cyclohexane 1:9 → 1:1) to provide alcohol **2** as a clear colorless oil (38.6 mg, 60% over two steps). *R_f_* = 0.3 (SiO_2_, EtOAc/cyclohexane 1:1). HPLC retention time = 6.7 min. MS (ESI) *m/z* (%) = expected 273.16; (pos) 174.29 ([M+2H-Boc]^+^, 46), 218.08 ([M+2H-^t^Bu]^+^, 100), 273.92 ([M+H]^+^, 4). ^1^H NMR (400 MHz, DMSO-D_6_) δ 4.97 – 4.86 (m, 1H), 4.72 – 4.65 (m, 1H), 4.65 – 4.60 (m, 1H), 3.90 – 3.72 (m, 1H), 3.57 – 3.43 (m, 2H), 3.43 – 3.34 (m, 2H), 1.39 (s, 9H), 1.33 (s, 3H), 1.24 (s, 3H). ^13^C NMR (101 MHz, DMSO-D_6_) (two rotamers) δ 153.5, 110.4, 82.4, 81.6, 79.1, 78.7, 78.6, 78.3, 65.0, 64.5, 60.8, 60.2, 52.8, 52.2, 28.1, 26.8, 24.7.

### (2S,3R,4S)-2-(dihydroxymethyl)pyrrolidine-3,4-diol (3) via 2

A solution of alcohol **2** (33.2 mg, 122 µmol, 1.0 equiv.) in wetted DCM (1 mL) was treated with Dess-Martin reagent (77.0 mg, 182 µmol, 1.5 equiv.) and stirred for 3 hours at room temperature. The resulting mixture was concentrated *in vacuo* and purified by preparative reversed phase chromatography (C18, acetonitrile/water with 0.1% TFA) to provide the intermediate aldehyde after drying *in vacuo* (HPLC retention time = 6.2 min. MS (ESI) *m/z* (%) = expected 271.14; (pos) 172.15 ([M+2H-Boc]^+^, 100), 215.94 ([M+2H-^t^Bu]^+^, 49)). The resulting intermediate aldehyde was immediately re-dissolved in a 2 M HCl solution in THF/water (1:1, 16.6 mL), stirred for 4.5 h at room temperature and subsequently concentrated *in vacuo*. The crude residue was purified by preparative reversed phase chromatography (C18, acetonitrile/water) to provide compound **4** as a slightly brownish solid (4.3 mg, 21% over two steps). HPLC retention time = 1.8 min. MS (ESI) *m/z* (%) = expected 149.07; (pos) 132.01 ([M+H-H_2_O]^+^, 28), 149.97 ([M+H]^+^, 100). ^1^H NMR (400 MHz, D_2_O) δ 5.38 (d, *J* = 4.6 Hz, 1H), 4.45 – 4.37 (m, 1H), 4.36 (dd, *J* = 7.3, 4.2 Hz, 1H), 3.55 (dd, *J* = 7.2, 4.8 Hz, 1H), 3.47 (dd, *J* = 12.7, 3.8 Hz, 1H), 3.38 (dd, *J* = 12.7, 2.2 Hz, 1H). ^13^C NMR (101 MHz, D_2_O) δ 87.0, 71.6, 69.7, 64.6, 49.6.

Alternatively, (2*S*,3*R*,4*S*)-2-(dihydroxymethyl)pyrrolidine-3,4-diol (**3**) could also be obtained as a mixture via intermediate **4**: To this end, the crude compound **4**, obtained via the Kamath *et al.* procedure (*37*) using **1** (100 mg, 348 µmol, 1.0 equiv.) was dissolved in a 2 M HCl solution in THF/water (1:1, 6 mL) and stirred for 18 h at room temperature. The resulting mixture was concentrated *in vacuo* and purified by preparative reversed phase chromatography (C18, acetonitrile/water) to provide a brownish solid (10.2 mg), consisting predominantly (according to NMR) of compound **3**.

### Full-length sequence of *gsnA*

*gsnA* (PSPTO_0834) was originally misannotated with a truncation. The recently updated locus tag PSPTO_RS04425 is correctly annotated. The full-length sequence is shown below.

>*gsnA*_CDS ATGAAAGCACTGGGCTTAATGGATAACCAGAGGCTTGAACTTGTGGATTGCAAGGATCCTGTCATGCTTGCTCCTGACGGCGTAGAAATCGATATCGTGCTATCAGGTATATGCGGAACTGATCTGGCGGTATTGTCGGGCCGTGAAGGTGGAGAGGTGGGCATTATACGCGGGCACGAAGCAGTTGGCATTATTATCGATGTAGGTAAGGATGTAGTACACCTACAAAAAGGGATGCGGGTGGTGGTTGATCCCAACGAATACTGTGGCGTTTGCGAACCTTGCCGTCTTGCTAAAACGCACCTATGCAATGGGGGGGTGAACGCTGGGTTGGATATCGCAGGTGTCAACAAACATGGAACTTTTGCCGAGCGCTTCGTTACTCGTGAGCGTTTTGTGTATCAATTGCCAGACGATATGAGCTGGGCAGCTGGTGTGTTGGTTGAGCCTGTTGCCTGCATTCTGAATAATATAGACCAGGCGTTCATTCGAGCGGGAGAGCGTGTGTTGATCCTAGGGTCTGGCCCTATGAGTCTGATTGCGCAGATCGTTCTGCGCTCAATGGGAGTTGACACGCTCGCCACTGATCGAAACACACATCGCATACAGTTCGGCCGCTCACAAAGTCTTGATGTTATACATGCCGATGATCTTGAGTTGCAGATGCAGCACCAAGAAAAGTTTGATGTTGTTATCGATACTGTCGGTAATCAGATCGATACAGCTTCACGCTACATCGGTCGCGGTGGGAGAATTGTACTTTTTGGATTTGATAGTGACTATCACTACATGCTGCCTGTAAAGTACTTCCTGGTTAACGCTATCAGTATTATTTCTGCTGGAGAATACAATCAGCACTTTCCTAGAGCAATTCGTCTTGTGCAAAAACTTCCTGAGCTAGGGCGGCTGGTAACGCATCGCTACGTACTAGAAAATCACTCGGAGGTTTTCGATGCACTTCTGAACGATGCTTCCGCCCCCAATATAAAAAGCGTATTCACACCAAATCTCGCTTATCTTTAA

>GsnA_protein MKALGLMDNQRLELVDCKDPVMLAPDGVEIDIVLSGICGTDLAVLSGREGGEVGIIRGHEAVGIIIDVGKDVVHLQKGMRVVVDPNEYCGVCEPCRLAKTHLCNGGVNAGLDIAGVNKHGTFAERFVTRERFVYQLPDDMSWAAGVLVEPVACILNNIDQAFIRAGERVLILGSGPMSLIAQIVLRSMGVDTLATDRNTHRIQFGRSQSLDVIHADDLELQMQHQEKFDVVIDTVGNQIDTASRYIGRGGRIVLFGFDSDYHYMLPVKYFLVNAISIISAGEYNQHFPRAIRLVQKLPELGRLVTHRYVLENHSEVFDALLNDASAPNIKSVFTPNLAYL

---

**Fig. S1.**
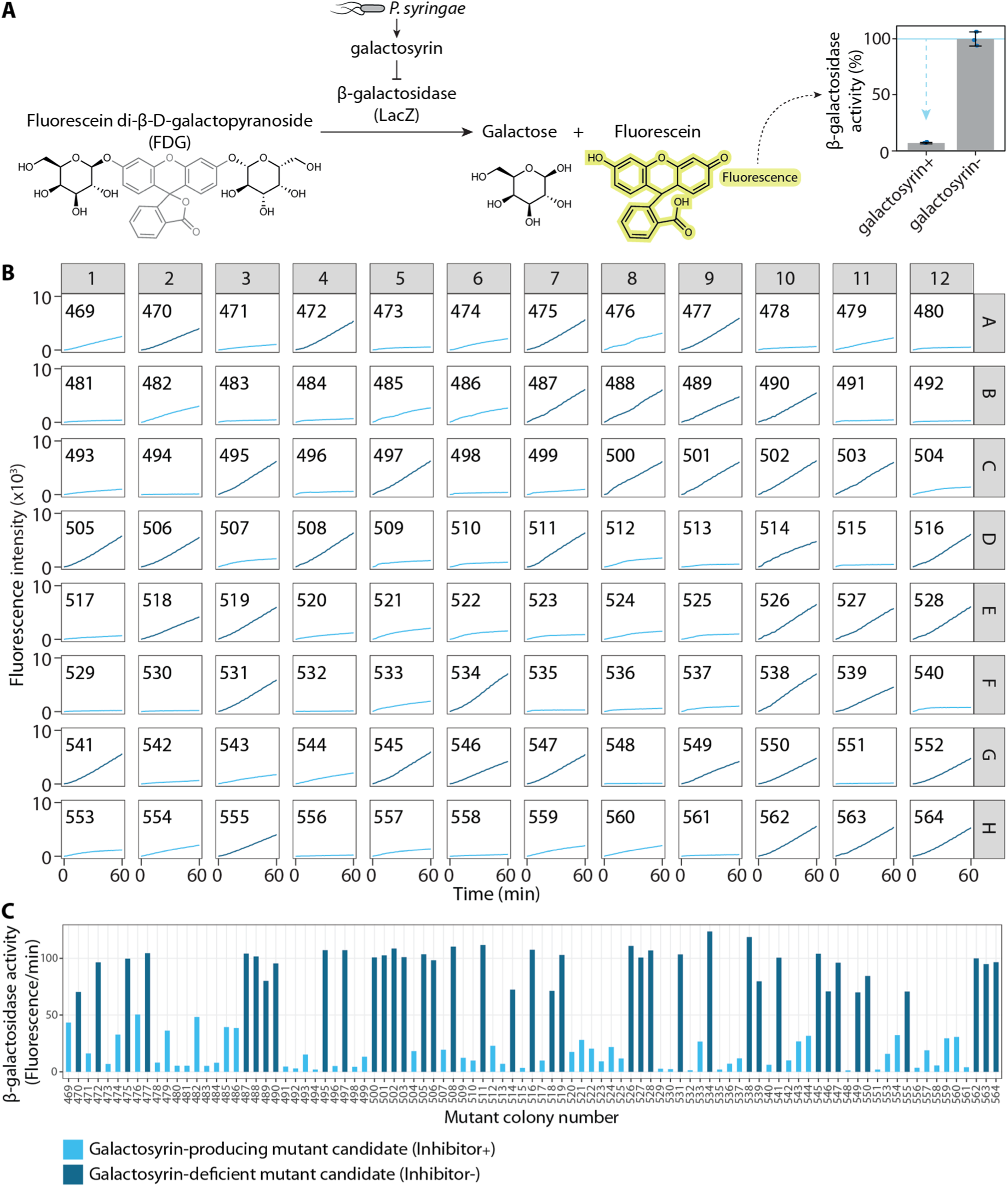
Validation screen for candidate galactosyrin mutants. **(A)** The enzymatic assay to monitor β-galactosidase inhibitor production. The cleavage of FDG substrate by β-galactosidase (LacZ) produces a fluorescent product fluorescein. The rate of fluorescence increase over time can be calculated as a measure of enzyme activity. The addition of galactosyrin produced by the bacteria inhibits LacZ activity. **(B)** Candidate galactosyrin mutants selected from plates were numbered and grown in virulence-inducing MG medium in each well of a 96-well plate overnight. The supernatants were then tested for the inhibitor in an enzymatic assay with FDG substrate and LacZ enzyme. Fluorescence signal from cleavage product of FDG by LacZ was measured over time to monitor enzyme activity and inhibition. **(C)** β-galactosidase activity was calculated for each mutant and was classified into 2 groups: galactosyrin-producing mutant (Inhibitor+, light blue), causing low enzyme activity in the assay, and galactosyrin-deficient mutant (Inhibitor-, dark blue), causing high enzyme activity in the assay. The validated Inhibitor-mutants were selected for sequencing and mapping of inhibitor-related genes.

**Fig. S2.**
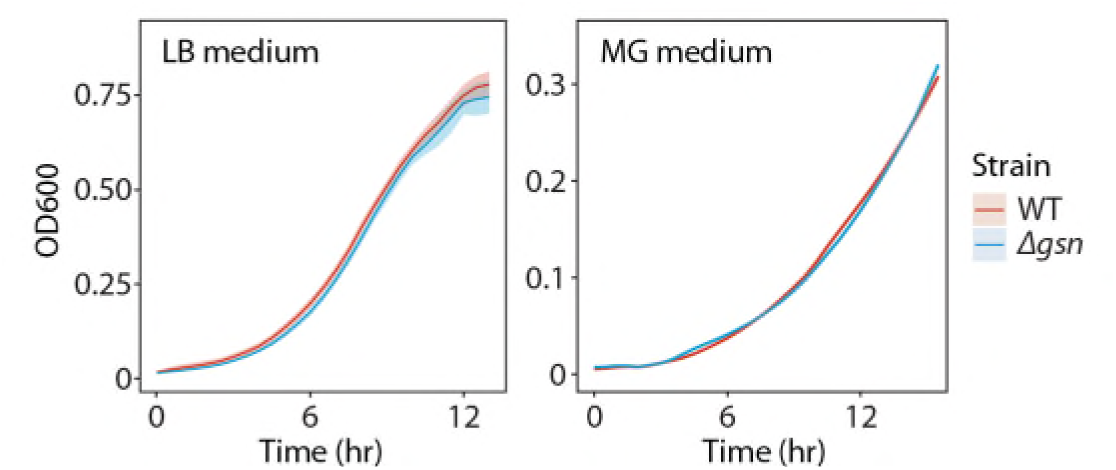
Deletion of the *gsn* cluster does not affect bacterial growth *in vitro*. Bacterial strains (wild-type (WT) or *gsn* cluster knockout (*`Δ`gsn*)) were inoculated in LB medium or virulence-inducing MG medium in a 96-well plate and grown at 28 °C. Optical density at 600 nm (OD600) was measured over time to monitor bacterial growth. Mean and standard deviation from 3 replicates are plotted.

**Fig. S3.**
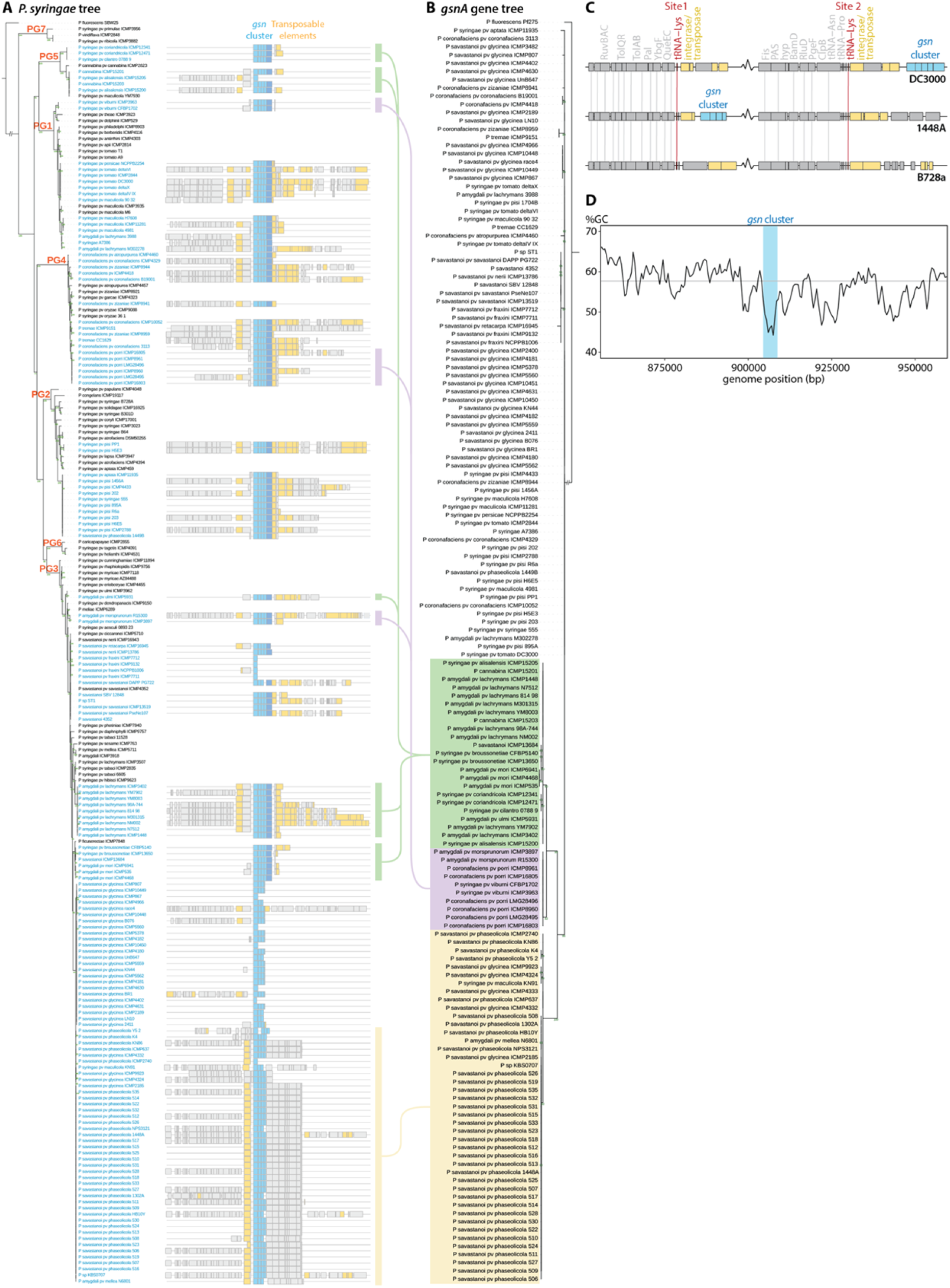
The distribution of the *gsn* cluster among *P. syringae* strains is indicative of horizontal gene transfer. **(A)** Phylogenetic tree of *P. syringae* strains. Maximum likelihood phylogenetic tree was constructed using concatenated coding DNA sequences of 4 conserved genes (*rpoD*, *gyrB*, *gltA* and *gapA*). Phylogroups (PG) are labelled with orange texts on each clade. *P. fluorescens* SBW25 is included as an outgroup. Strains with the *gsn* cluster are highlighted in blue. Genomic regions (20 kbp) around the *gsn* cluster are displayed on the right of each strain with annotated genes displayed as rectangles, the *gsn* cluster highlighted in blue and transposable elements highlighted in yellow. Many genomes are not completely assembled so truncated genomic regions are commonly observed. **(B)** Phylogenetic tree of *gsnA* genes. Maximum likelihood phylogenetic tree was constructed using coding DNA sequences of *gsnA* homologs identified from BLAST against genomes of *P. syringae* strains. A homolog from *P. fluorescens* Pf275 is included as an outgroup. Selected clades of closely related *gsnA* sequences are highlighted in the same colour (green, purple or orange) with lines connecting to the corresponding strains in the *P. syringae* tree, showing phylogenetic incongruence indicative of horizontal gene transfer. (B, C) Branch support values are calculated from ultrafast bootstrap with 1,000 replications. Branches with support values of less than 80 are collapsed. Branch lengths are drawn to scale representing the number of substitutions per site. **(C)** *gsn* clusters are found downstream of one of the two tRNA-lysine sites in the genome. Genomic regions surrounding the two tRNA-lysine sites (red) in the genomes of three *P. syringae* reference strains. In the strains with the *gsn* cluster (blue) (DC3000 and 1448A), the *gsn* clusters are found associated with transposable elements (yellow) downstream of the tRNA^lys^ site, which are usual targets for integration. The *gsn* cluster is absent from the B728a strain. **(D)** *gsn* cluster has a lower GC content than average. Genome sequence of *P. syringae* pv. *tomato* DC3000 was analysed for percent GC content in 500 bp sliding windows across a 100 kbp region surrounding the *gsn* cluster (blue). Grey line indicates the overall GC content of the genome. A GC content different from the genome is suggestive of horizontal gene transfer origin.

**Fig. S4.**
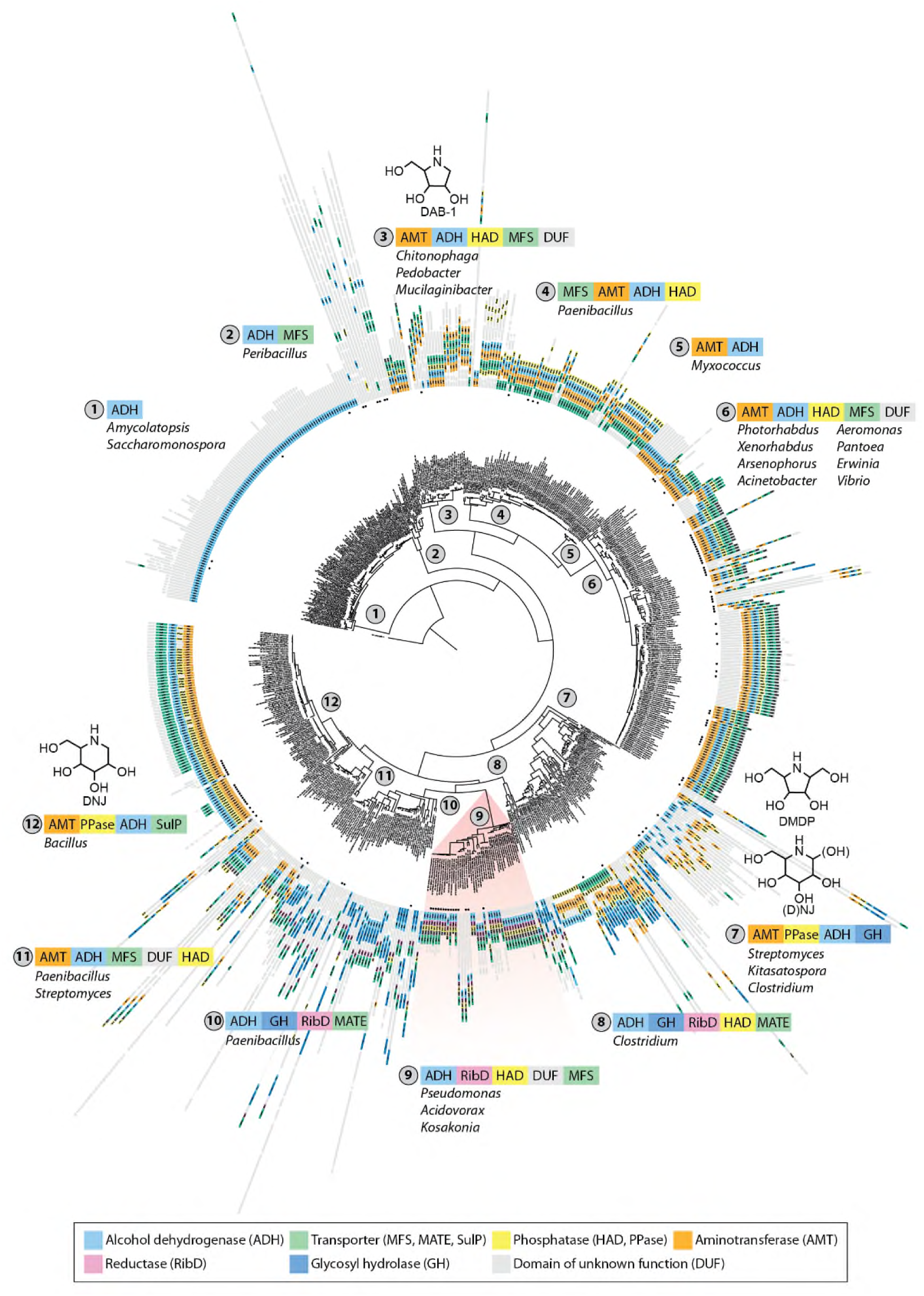
*gsnA* homologs are part of different putative iminosugar biosynthesis gene clusters. Maximum likelihood phylogenetic tree was constructed using amino acid sequences of *gsnA* homologs [alcohol dehydrogenase (ADH)] identified by PSI-BLAST on Refseq database. Each homolog is labelled with its protein ID and the names of the species in which it is found. For brevity, a maximum of 5 homologs identified from each species are included. Branch support values are calculated from ultrafast bootstrap with 1,000 replications. Branches with support values of less than 80 are collapsed. Branch lengths are drawn to scale representing the number of substitutions per site. Threonine dehydrogenase from *E. coli* is included as an outgroup. Genomic regions of putative gene clusters are displayed next to each entry with annotated genes displayed as rectangles and selected commonly occurring genes and families are highlighted in different colours. Solid black dots indicate gene cluster association with transposable elements. Homologs and clusters are classified into 12 distinct clades (labelled with numbers in a circle) based on tree topology. Representative gene cluster composition and bacterial genus are shown for each clade. Iminosugars reported to be produced by members of the clades are shown (*7–9*). Clade 9 containing the *P. syringae gsn* cluster is highlighted in red.

**Fig. S5.**
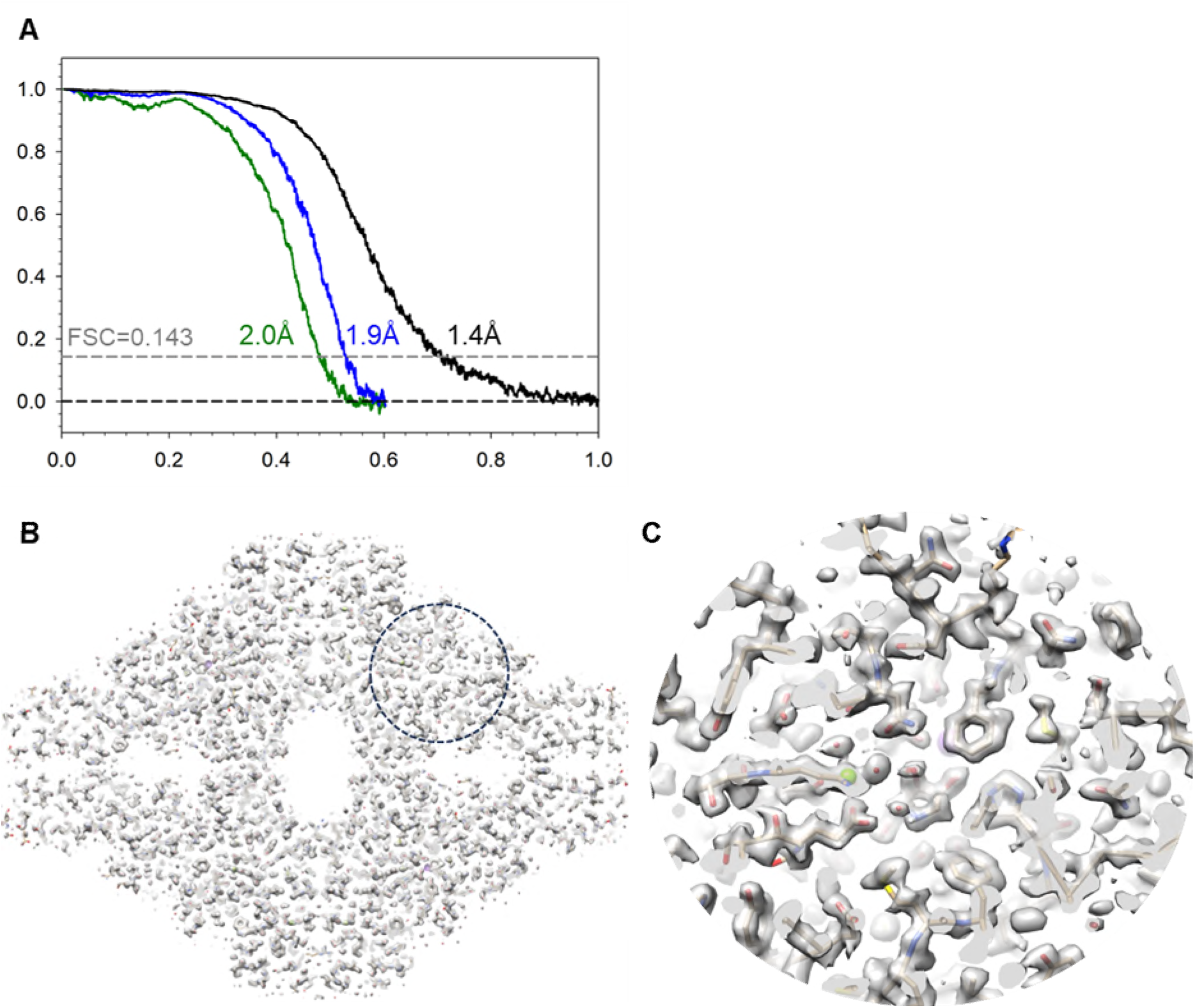
Cryo-EM structure determination of β-galactosidase (LacZ) bound with galactosyrin. **(A)** Fourier Shell Correlation (FSC) between the two independently refined half-maps for LacZ in complex with the native inhibitor (WT, blue), the negative control Δ*gsn* (green), and with the synthetic galactosyrin (black). **(B)** Cryo-EM map of LacZ bound with the synthetic galactosyrin, shown with a central slice for clarity. **(C)** A zoom-in view of the LacZ active site circled in (B). Atomic details of the density map are clearly displayed.

**Fig. S6.**
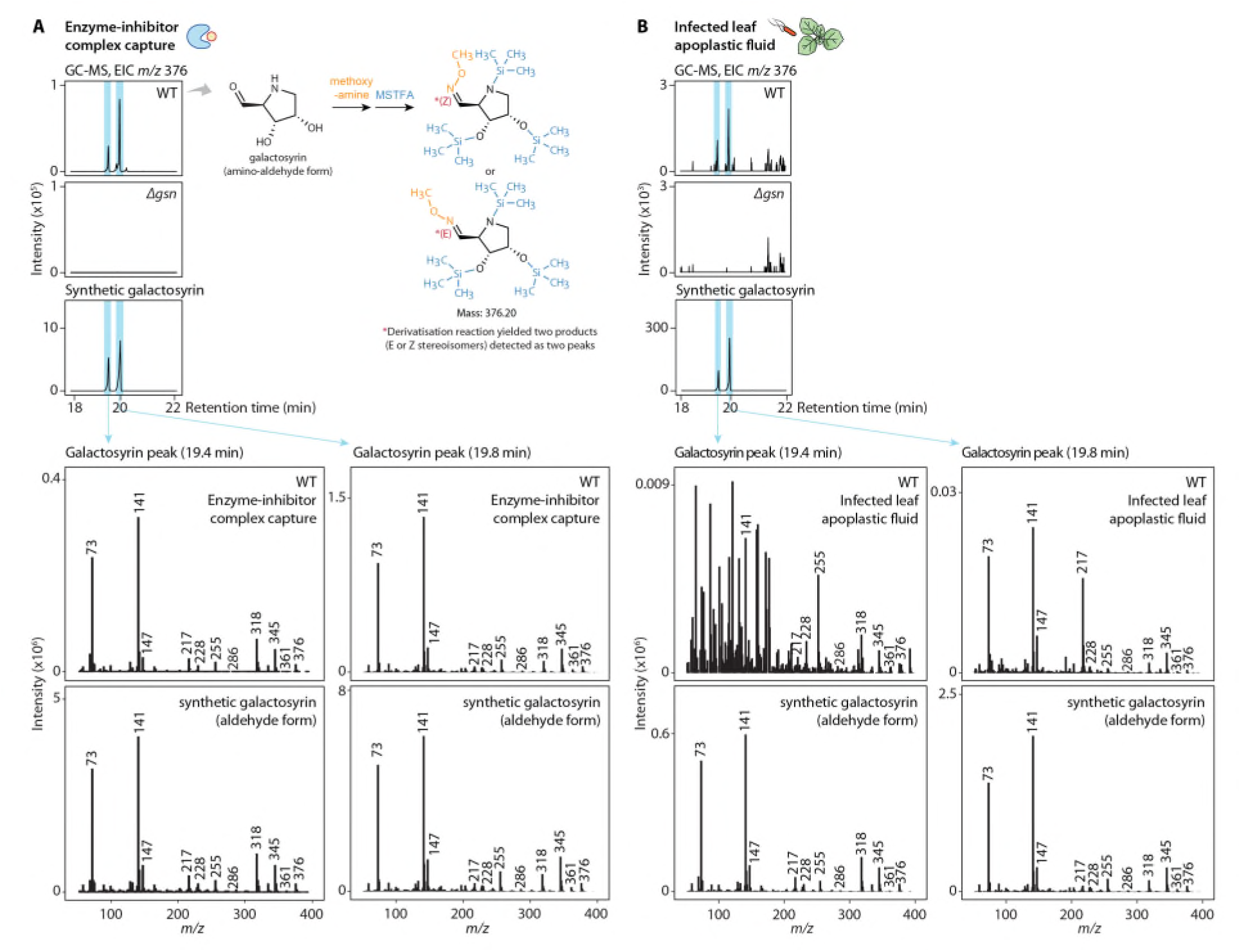
Galactosyrin was detected in enzyme-inhibitor complex and apoplast of infected plants. (Top) Extracted ion chromatogram (EIC) of *m/z* 376 from GC-MS analysis of synthetic galactosyrin and soluble metabolite extracts from **(A)** enzyme-inhibitor complex capture from cultures of galactosyrin-producing (WT) or galactosyrin-deficient (*`Δ`gsn*) strains (Fig. 2) and **(B)** apoplastic fluid from *N. benthamiana* leaves infected with WT or *`Δ`gsn* strains. Samples were analysed by GC-MS after chemical modification with methoxyamine, which modifies the aldehyde group, and N-Methyl-N-(trimethylsilyl)trifluoroacetamide (MSTFA), which modifies hydroxyl and amine groups, to enable carbohydrate analysis. Since methoxyamine modification yields two stereoisomers, E and Z (*34*), galactosyrin was detected as two peaks. Blue stripes highlight galactosyrin peaks. (Bottom) Mass spectra of the detected galactosyrin peaks showing identical expected masses. Compound structures, derivatisation and mass annotations are shown in **Fig. S9**. The galactosyrin peak (19.4 min) detected in the infected apoplastic fluid sample had a relatively low intensity and co-eluted with an unknown metabolite form the plant apoplast so high background noise was observed in the mass spectrum.

**Fig. S7.**
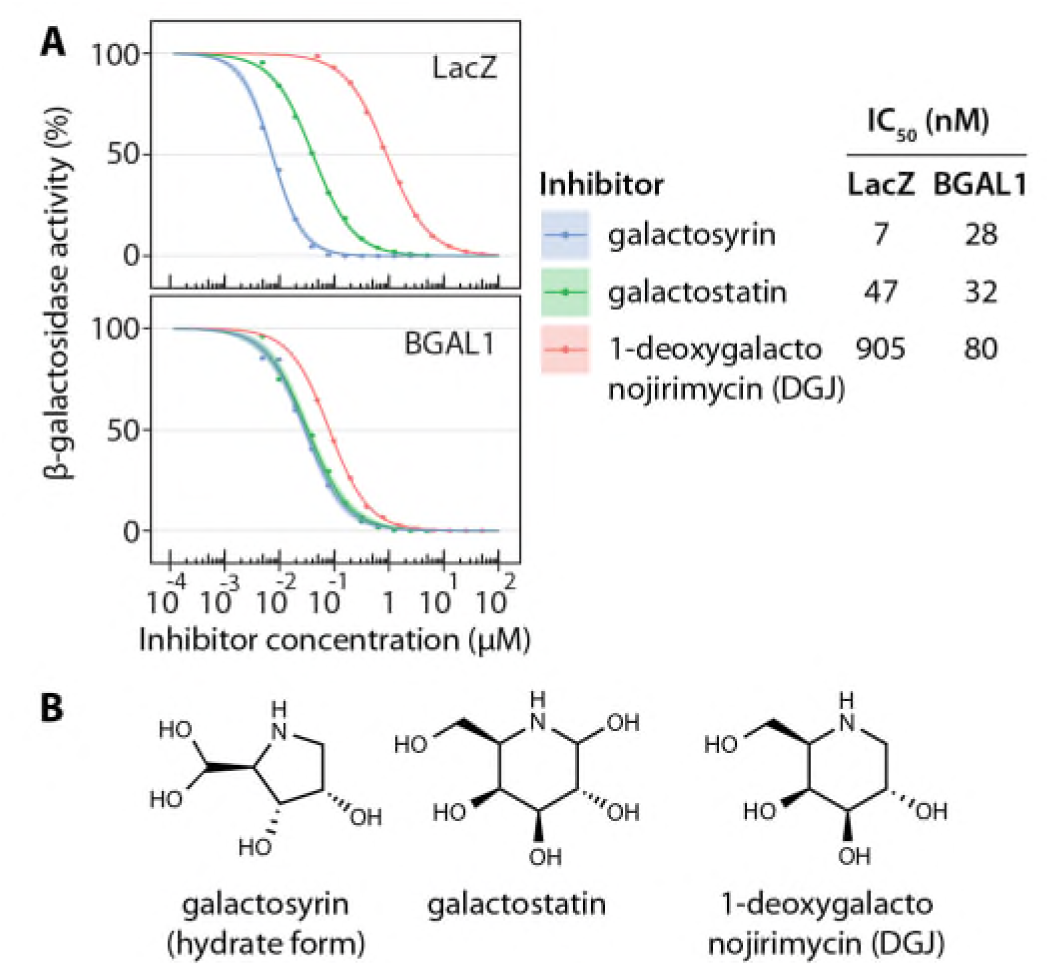
Galactosyrin is a potent β-galactosidase inhibitor. **(A)** Dose-response curves of galactosyrin, galactostatin and 1-deoxygalactonojirimycin (DGJ) inhibition of LacZ or BGAL1 with FDG substrate. β-galactosidase activity is expressed as a percentage of the activity relative to no inhibitor control. A four-parameter logistic model was fitted and half maximal inhibitory concentration (IC50) is shown for each enzyme and inhibitor combination. (B) Structures of inhibitors tested.

**Fig. S8.**
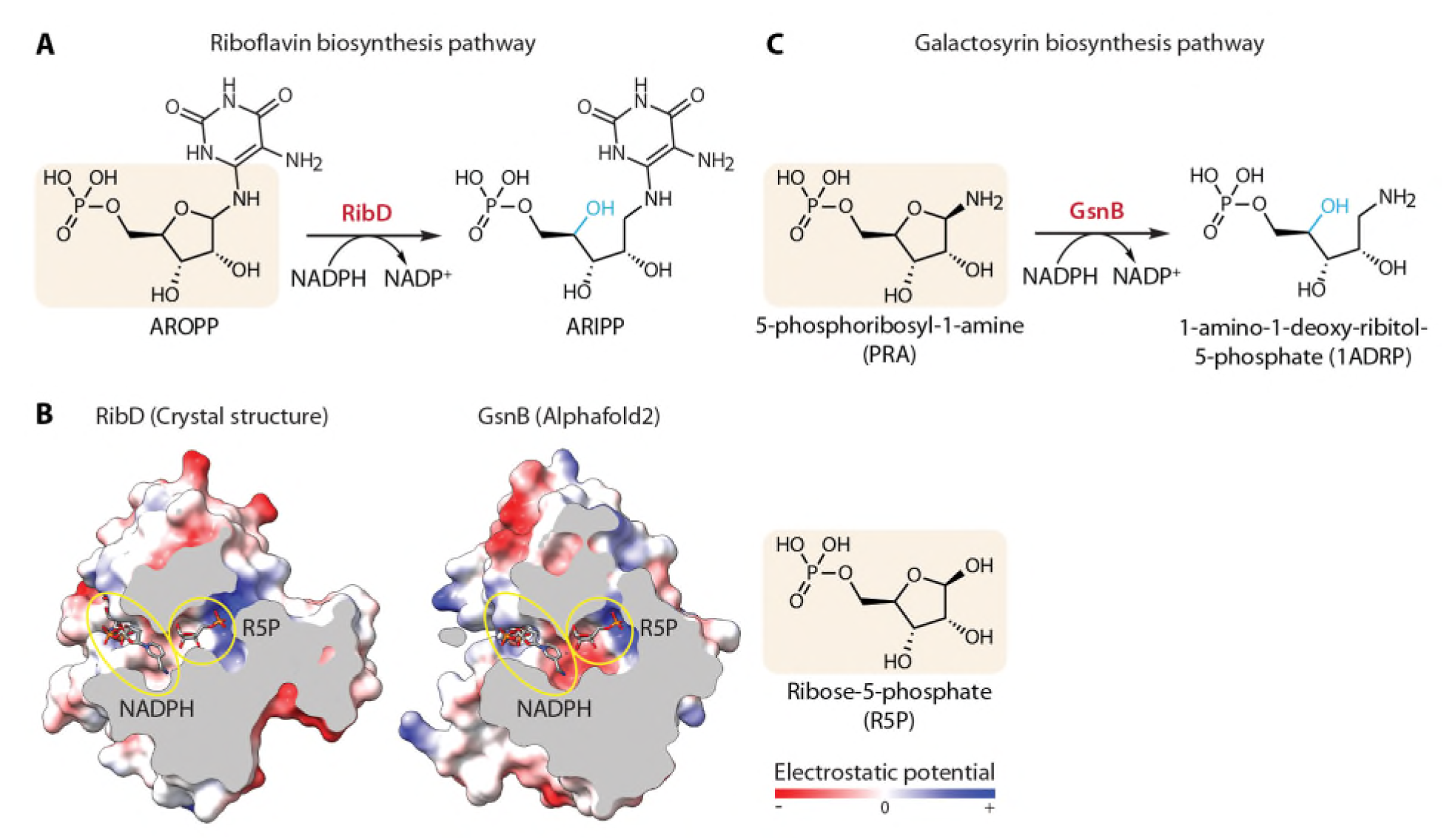
Homology and structural modelling predict GsnB function. **(A)** RibD, a close domain family member of GsnB, functions in the riboflavin biosynthesis pathway by catalysing the reduction of 5-amino-6-ribosylamino-2,4(1H,3H)-pyrimidinedione 5′-phosphate (AROPP) to 5-amino-6-ribitylamino-2,4(1H,3H)-pyrimidinedione 5′-phosphate (ARIPP). **(B)** Comparison of crystal structures of RibD from *E. coli* in complex with the ribose-5-phosphate (R5P) substrate analog (PDB: 2OBC) or NADPH cofactor (PDB: 2O7P) and Alphafold2 predicted structure of GsnB (RMSD (angstrom) of 1.07 between 125 pruned atom pairs, 7.79 across all 187 pairs compared to 2OBC and 1.06 between 125 pruned atom pairs, 4.53 across all 183 pairs compared to 2O7P). The enzyme structure cross section shows the conserved active site pockets (yellow circles) with the overlay of the structure of the substrate analog (R5P) and cofactor (NADPH). Protein structures are coloured based on electrostatic potential of amino acid residues (red: negative - blue: positive) to show the conserved properties at the sites for substrate and cofactor binding. **(C)** GsnB was predicted then proven to function in galactosyrin biosynthesis pathway by catalysing the reduction of 5-phosphoribosyl-1-amine (PRA) to 1-amino-1-deoxy-ribitol-5-phosphate (1ADRP). Chemical change in the product compared to the substrate is highlighted in blue. Similar structures of the substrates and analogs are highlighted in rounded rectangle.

**Fig. S9.**
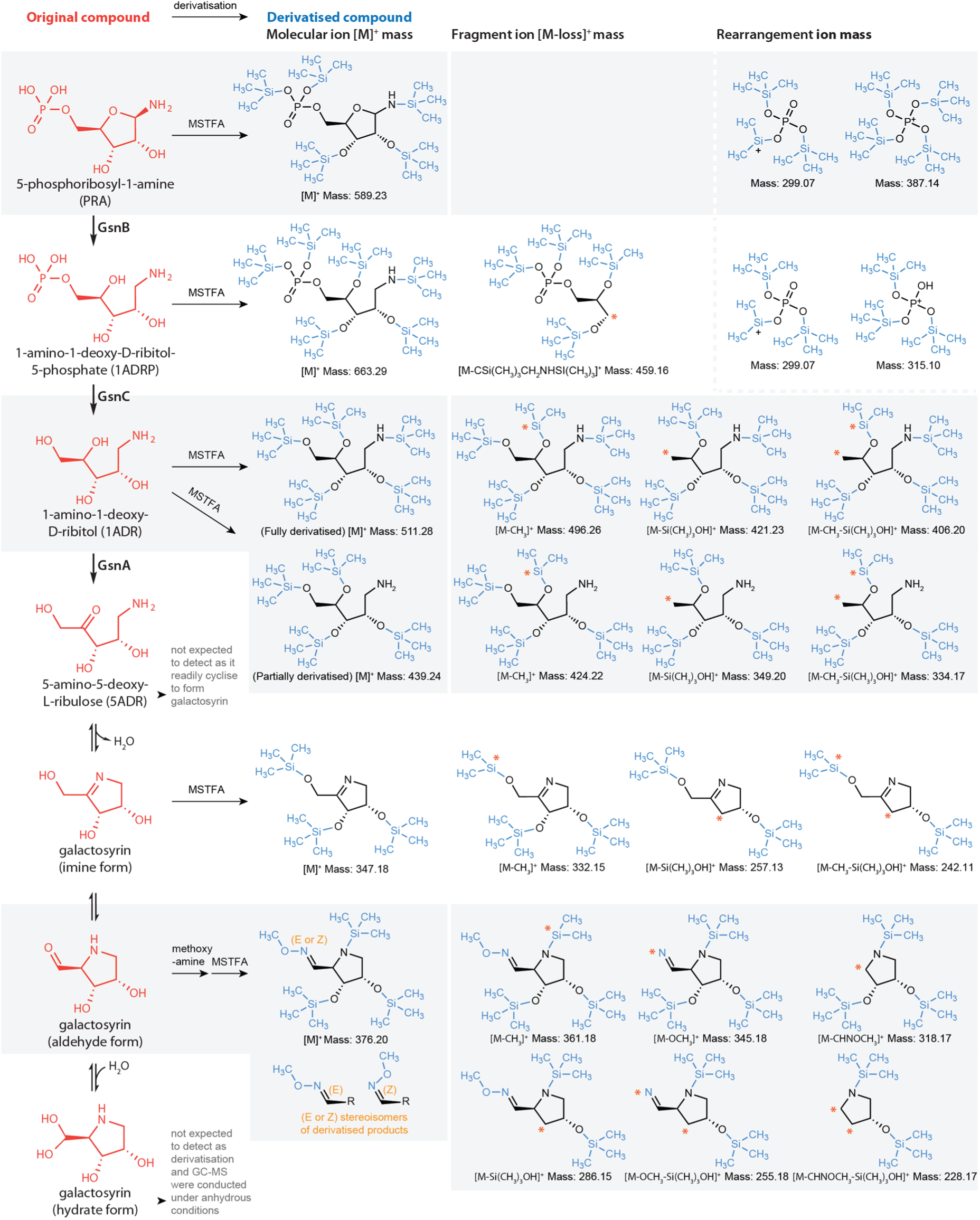
Derivatised products and fragments detected by GC-MS. (Left) Compounds in the galactosyrin biosynthesis pathway with structures shown in red and chemical derivatisation for GC-MS analysis indicated on arrows. MSTFA (N-Methyl-N-(trimethylsilyl)trifluoroacetamide) derivatises hydroxyl and amine groups, while methoxyamine derivatises aldehyde groups yielding two products with stereoisomers E or Z, detected as two peaks for the aldehyde form of galactosyrin. Compounds not expected to be detected in GC-MS are indicated with a description in grey. (Right) Derivatised compounds with the derivatisation on the original structure shown in blue. The mass expected in GC-MS detection are indicated below each structure. Molecular ion shows the intact derivatised compounds. Fragment ion shows expected fragments from the fragmentation of molecular ion caused by electron ionisation (EI) during GC-MS. Orange asterisks indicate the group lost by fragmentation. Rearrangement ion shows expected products from rearrangement of the molecular and fragment ions during GC-MS. Data interpretation was facilitated by general principles of derivatisation, fragmentation and rearrangement in GC-MS (*35*, *36*).

**Fig. S10.**
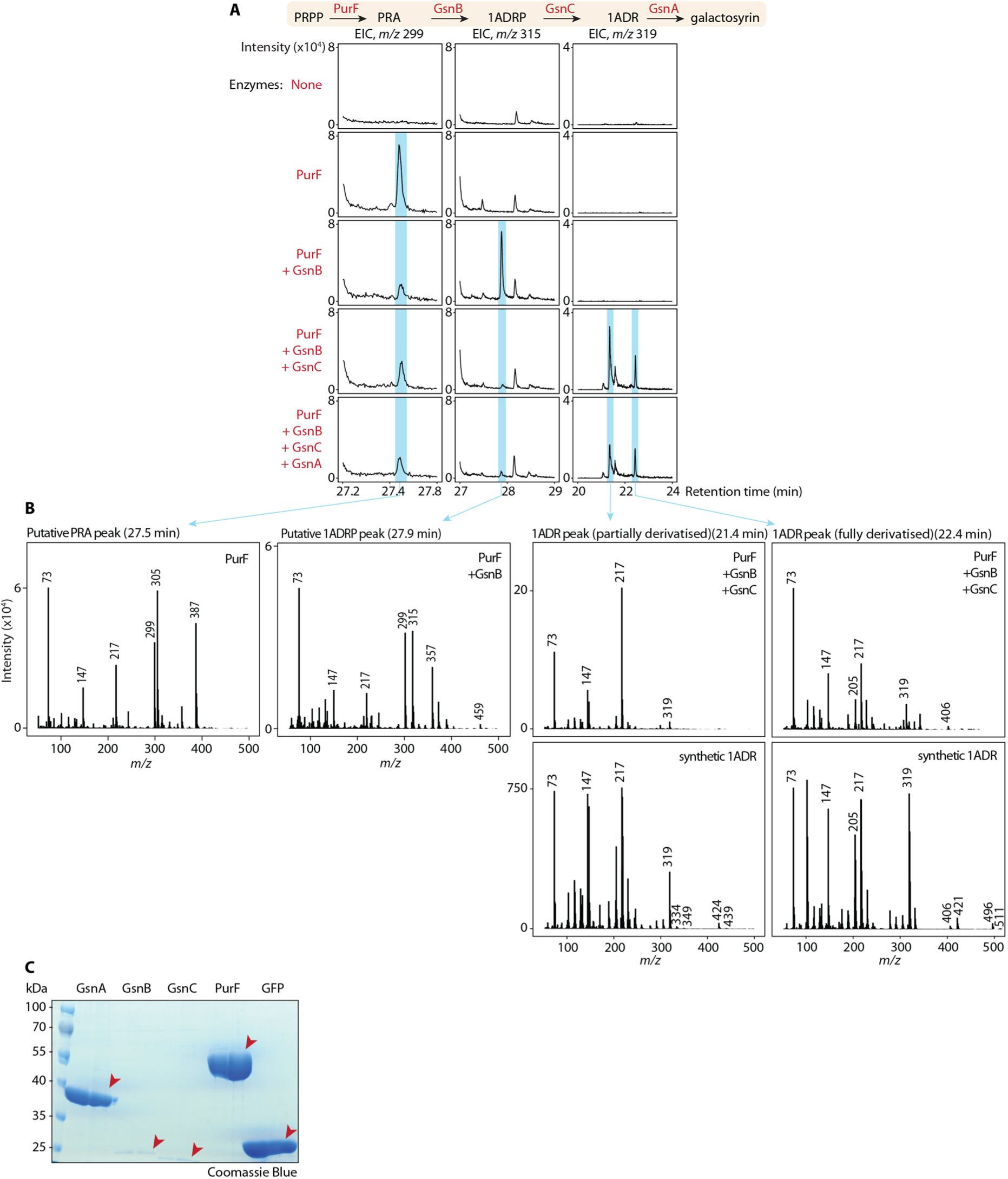
Detection of intermediates in galactosyrin biosynthesis *in vitro*. **(A)** Summary diagram of the biosynthesis pathway is shown at the top. Extracted ion chromatograms (EIC) of mass signature corresponding to each intermediate (PRA, 1ADRP and 1ADR) are shown in each column. For each row, each mixture of enzymes (listed in red on the left) were mixed with the starting precursor (PRPP) and cofactors to reconstruct the biosynthesis pathway, then soluble metabolite fractions were analysed by GC-MS. Blue stripes highlight peaks of the detected intermediates, accumulating in samples containing enzymes that produce them and decreasing in samples containing enzymes that use them. **(B)** Mass spectra of the detected peaks. For 1ADR, the matching spectra of a synthetic standard were also shown underneath. Compound structures, derivatisation and mass annotations are shown in **Fig. S9**. **(C)** Purified enzymes used in the experiment. Proteins were separated on SDS-PAGE and stained with Coomassie blue. Orange arrows indicate the expected protein band.

**Fig. S11.**
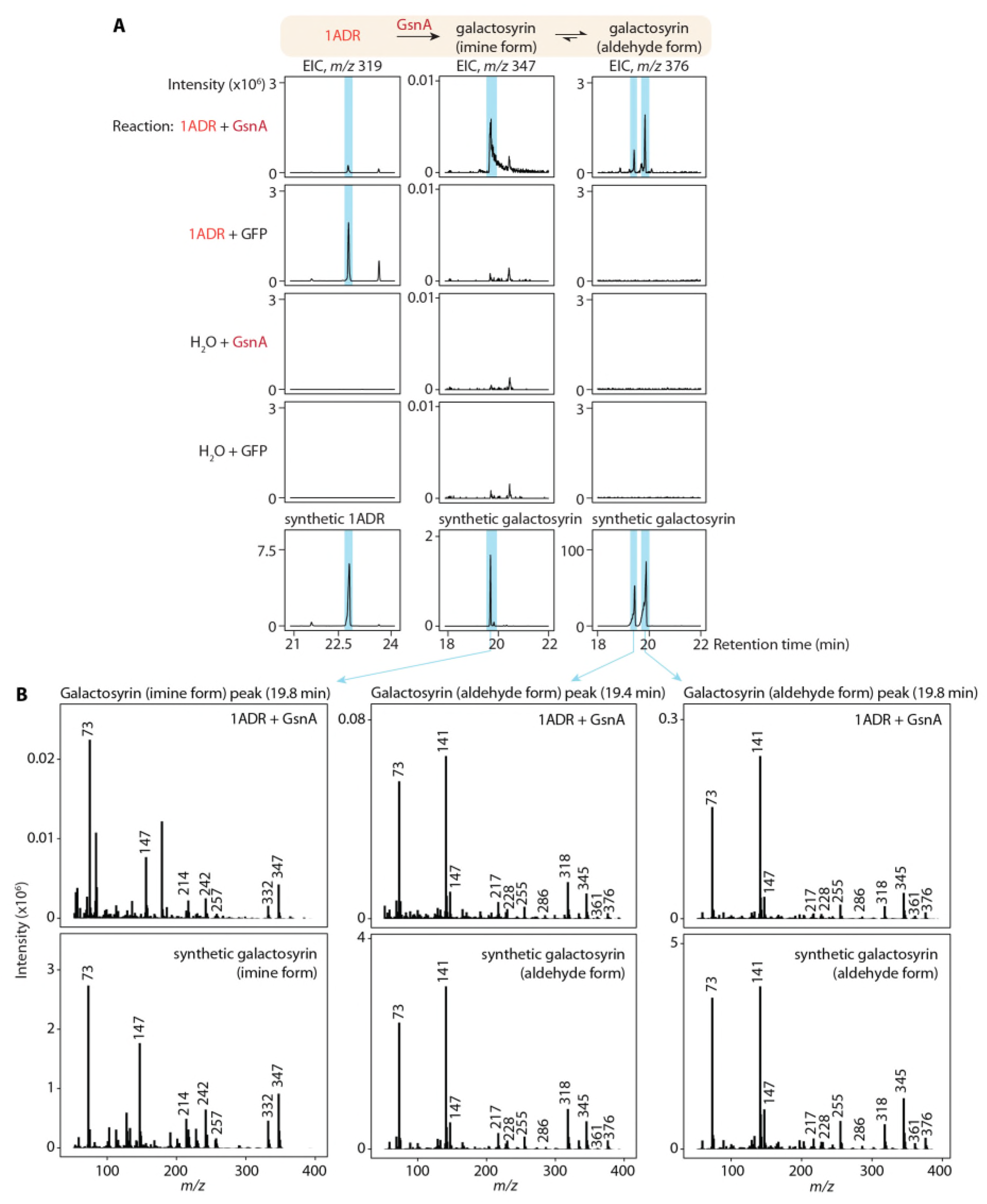
GsnA catalyses galactosyrin biosynthesis from 1ADR. **(A)** Summary diagram of the biosynthesis step is shown at the top. 1ADR is converted by GsnA into the galactosyrin imine form which can spontaneously convert into the aldehyde form. Extracted ion chromatograms (EIC) of mass signature corresponding to each compound (1ADR, galactosyrin imine and aldehyde form) are shown in each column. For each row, each mixture of enzyme and substrate (listed in red and orange on the left, GFP used as negative control) were mixed with the cofactors to reconstruct the biosynthesis, then soluble metabolite fractions were analysed by GC-MS. Blue stripes highlight peaks of the detected compounds, which are confirmed to match with synthetic standards shown in the bottom row. Galactosyrin (aldehyde form) was derivatized with methoxyamine, which yielded two stereoisomers detected as two peaks. **(B)** Mass spectra of the detected peaks with the matching spectra of synthetic standards shown underneath. Compound structures, derivatisation and mass annotations are shown in **Fig. S9**.

**Fig. S12.**
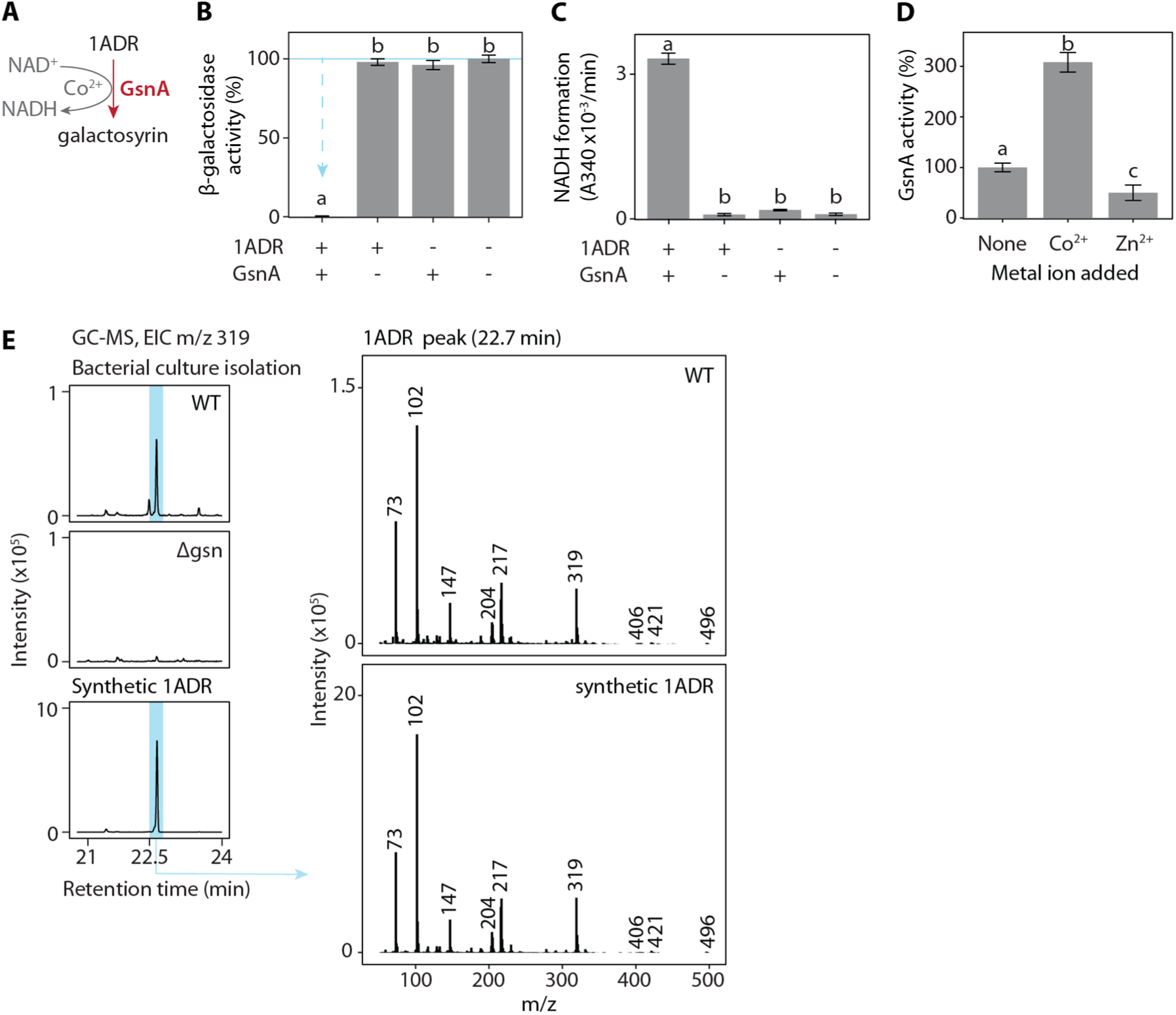
GsnA catalyses the production of galactosyrin using 1ADR as a substrate and NAD^+^ and Co^2+^ as cofactors. **(A)** Summary of the GsnA catalysed reaction. **(B)** Galactosyrin is produced by GsnA using 1ADR substrate. The enzyme GsnA (+) or GFP (-) and substrate 1ADR (+) or water (-) were mixed with cofactors, then the mixture was tested for galactosyrin formation in an enzyme activity assay with FDG substrate and LacZ enzyme. β-galactosidase activity is expressed as a percentage of the activity relative to the mean of no enzyme/substrate control. Arrows indicate inhibition. **(C)** NADH is formed from the oxidation of 1ADR catalysed by GsnA. The same reactions as (B) were monitored for NADH formation by measuring absorbance at 340 nm (A340). **(D)** GsnA has increased activity with Co^2+^ as a cofactor. Purified GsnA was pre-treated with EDTA to remove endogenously bound metal then used in 1ADR and NAD^+^ reaction mixture with or without Co^2+^ or Zn^2+^. The activity of GsnA was monitored with the rate of NADH formation and expressed as a percentage relative to no metal added sample. (B, C, D) Mean and standard deviation from 3 replicates are plotted. Different letters indicate different groups with statistically significant difference (P < 0.05) using one-way ANOVA and post-hoc Tukey HSD test. **(E)** 1ADR was also detected in bacterial culture. (Left) Extracted ion chromatogram (EIC) of *m/z* 319 from GC-MS analysis of cation-exchanged-isolated soluble metabolite extracts from cultures of galactosyrin-producing (WT) or galactosyrin-deficient (*`Δ`gsn*) strains. Blue stripes highlight the peak of 1ADR matching with the synthetic standard. (Right) Mass spectra of the detected 1ADR peaks showing identical expected masses. Compound structures, derivatisation and mass annotations are shown in **Fig. S9**.

**Fig. S13.**
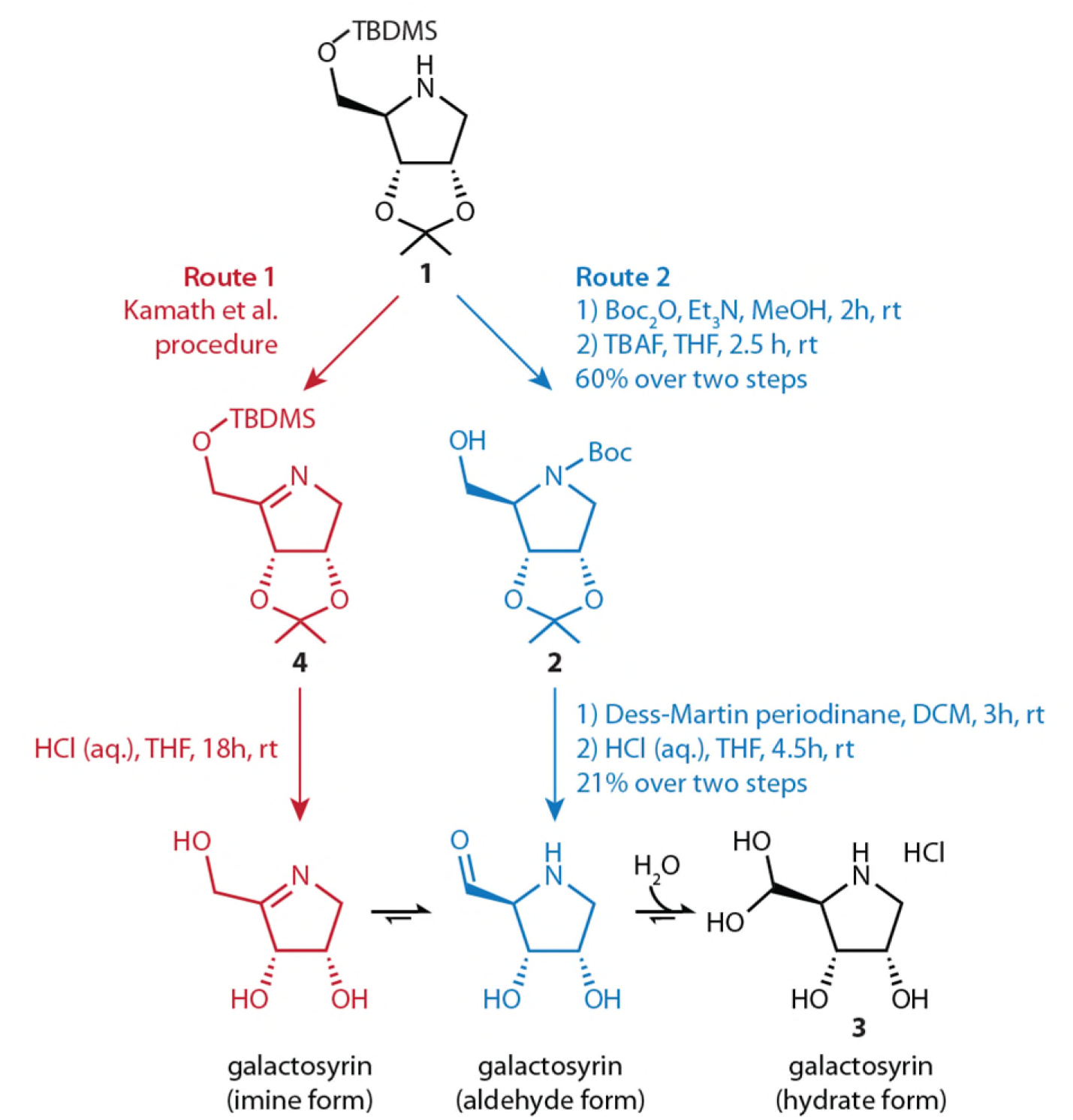
Chemical synthesis of galactosyrin. The hydrate form of galactosyrin can be obtained through two synthesis routes: route 1 via galactosyrin imine form using Kamath et al. procedure (*37*) and route 2 via galactosyrin aldehyde form.

**Fig. S14.**
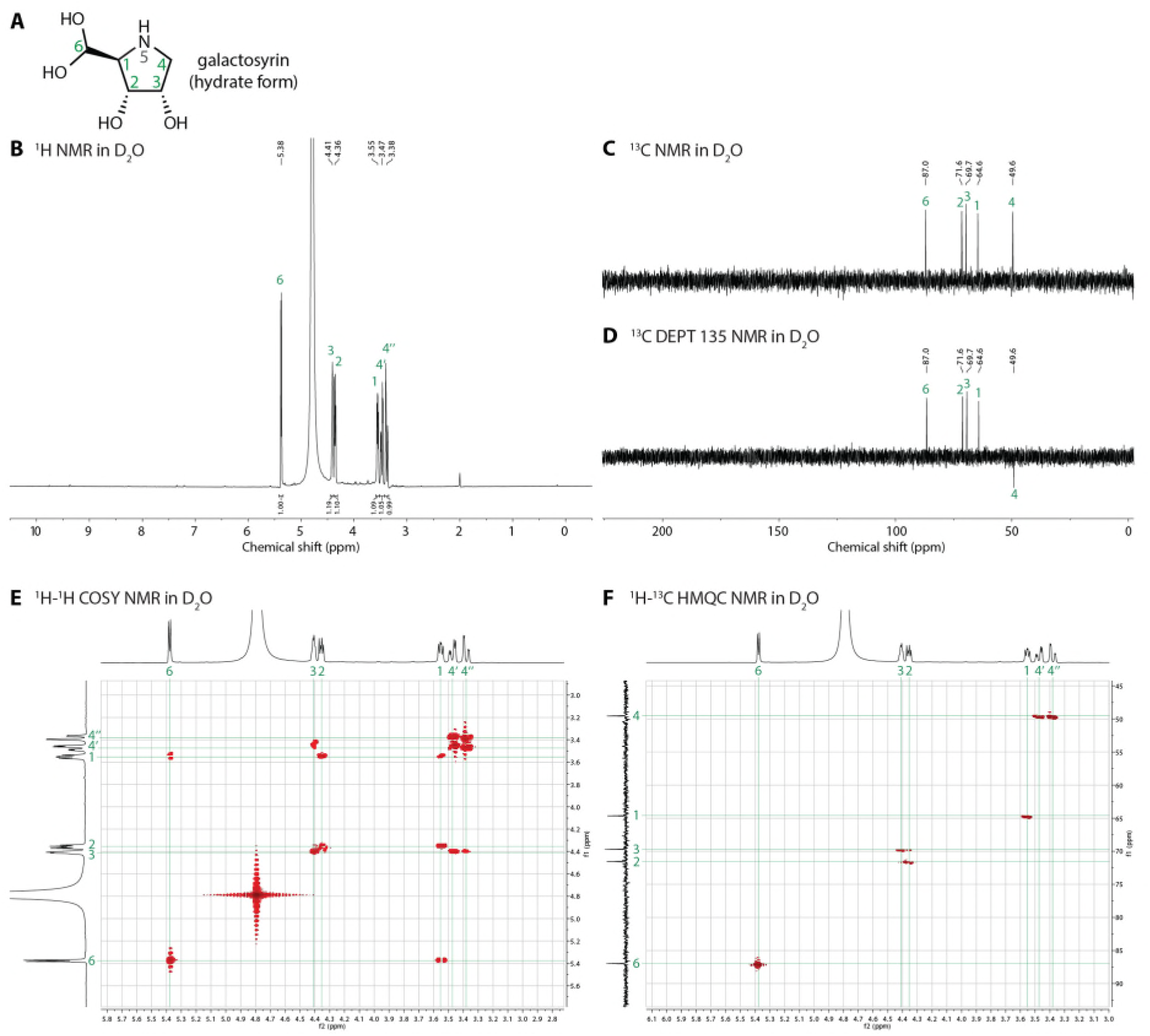
NMR spectra confirm the structure of the hydrate form of galactosyrin. **(A)** Structure of galactosyrin hydrate form with atom positions numbered (green) for assignment in NMR spectra. **(B)** ^1^H NMR. **(C)** ^13^C NMR. **(D)** ^13^C distortionless enhancement by polarization transfer (DEPT) 135 NMR. **(E)** ^1^H-^1^H homonuclear correlated spectroscopy (COSY) NMR. **(F)** ^1^H-^13^C heteronuclear multiple quantum correlation (HMQC) NMR.

**Fig. S15.**
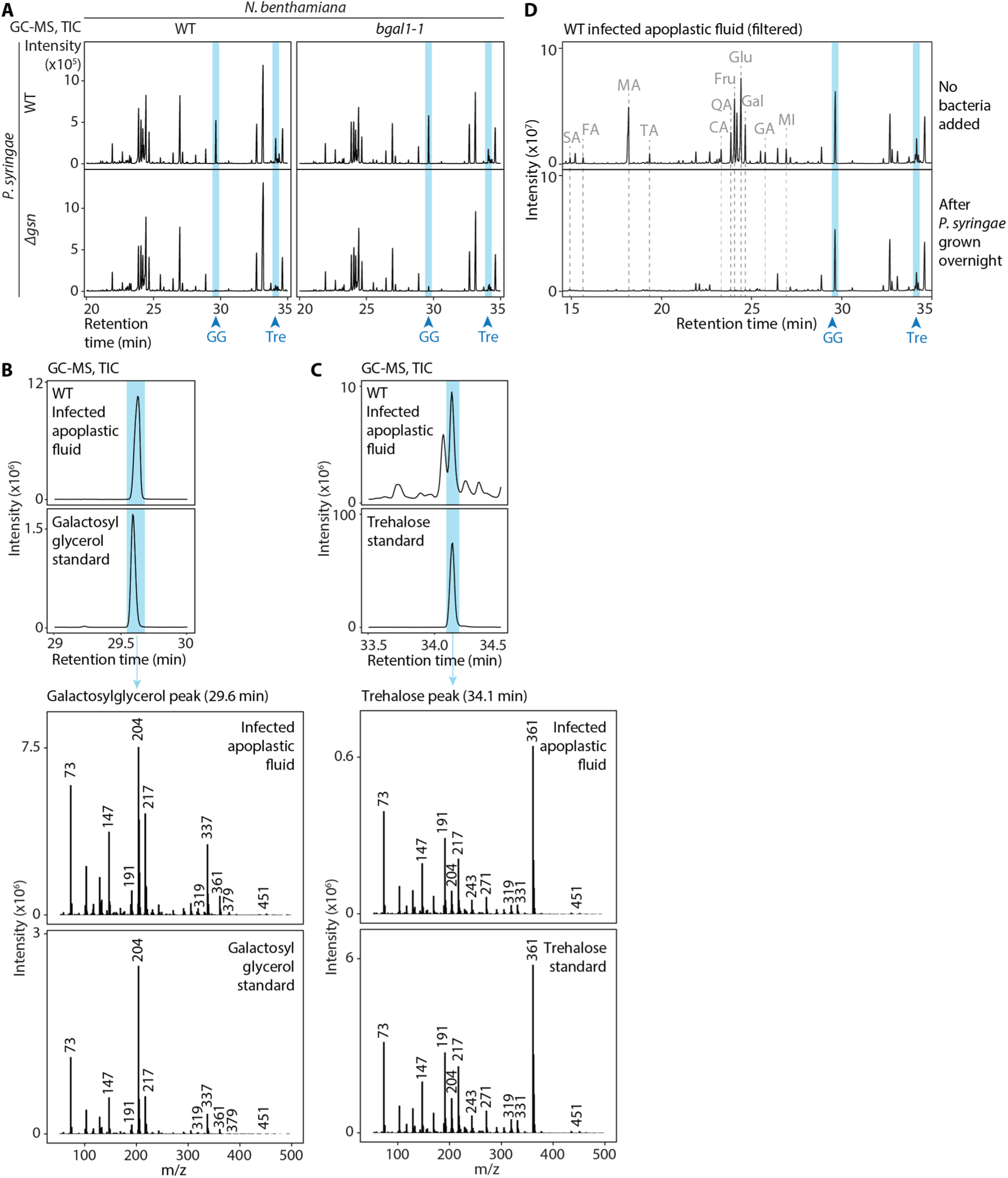
Galactosyrin production causes accumulation of galactosylglycerol and trehalose in the apoplast independent of BGAL1. **(A)** Total ion chromatograms (TIC) from GC-MS of soluble metabolites extracted from apoplastic fluids from *N. benthamiana* (wild-type (WT) or BGAL1 knockout mutant (*bgal1-1*) infected by galactosyrin-producing *P. syringae* (WT) or galactosyrin-deficient mutant (*`Δ`gsn*). Blue stripes highlight peaks of galactosylglycerol (GG) and trehalose (Tre). **(B)** Galactosylglycerol and **(C)** trehalose detection in the infected apoplastic fluid were confirmed by synthetic standards. (Top) Total ion chromatograms (TIC) from GC-MS of *P. syringae* (WT) infected apoplastic fluid. Blue stripes highlight peaks of the detected compounds, which are confirmed to match with synthetic standards shown underneath. (Bottom) Mass spectra of the detected peaks with the matching spectra of synthetic standards shown underneath. **(D)** Galactosylglycerol and trehalose are not preferred nutrient sources of *P. syringae*. Apoplastic fluid from *P. syringae* (WT)-infected *N. benthamiana* (WT) leaves was filter-sterilised then used as a culture medium for *P. syringae* WT with OD600 of 1 overnight at 28°C or incubated without the bacteria. Soluble metabolites were extracted from these samples, analysed with GC-MS and total ion chromatograms (TIC) are shown. Blue stripes highlight peaks of galactosylglycerol (GG) and trehalose (Tre), which were not depleted after *P. syringae* growth. Dashed lines highlight peaks of metabolites that were depleted, including sugars: fructose (Fru), glucose (Glu), galactose (Gal), organic acids: succinic (SA), fumaric (FA), malic (MA), threonic (TA), citric (CA), quinic (QA), gluconic (GA) acid and myo-inositol (MI).

**Fig. S16.**
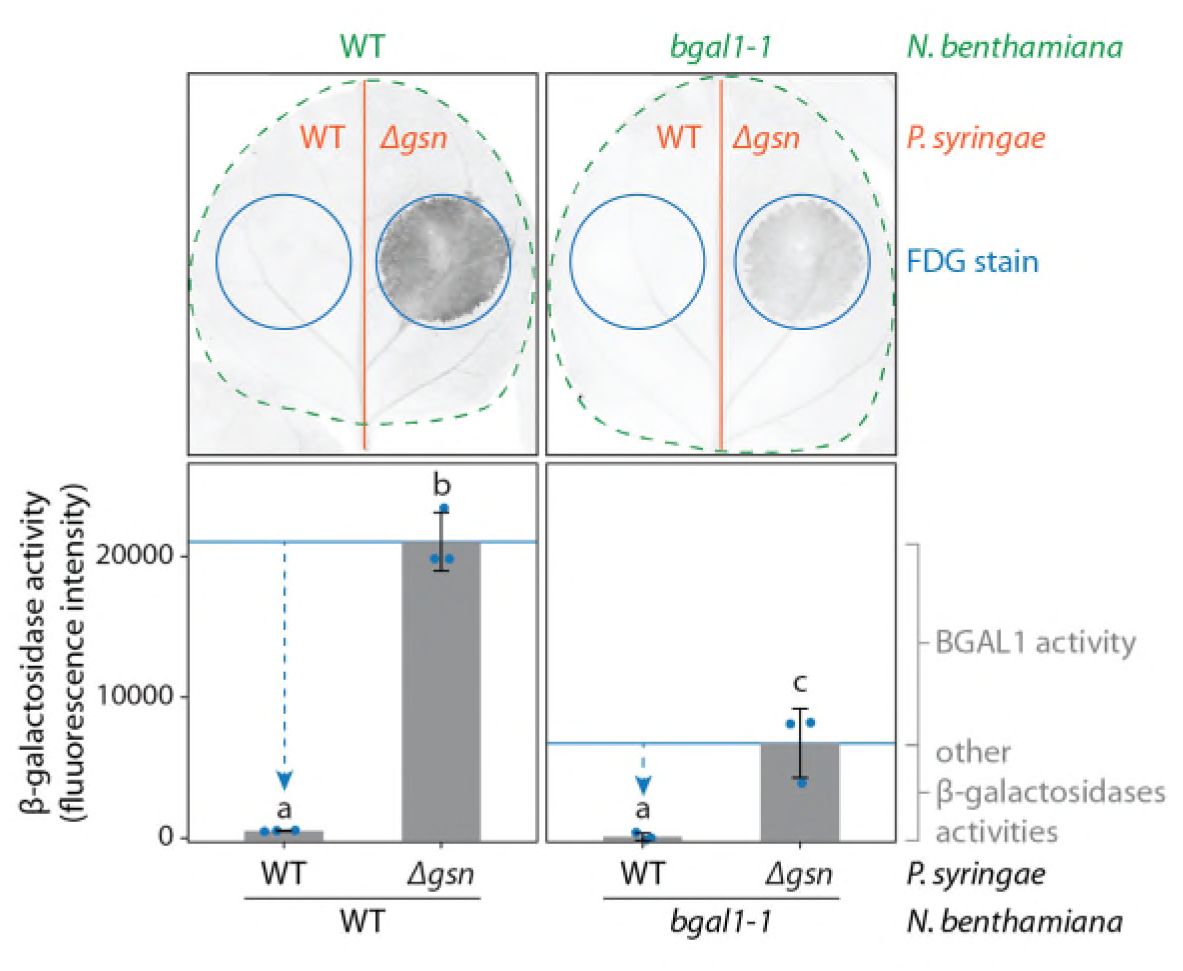
Galactosyrin inhibits not only BGAL1 but also other β-galactosidases in *N. benthamiana*. (Top) *N. benthamiana* leaves (WT or *bgal1-1* mutant lacking BGAL1 β-galactosidase) were infected with *P. syringae* (WT or the galactosyrin-deficient *`Δ`gsn* mutant) infiltrated into each half of each leaf. At three days post infection, the fluorogenic substrate FDG was infiltrated as spots into the leaves and β-galactosidase activity was measured by imaging the fluorescence signal of FDG cleavage product. (Bottom) β-galactosidase activity was quantified using integrated density of fluorescence signal from each spot. Error bars represent standard deviation from 3 replicates. Different letters indicate different groups with statistically significant difference (P < 0.01) using one-way ANOVA and post-hoc Tukey HSD test. Arrows highlight inhibition. β-galactosidase activity in the *bgal1-1* mutant was lower than WT plant since it lacks BGAL1 but activities of other β-galactosidases were still detectable. All of these β-galactosidase activities were suppressed by galactosyrin produced from WT *P. syringae*.

**Fig. S17.**
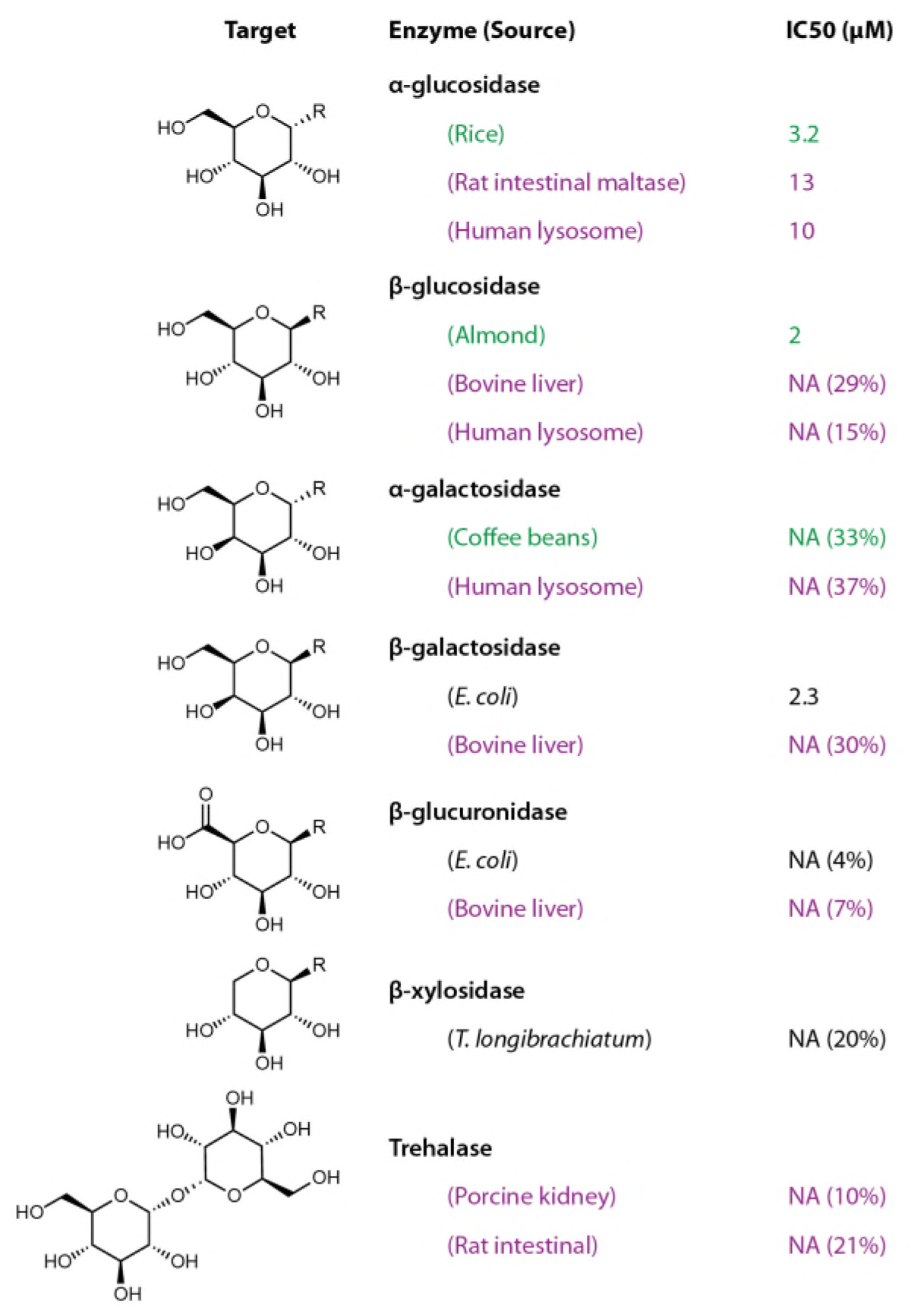
Galactosyrin inhibits different glycosidases. Half maximal inhibitory concentration (IC_50_) of galactosyrin inhibition of various glycosidases. NA denotes less than 50% inhibition at 100 μM galactosyrin, with numbers in brackets showing percent inhibition observed at 100 μM galactosyrin. Enzymes sources are colored: green:plants, purple:mammals and black:microbes.

**Table S1.**
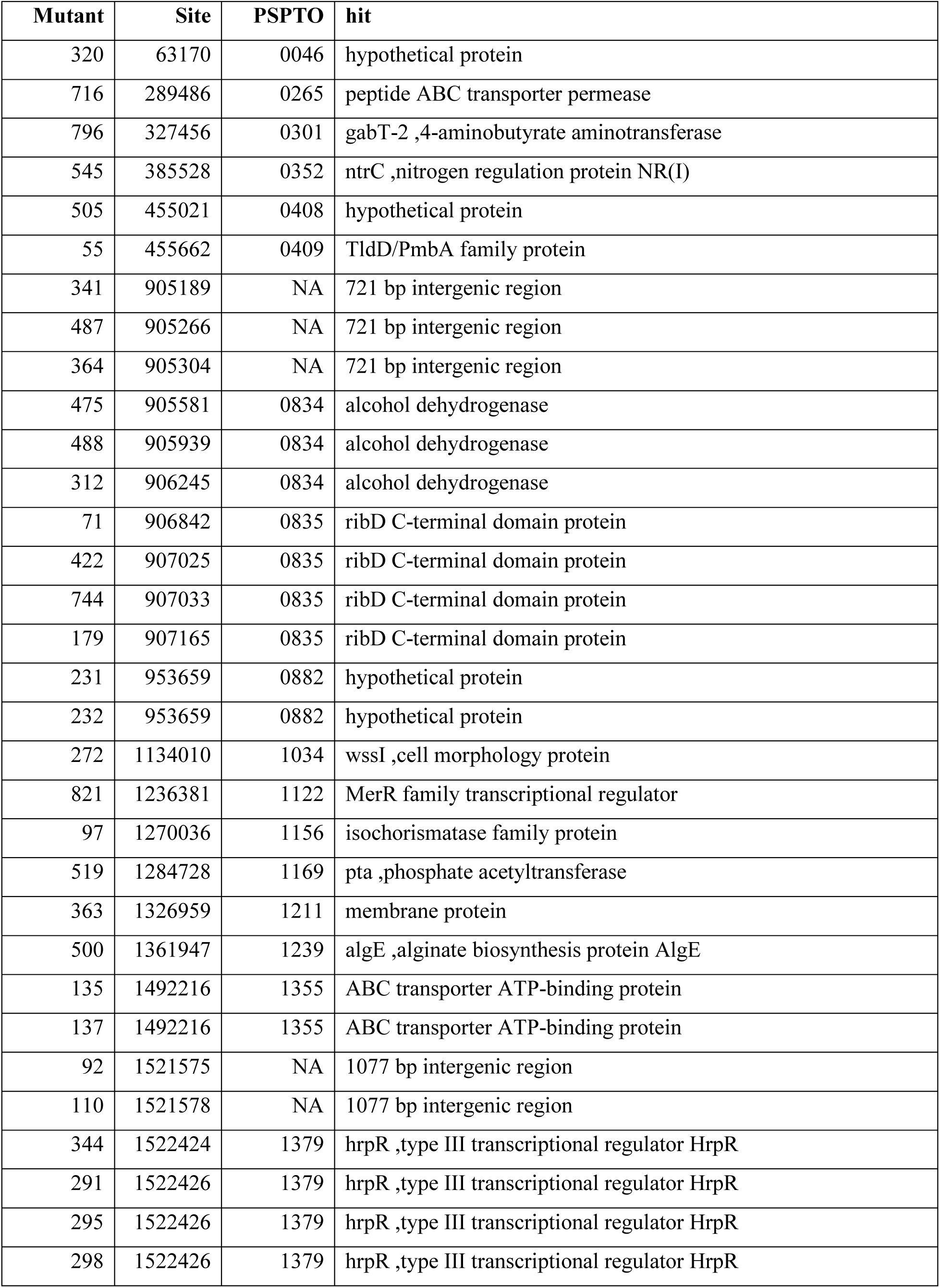
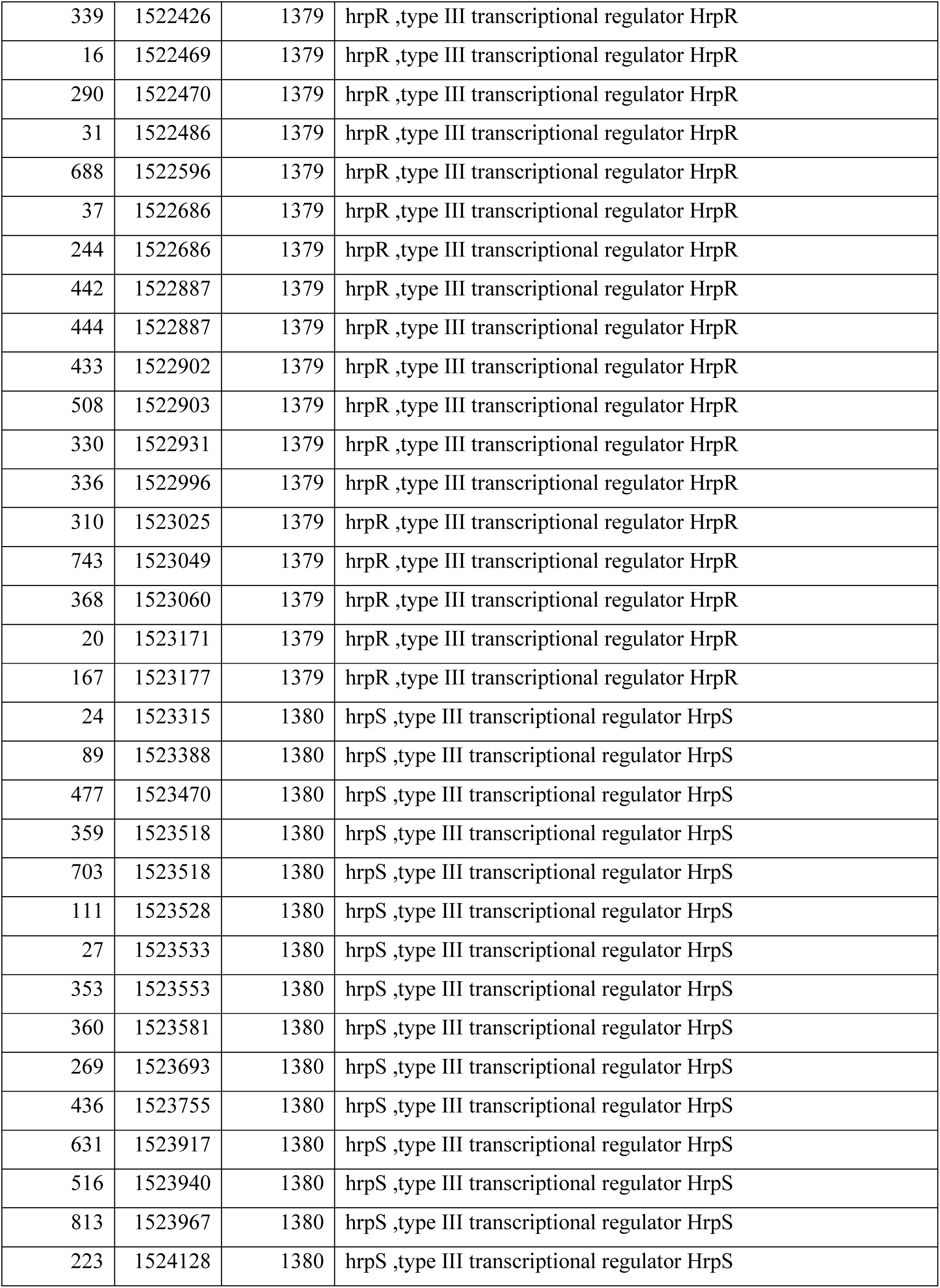
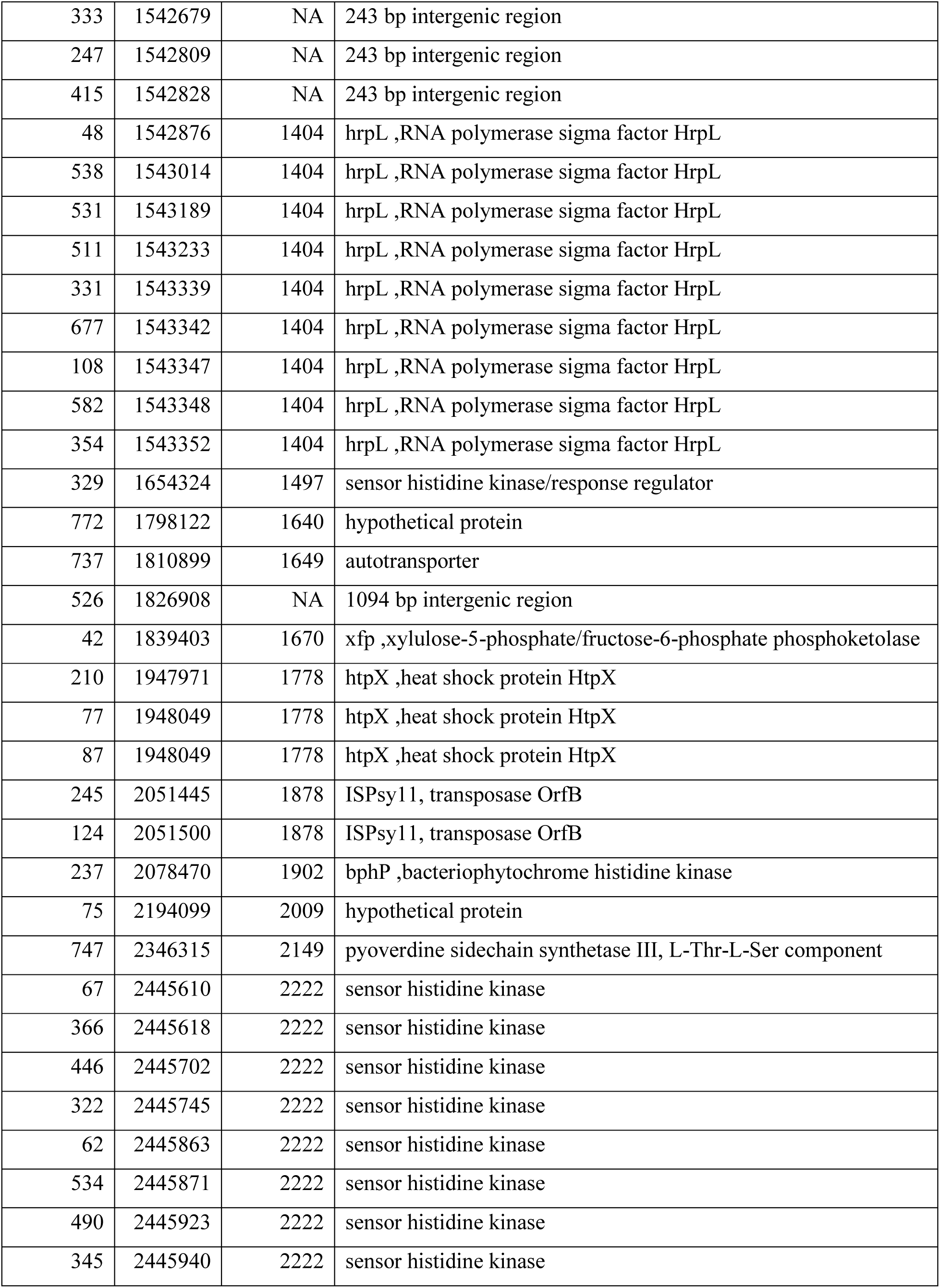
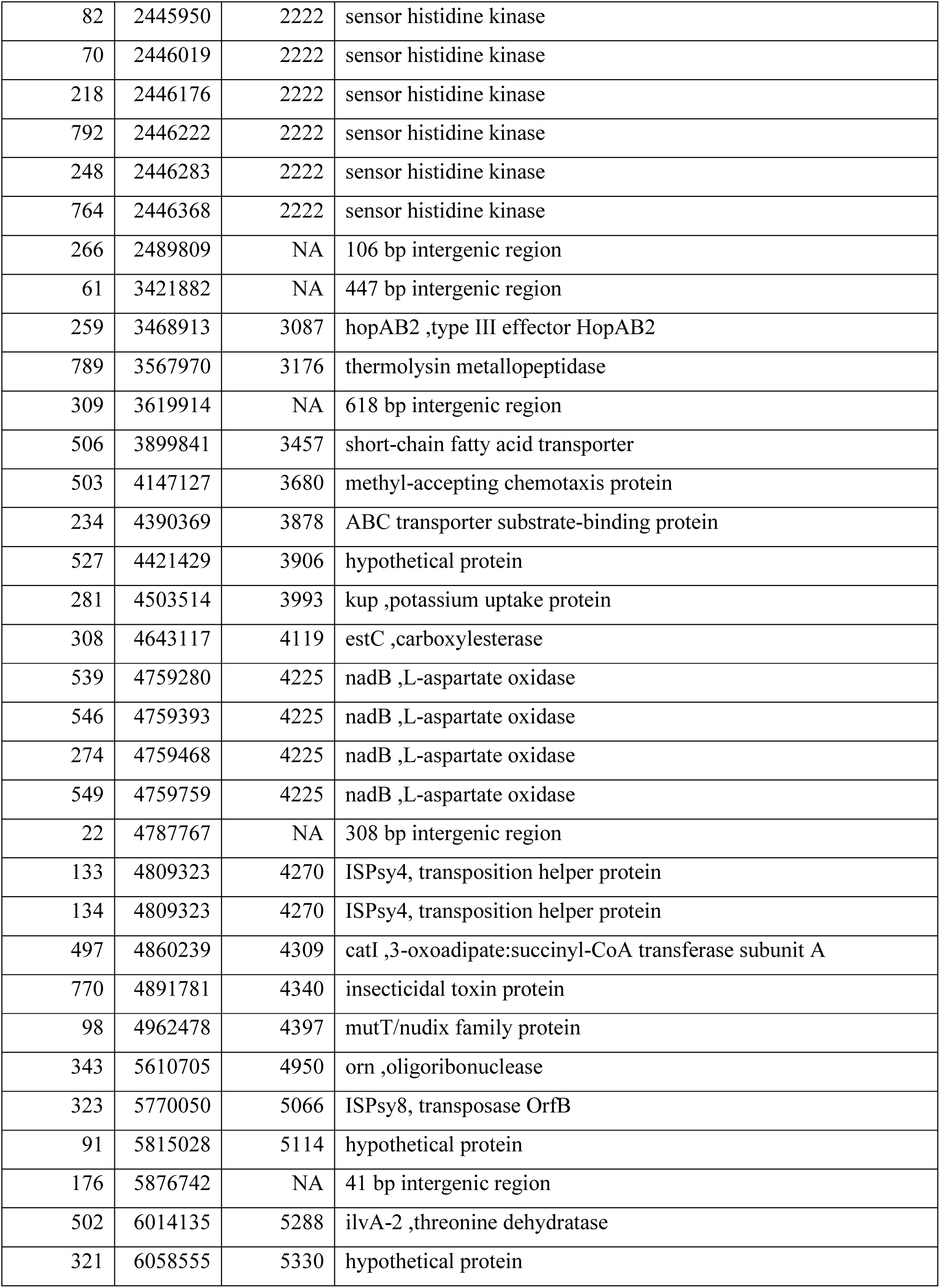
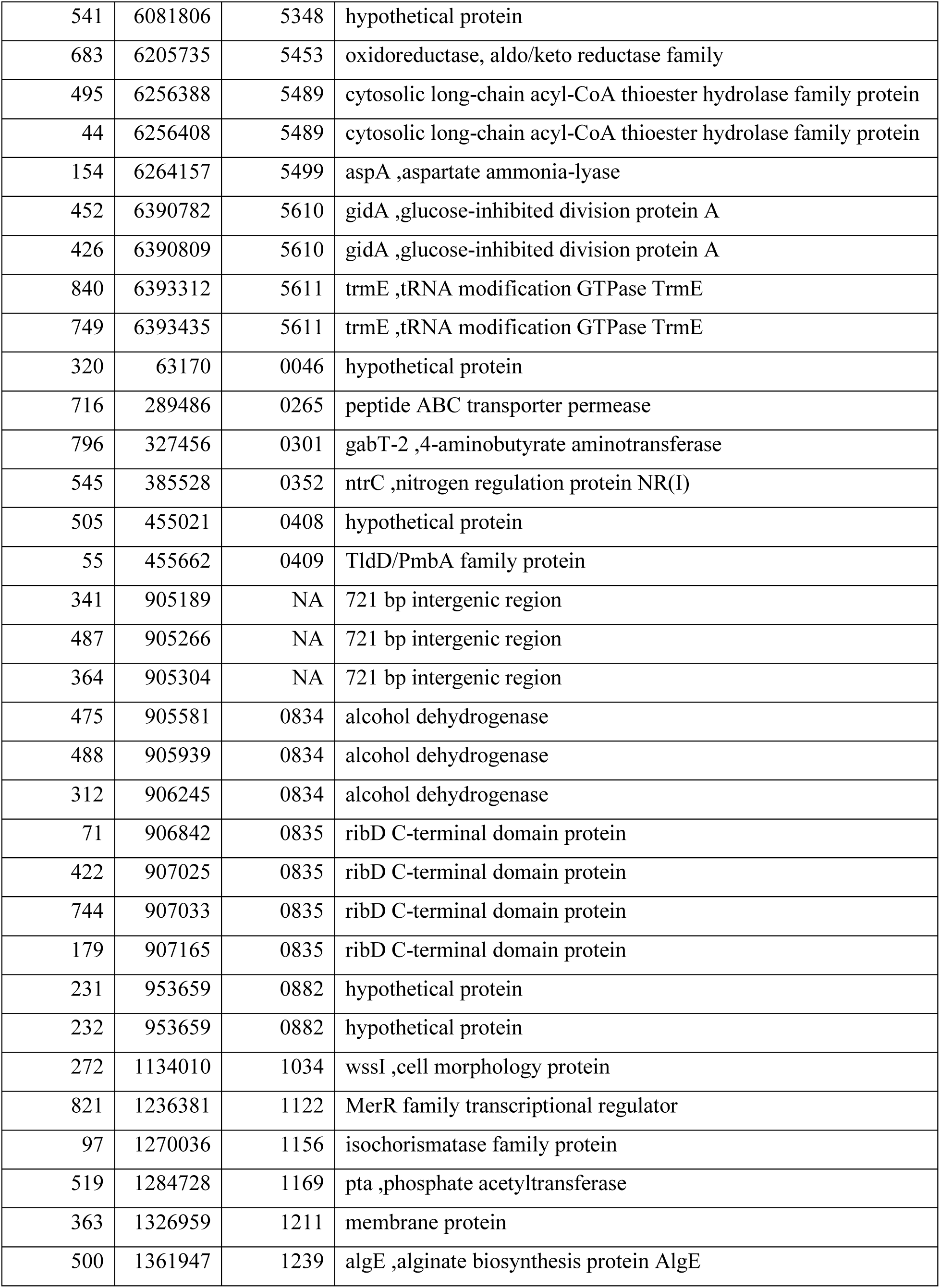
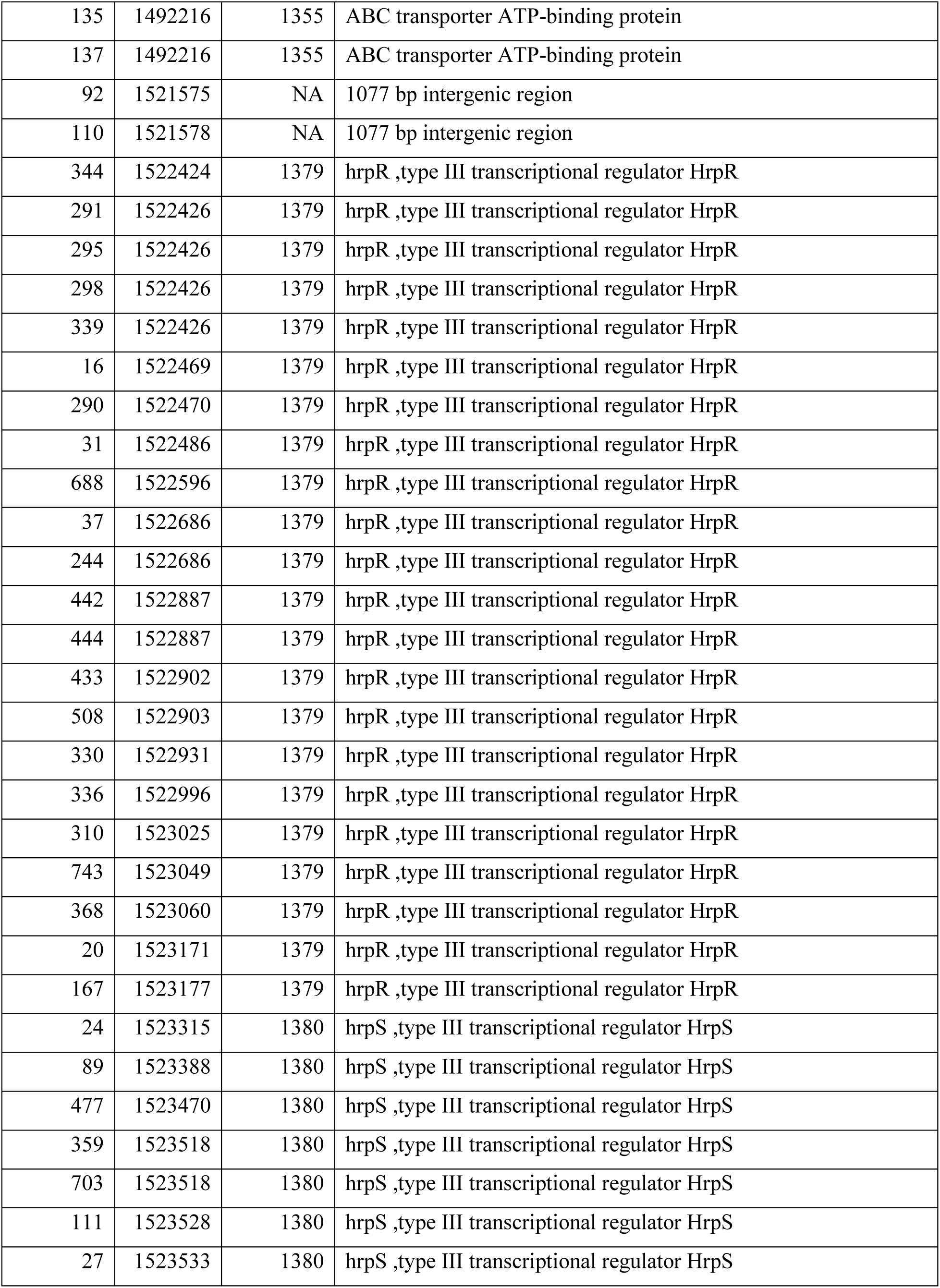
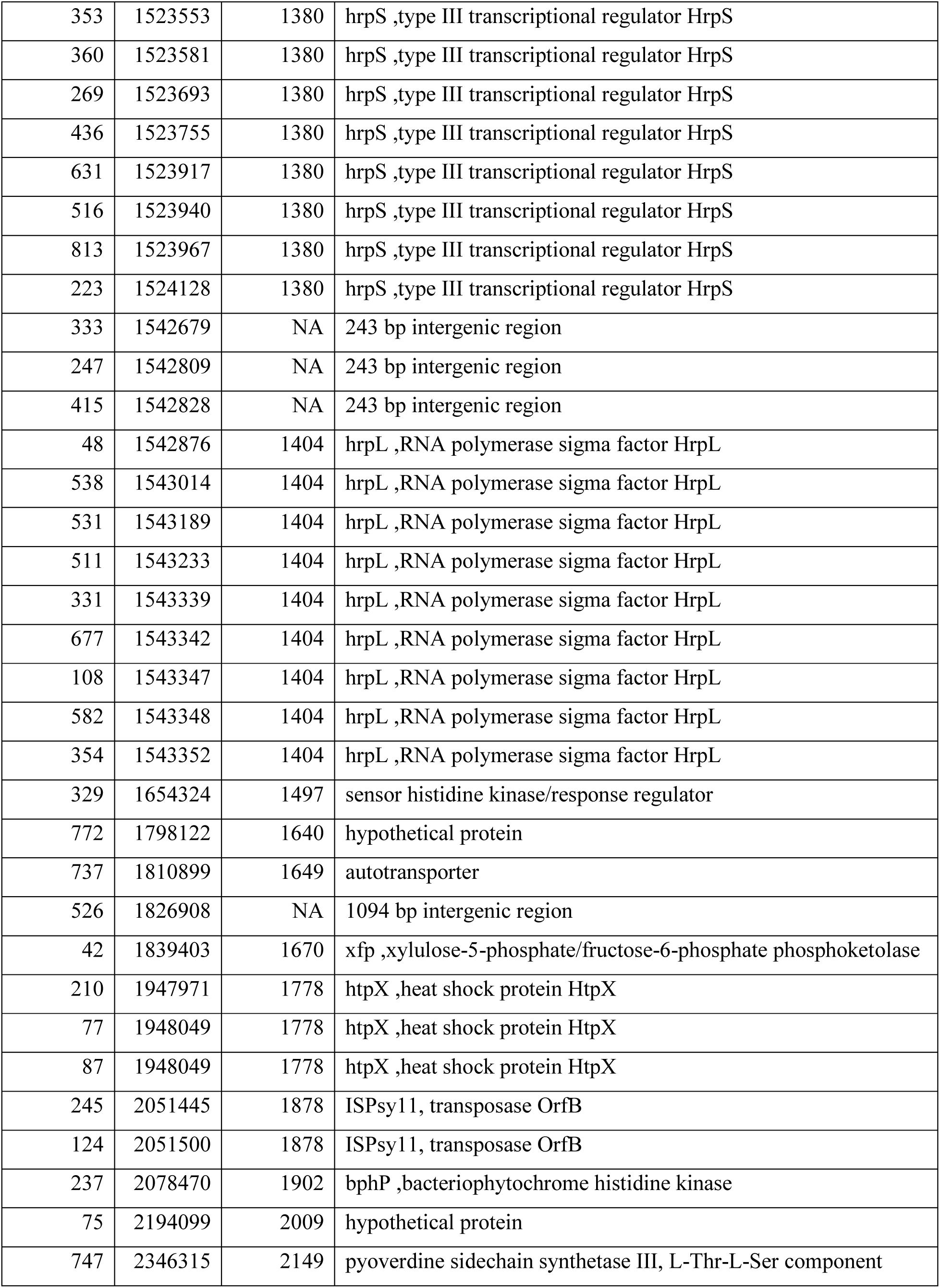
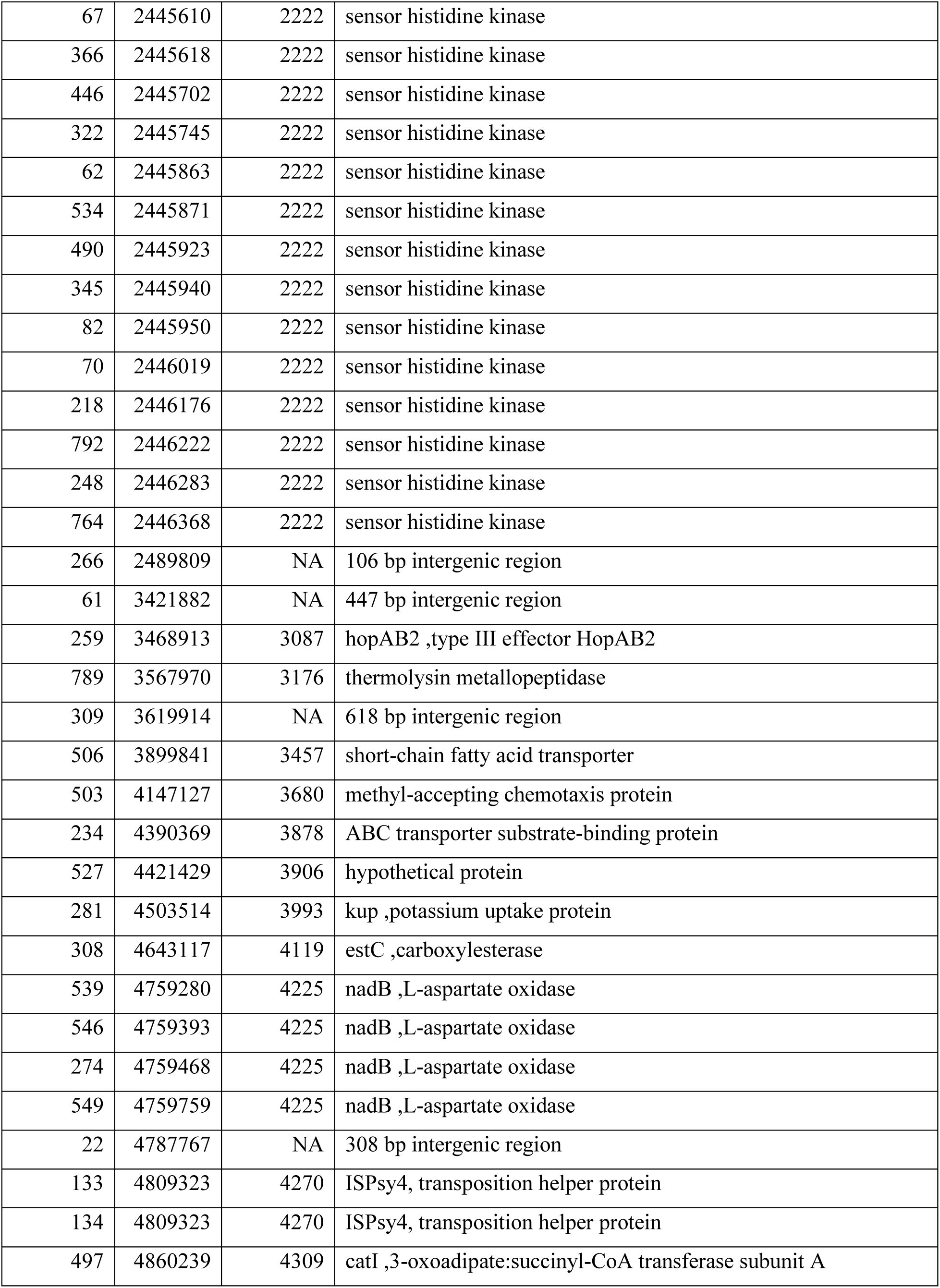
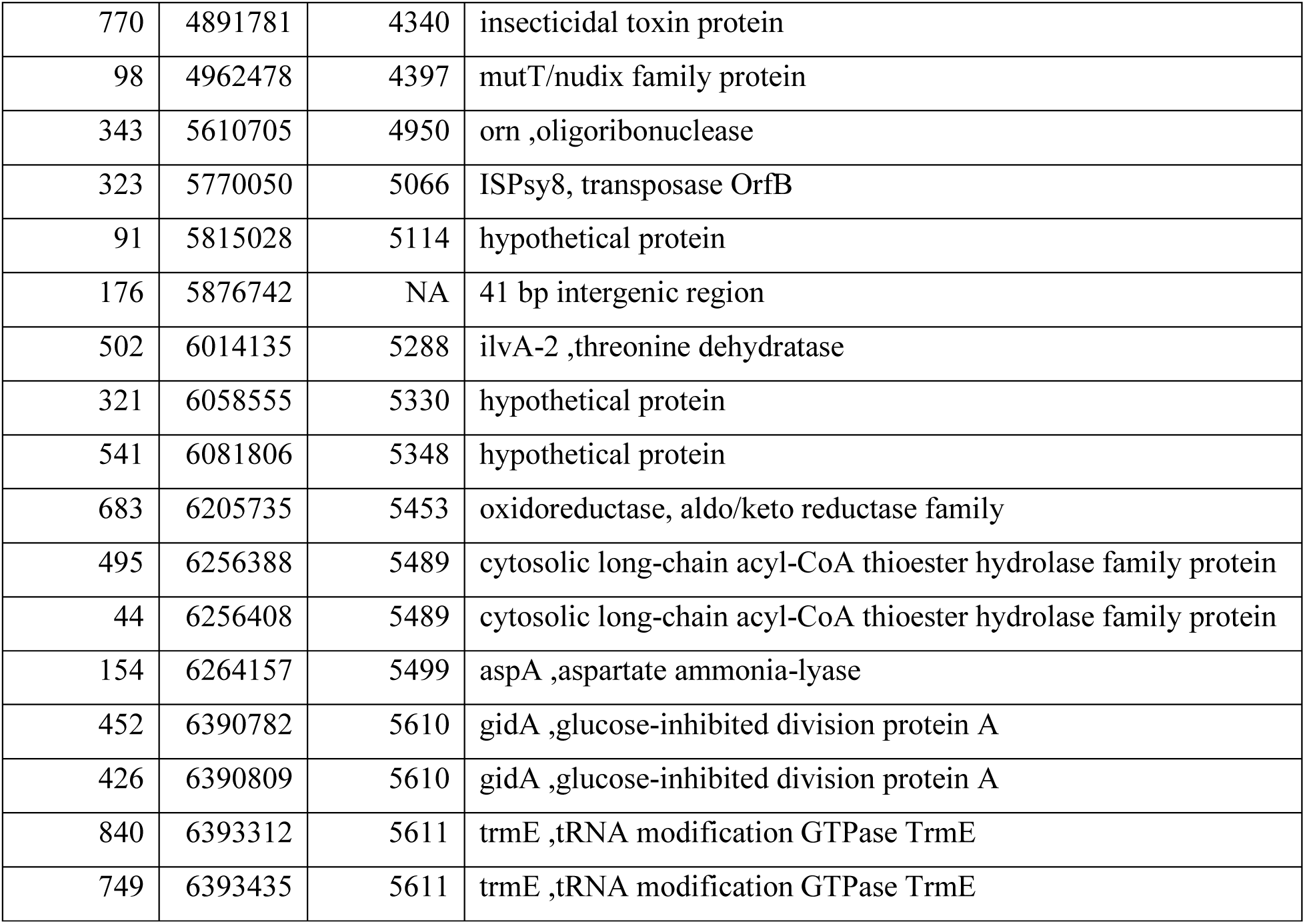
Tn5 insertion sites in galactosyrin-deficient mutants.

**Table S2.**
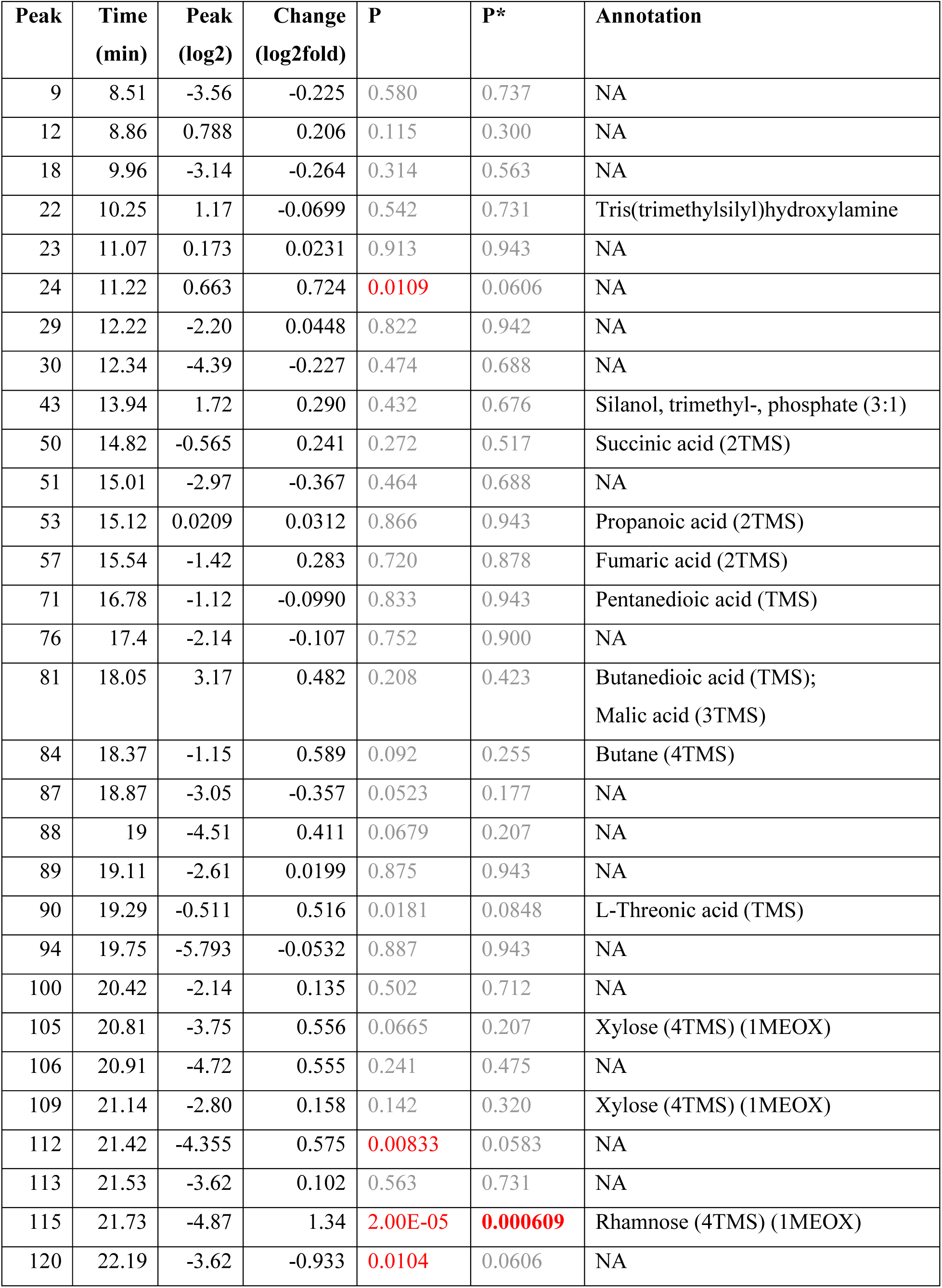
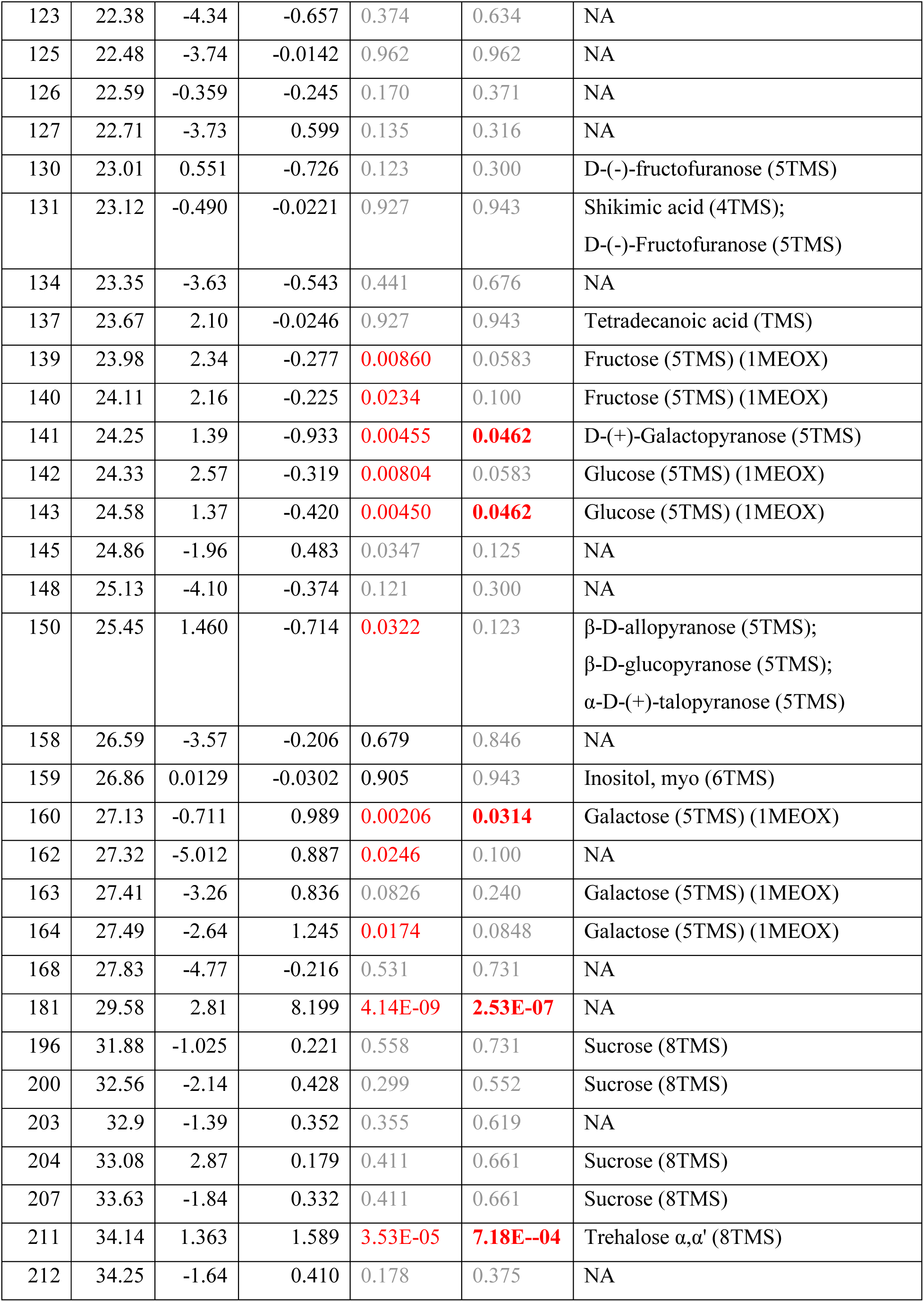
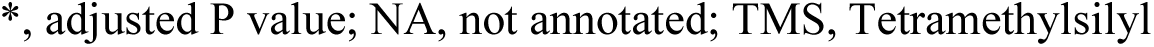
Metabolome of WT/Δgsn infection.

| Peak | Time<br>(min) | Peak<br>(log2) | Change<br>(log2fold) | P | P* | Annotation |
| --- | --- | --- | --- | --- | --- | --- |
| 9 | 8.51 | -3.56 | -0.225 | 0.580 | 0.737 | NA |
| 12 | 8.86 | 0.788 | 0.206 | 0.115 | 0.300 | NA |
| 18 | 9.96 | -3.14 | -0.264 | 0.314 | 0.563 | NA |
| 22 | 10.25 | 1.17 | -0.0699 | 0.542 | 0.731 | Tris(trimethylsilyl)hydroxylamine |
| 23 | 11.07 | 0.173 | 0.0231 | 0.913 | 0.943 | NA |
| 24 | 11.22 | 0.663 | 0.724 | 0.0109 | 0.0606 | NA |
| 29 | 12.22 | -2.20 | 0.0448 | 0.822 | 0.942 | NA |
| 30 | 12.34 | -4.39 | -0.227 | 0.474 | 0.688 | NA |
| 43 | 13.94 | 1.72 | 0.290 | 0.432 | 0.676 | Silanol, trimethyl-, phosphate (3:1) |
| 50 | 14.82 | -0.565 | 0.241 | 0.272 | 0.517 | Succinic acid (2TMS) |
| 51 | 15.01 | -2.97 | -0.367 | 0.464 | 0.688 | NA |
| 53 | 15.12 | 0.0209 | 0.0312 | 0.866 | 0.943 | Propanoic acid (2TMS) |
| 57 | 15.54 | -1.42 | 0.283 | 0.720 | 0.878 | Fumaric acid (2TMS) |
| 71 | 16.78 | -1.12 | -0.0990 | 0.833 | 0.943 | Pentanedioic acid (TMS) |
| 76 | 17.4 | -2.14 | -0.107 | 0.752 | 0.900 | NA |
| 81 | 18.05 | 3.17 | 0.482 | 0.208 | 0.423 | Butanedioic acid (TMS);<br>Malic acid (3TMS) |
| 84 | 18.37 | -1.15 | 0.589 | 0.092 | 0.255 | Butane (4TMS) |
| 87 | 18.87 | -3.05 | -0.357 | 0.0523 | 0.177 | NA |
| 88 | 19 | -4.51 | 0.411 | 0.0679 | 0.207 | NA |
| 89 | 19.11 | -2.61 | 0.0199 | 0.875 | 0.943 | NA |
| 90 | 19.29 | -0.511 | 0.516 | 0.0181 | 0.0848 | L-Threonic acid (TMS) |
| 94 | 19.75 | -5.793 | -0.0532 | 0.887 | 0.943 | NA |
| 100 | 20.42 | -2.14 | 0.135 | 0.502 | 0.712 | NA |
| 105 | 20.81 | -3.75 | 0.556 | 0.0665 | 0.207 | Xylose (4TMS) (1MEOX) |
| 106 | 20.91 | -4.72 | 0.555 | 0.241 | 0.475 | NA |
| 109 | 21.14 | -2.80 | 0.158 | 0.142 | 0.320 | Xylose (4TMS) (1MEOX) |
| 112 | 21.42 | -4.355 | 0.575 | 0.00833 | 0.0583 | NA |
| 113 | 21.53 | -3.62 | 0.102 | 0.563 | 0.731 | NA |
| 115 | 21.73 | -4.87 | 1.34 | 2.00E-05 | 0.000609 | Rhamnose (4TMS) (1MEOX) |
| 120 | 22.19 | -3.62 | -0.933 | 0.0104 | 0.0606 | NA |
| 123 | 22.38 | -4.34 | -0.657 | 0.374 | 0.634 | NA |
| 125 | 22.48 | -3.74 | -0.0142 | 0.962 | 0.962 | NA |
| 126 | 22.59 | -0.359 | -0.245 | 0.170 | 0.371 | NA |
| 127 | 22.71 | -3.73 | 0.599 | 0.135 | 0.316 | NA |
| 130 | 23.01 | 0.551 | -0.726 | 0.123 | 0.300 | D-(-)-fructofuranose (5TMS) |
| 131 | 23.12 | -0.490 | -0.0221 | 0.927 | 0.943 | Shikimic acid (4TMS);<br>D-(-)-Fructofuranose (5TMS) |
| 134 | 23.35 | -3.63 | -0.543 | 0.441 | 0.676 | NA |
| 137 | 23.67 | 2.10 | -0.0246 | 0.927 | 0.943 | Tetradecanoic acid (TMS) |
| 139 | 23.98 | 2.34 | -0.277 | 0.00860 | 0.0583 | Fructose (5TMS) (1MEOX) |
| 140 | 24.11 | 2.16 | -0.225 | 0.0234 | 0.100 | Fructose (5TMS) (1MEOX) |
| 141 | 24.25 | 1.39 | -0.933 | 0.00455 | 0.0462 | D-(+)-Galactopyranose (5TMS) |
| 142 | 24.33 | 2.57 | -0.319 | 0.00804 | 0.0583 | Glucose (5TMS) (1MEOX) |
| 143 | 24.58 | 1.37 | -0.420 | 0.00450 | 0.0462 | Glucose (5TMS) (1MEOX) |
| 145 | 24.86 | -1.96 | 0.483 | 0.0347 | 0.125 | NA |
| 148 | 25.13 | -4.10 | -0.374 | 0.121 | 0.300 | NA |
| 150 | 25.45 | 1.460 | -0.714 | 0.0322 | 0.123 | β-D-allopyranose (5TMS);<br>β-D-glucopyranose (5TMS);<br>α-D-(+)-talopyranose (5TMS) |
| 158 | 26.59 | -3.57 | -0.206 | 0.679 | 0.846 | NA |
| 159 | 26.86 | 0.0129 | -0.0302 | 0.905 | 0.943 | Inositol, myo (6TMS) |
| 160 | 27.13 | -0.711 | 0.989 | 0.00206 | 0.0314 | Galactose (5TMS) (1MEOX) |
| 162 | 27.32 | -5.012 | 0.887 | 0.0246 | 0.100 | NA |
| 163 | 27.41 | -3.26 | 0.836 | 0.0826 | 0.240 | Galactose (5TMS) (1MEOX) |
| 164 | 27.49 | -2.64 | 1.245 | 0.0174 | 0.0848 | Galactose (5TMS) (1MEOX) |
| 168 | 27.83 | -4.77 | -0.216 | 0.531 | 0.731 | NA |
| 181 | 29.58 | 2.81 | 8.199 | 4.14E-09 | 2.53E-07 | NA |
| 196 | 31.88 | -1.025 | 0.221 | 0.558 | 0.731 | Sucrose (8TMS) |
| 200 | 32.56 | -2.14 | 0.428 | 0.299 | 0.552 | Sucrose (8TMS) |
| 203 | 32.9 | -1.39 | 0.352 | 0.355 | 0.619 | NA |
| 204 | 33.08 | 2.87 | 0.179 | 0.411 | 0.661 | Sucrose (8TMS) |
| 207 | 33.63 | -1.84 | 0.332 | 0.411 | 0.661 | Sucrose (8TMS) |
| 211 | 34.14 | 1.363 | 1.589 | 3.53E-05 | 7.18E-04 | Trehalose $\alpha,\alpha'$ (8TMS) |
| 212 | 34.25 | -1.64 | 0.410 | 0.178 | 0.375 | NA |

\*, adjusted P value; NA, not annotated; TMS, Tetramethylsilyl

**Table S3.**
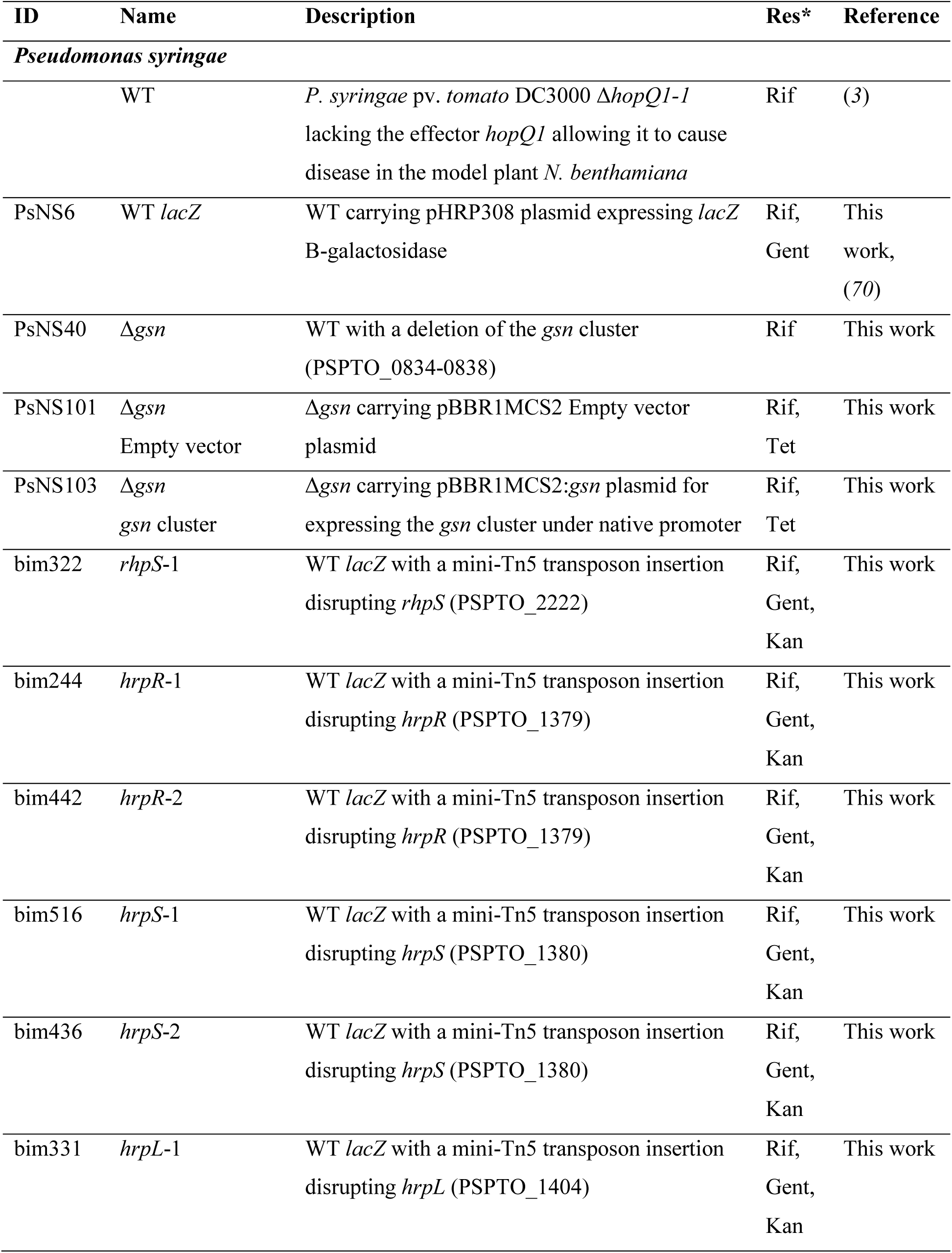
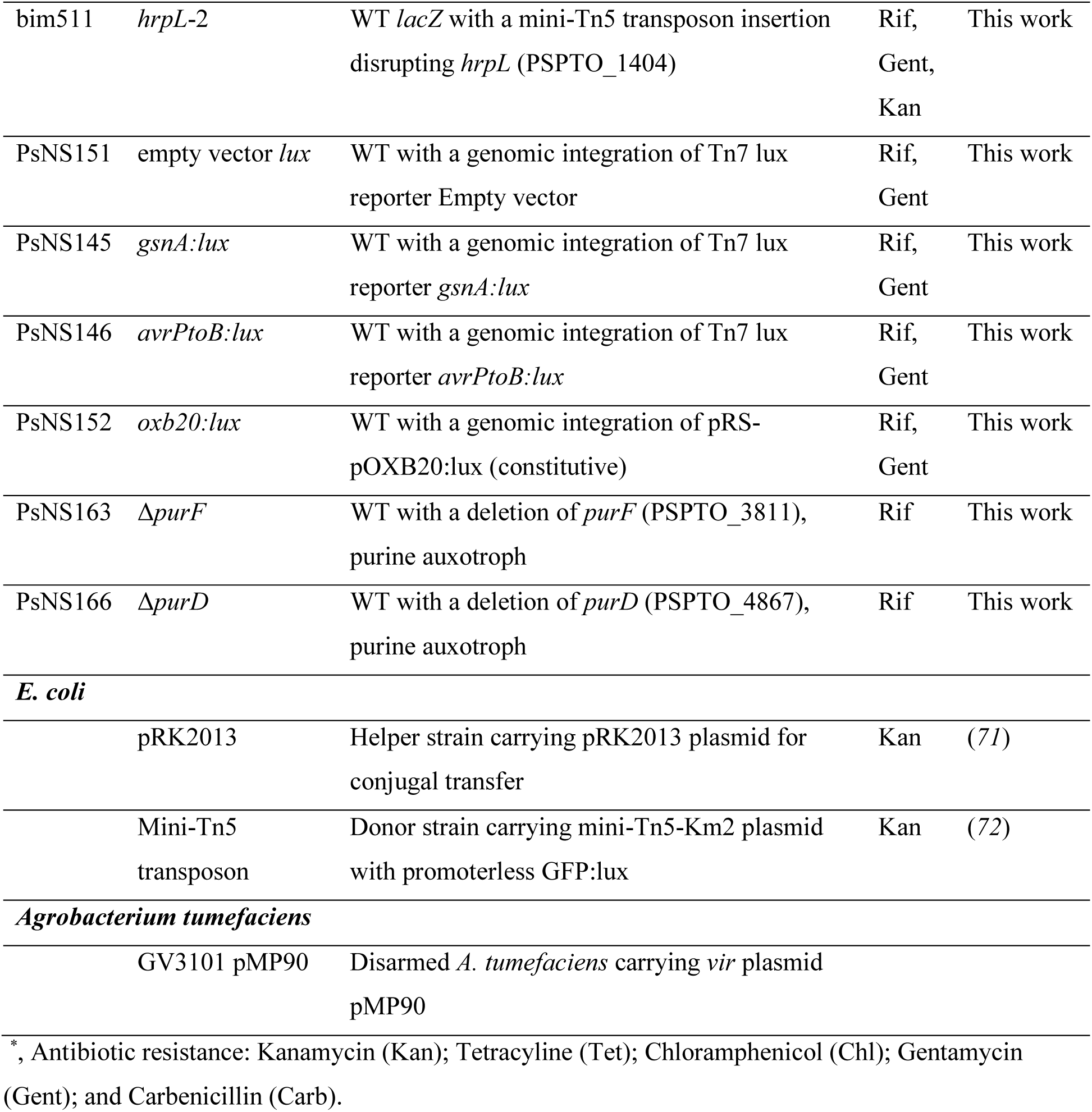
Bacterial strains used in this study.

| ID | Name | Description | Res* | Reference |
| --- | --- | --- | --- | --- |
| *Pseudomonas syringae* |  |  |  |  |
| | WT | *P. syringae* pv. *tomato* DC3000 Δ*hopQ1-1*<br>lacking the effector *hopQ1* allowing it to cause<br>disease in the model plant *N. benthamiana* | Rif | (3) |
| PsNS6 | WT *lacZ* | WT carrying pHRP308 plasmid expressing *lacZ*<br>B-galactosidase | Rif,<br>Gent | This<br>work,<br>(70) |
| PsNS40 | Δ*gsn* | WT with a deletion of the *gsn* cluster<br>(PSPTO_0834-0838) | Rif | This work |
| PsNS101 | Δ*gsn*<br>Empty vector | Δ*gsn* carrying pBBR1MCS2 Empty vector<br>plasmid | Rif,<br>Tet | This work |
| PsNS103 | Δ*gsn*<br>*gsn* cluster | Δ*gsn* carrying pBBR1MCS2:*gsn* plasmid for<br>expressing the *gsn* cluster under native promoter | Rif,<br>Tet | This work |
| bim322 | *rhpS-1* | WT *lacZ* with a mini-Tn5 transposon insertion<br>disrupting *rhpS* (PSPTO_2222) | Rif,<br>Gent,<br>Kan | This work |
| bim244 | *hrpR-1* | WT *lacZ* with a mini-Tn5 transposon insertion<br>disrupting *hrpR* (PSPTO_1379) | Rif,<br>Gent,<br>Kan | This work |
| bim442 | *hrpR-2* | WT *lacZ* with a mini-Tn5 transposon insertion<br>disrupting *hrpR* (PSPTO_1379) | Rif,<br>Gent,<br>Kan | This work |
| bim516 | *hrpS-1* | WT *lacZ* with a mini-Tn5 transposon insertion<br>disrupting *hrpS* (PSPTO_1380) | Rif,<br>Gent,<br>Kan | This work |
| bim436 | *hrpS-2* | WT *lacZ* with a mini-Tn5 transposon insertion<br>disrupting *hrpS* (PSPTO_1380) | Rif,<br>Gent,<br>Kan | This work |
| bim331 | *hrpL-1* | WT *lacZ* with a mini-Tn5 transposon insertion<br>disrupting *hrpL* (PSPTO_1404) | Rif,<br>Gent,<br>Kan | This work |
| bim511 | *hrpL-2* | WT *lacZ* with a mini-Tn5 transposon insertion disrupting *hrpL* (PSPTO_1404) | Rif,<br>Gent,<br>Kan | This work |
| PsNS151 | empty vector *lux* | WT with a genomic integration of Tn7 *lux* reporter Empty vector | Rif,<br>Gent | This work |
| PsNS145 | *gsnA:lux* | WT with a genomic integration of Tn7 *lux* reporter *gsnA:lux* | Rif,<br>Gent | This work |
| PsNS146 | *avrPtoB:lux* | WT with a genomic integration of Tn7 *lux* reporter *avrPtoB:lux* | Rif,<br>Gent | This work |
| PsNS152 | *oxb20:lux* | WT with a genomic integration of pRS-pOXB20:*lux* (constitutive) | Rif,<br>Gent | This work |
| PsNS163 | $\Delta purF$ | WT with a deletion of *purF* (PSPTO_3811), purine auxotroph | Rif | This work |
| PsNS166 | $\Delta purD$ | WT with a deletion of *purD* (PSPTO_4867), purine auxotroph | Rif | This work |
| ***E. coli*** |  |  |  |  |
|  | pRK2013 | Helper strain carrying pRK2013 plasmid for conjugal transfer | Kan | (71) |
|  | Mini-Tn5 transposon | Donor strain carrying mini-Tn5-Km2 plasmid with promoterless GFP:*lux* | Kan | (72) |
| ***Agrobacterium tumefaciens*** |  |  |  |  |
|  | GV3101 pMP90 | Disarmed *A. tumefaciens* carrying *vir* plasmid pMP90 |  |  |

\*, Antibiotic resistance: Kanamycin (Kan); Tetracycline (Tet); Chloramphenicol (Chl); Gentamycin (Gent); and Carbenicillin (Carb).

**Table S4.** Plasmids used in this study.

| **ID** | **Name** | **Description** | **Res<sup>\*</sup>** | **Reference** |
| --- | --- | --- | --- | --- |
| pNS38 | pK18mobsacB<br>*Δgsn* | used for generation of *Δgsn* deletion mutant | *Kan* | This work |
| pNS172 | pK18mobsacB<br>*ΔpurF* | used for generation of *ΔpurF* deletion mutant | *Kan* | This work |
| pNS173 | pK18mobsacB<br>*ΔpurD* | used for generation of *ΔpurD* deletion mutant | *Kan* | This work |
| pNS35 | pBBR1MCS2:*gsn* | plasmid with the *gsn* cluster under native promoter | *Tet* | This work |
| pNS99 | pBBR1MCS3:*gsn* | plasmid with the *gsn* cluster under a constitutive Lac promoter | *Tet* | This work |
| pNS161 | Tn7 lux reporter<br>Empty vector | Tn7 transposon construct with no promoter:*luxCDABE*, derived from pRS-pOXB20:*lux* by removing pOXB20 promoter and inserting T0T1 terminator upstream to prevent unwanted transcription | *Gent* | This work |
| pNS163 | Tn7 lux reporter<br>*gsnA:lux* | Tn7 transposon construct with *gsnA* promoter:*luxCDABE* fusion for transcriptional activity reporter | *Gent* | This work |
| pNS164 | Tn7 lux reporter<br>*avrPtoB:lux* | Tn7 transposon construct with *avrPtoB* promoter:*luxCDABE* fusion for transcriptional activity reporter | *Gent* | This work |
| pNS141 | pET-28b:LacZHis | pET-28b expressing LacZ with a C-terminal 6xHistidine tag, in T7 Express lysY/Iq expression strain (New England Biolabs) | *Kan* | This work |
| pNS177 | pET-28b:PurF | pET-28b expressing PurF with a C-terminal Strep-tagII, in Lemo21(DE3) expression strain (New England Biolabs) | *Kan*,<br>*Chl* | This work |
| pNS70 | pET-28b:GsnA | pET-28b expressing GsnA with a C-terminal Strep-tagII, in T7 Express lysY/Iq expression strain (New England Biolabs) | *Kan* | This work |
| pNS155 | pET-28b:GsnB | pET-28b expressing GsnB with a C-terminal Strep-tagII, in Lemo21(DE3) expression strain (New England Biolabs) | *Kan*,<br>*Chl* | This work |
| pNS77 | pET-28b:GsnC | pET-28b expressing GsnC with a C-terminal Strep-tagII, in T7 Express lysY/Iq expression strain (New England Biolabs) | *Kan* | This work |
| pNS153 | pET-28b:sfGFP | pET-28b expressing sfGFP with a C-terminal Strep-tagII, in T7 Express lysY/Iq expression strain (New England Biolabs) | *Kan* | This work |
| pRK2013 | pRK2013 | Helper strain carrying pRK2013 plasmid for conjugal transfer | *Kan* | (71) |
| pK18mobsacB | pK18mobsacB<br>Empty vector | pK18mobsacB plasmid used for generation of deletion mutants | *Kan* | (42) |
| pBBR1MCS2 | pBBR1MCS2<br>Empty vector | pBBR1MCS2 plasmid used for gene expression | *Tet* | (43) |
| pBBR1MCS3 | pBBR1MCS3<br>Empty vector | pBBR1MCS3 plasmid used for gene expression | *Tet* | (43) |
| pUXBF13 | pUXBF13 | pUXBF13 plasmid for Tn7 transposon integration | *Carb* | (73) |
| OXB20:lux | pRS-pOXB20:lux | Tn7 transposon construct with pOXB20:*luxCDABE* constitutive expression | *Gent* | (44) |
| pET-28b | pET-28b Empty vector | pET-28b plasmid for recombinant protein production under IPTG-inducible T7 promoter | *Kan* | Novagen |
| pBK26 | 35S::BGAL1 | Binary vector for overexpression of BGAL1 by agroinfiltration | *Kan* | (2) |

\*, Antibiotic resistance: Kanamycin (Kan); Tetracycline (Tet); Chloramphenicol (Chl); Gentamycin (Gent); and Carbenicillin (Carb).

**Table S5.**
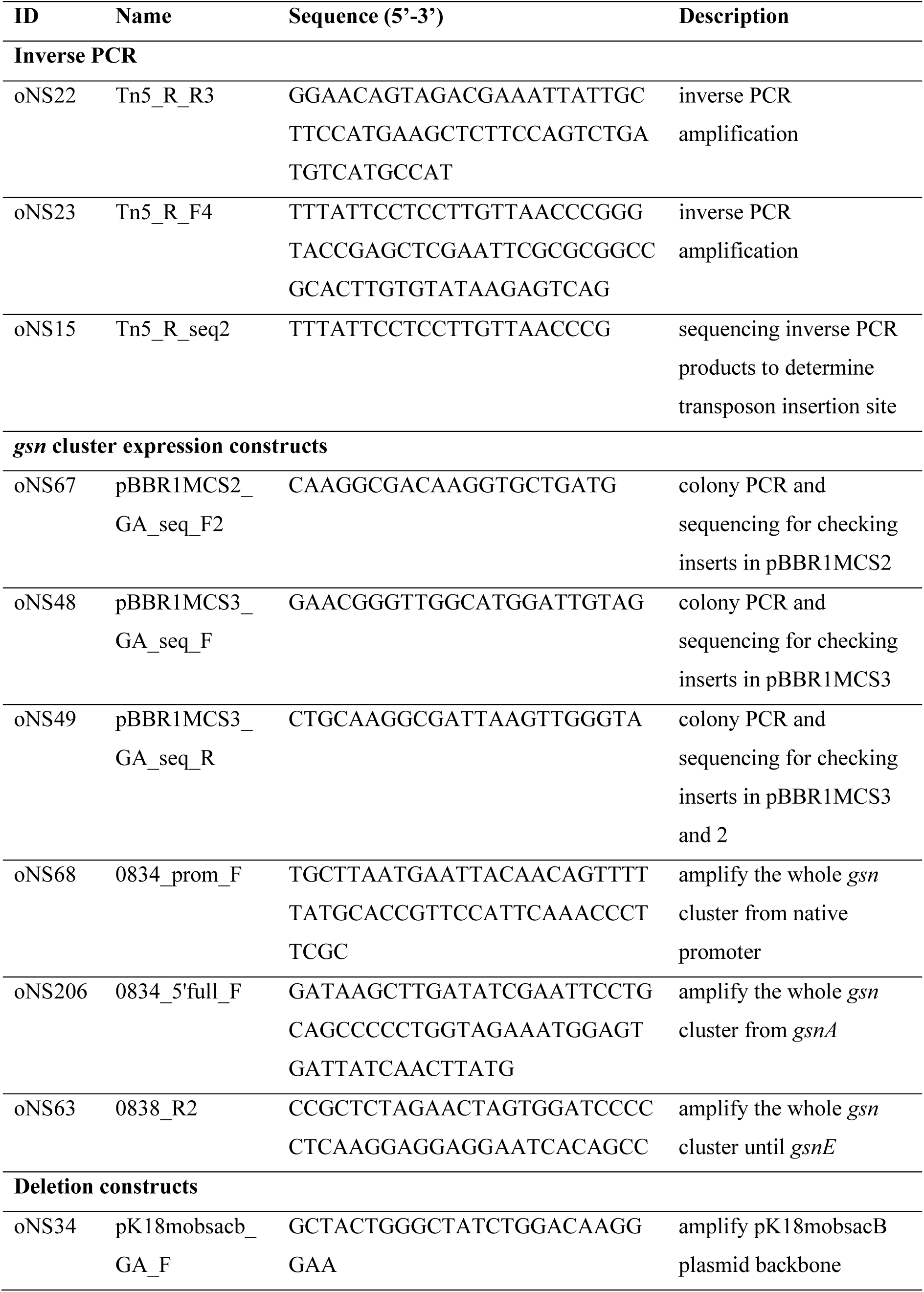
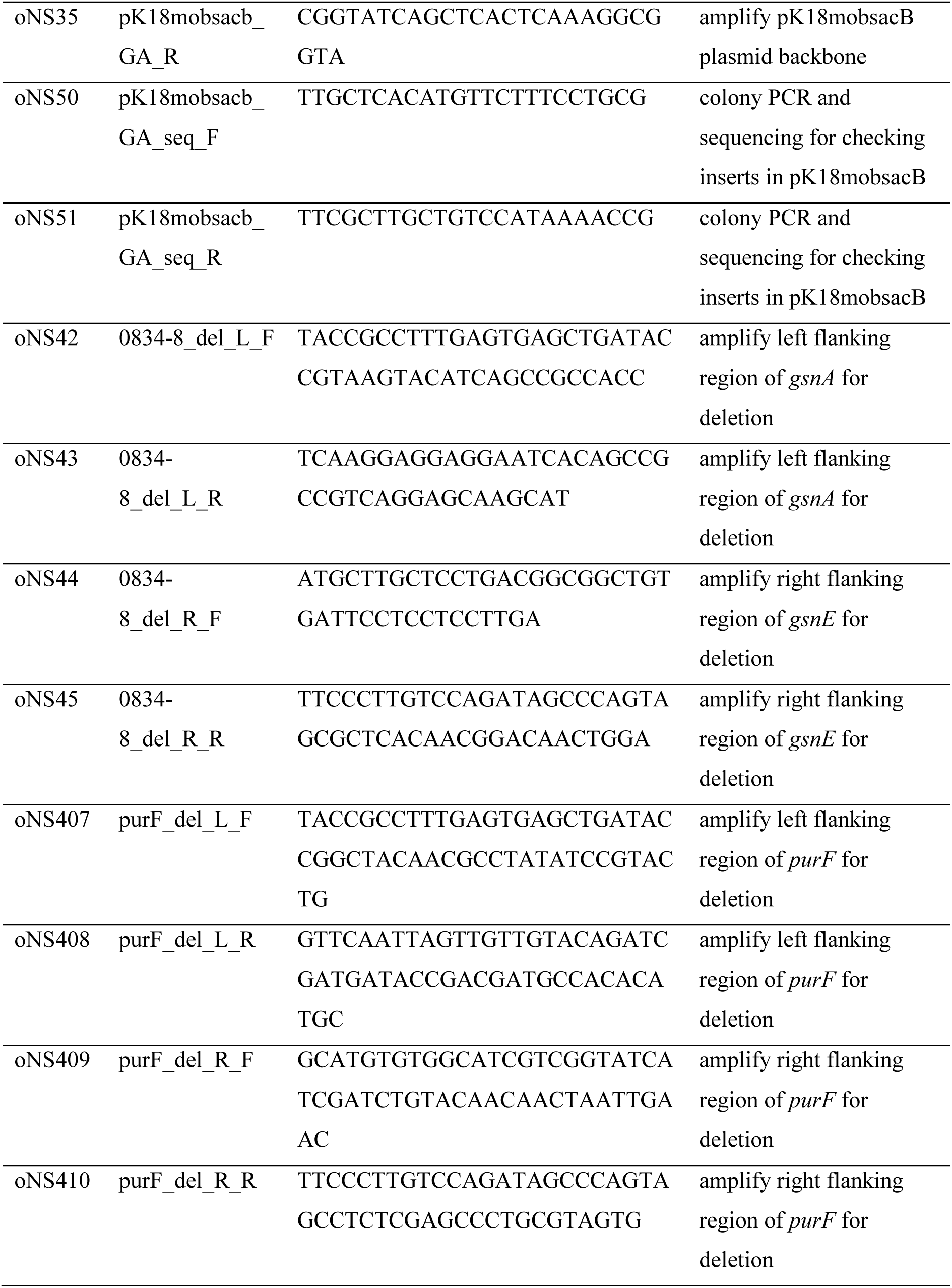
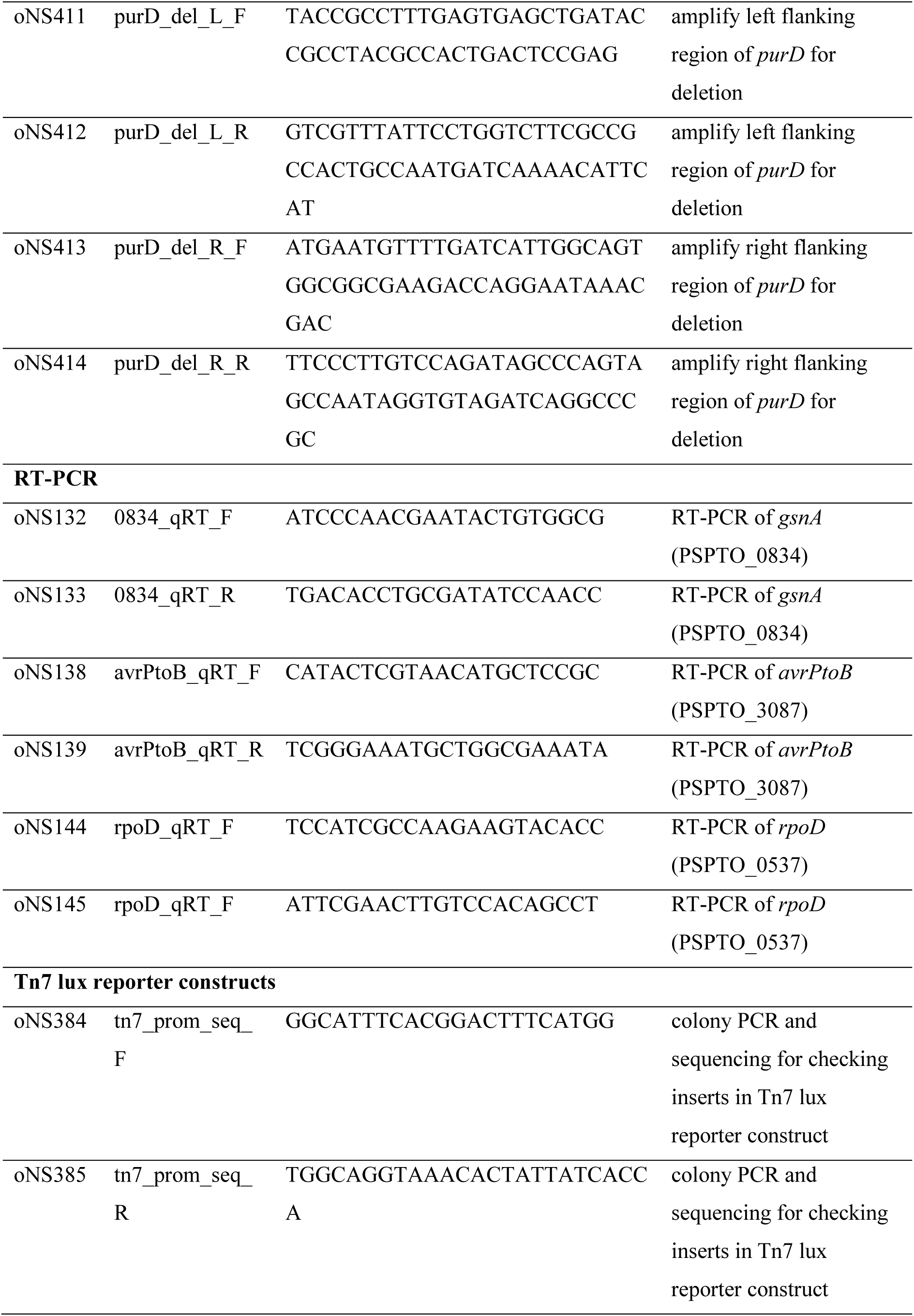
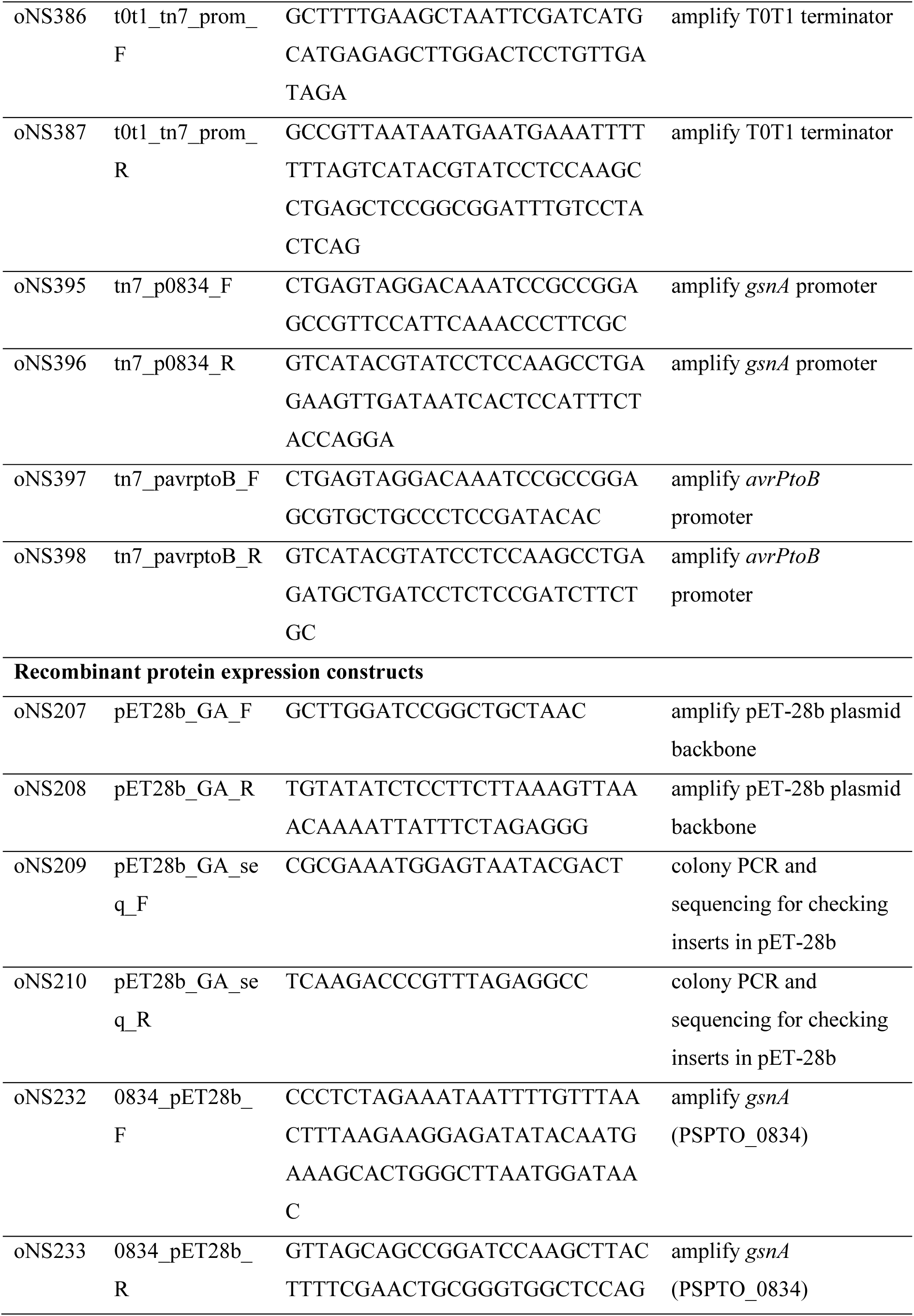
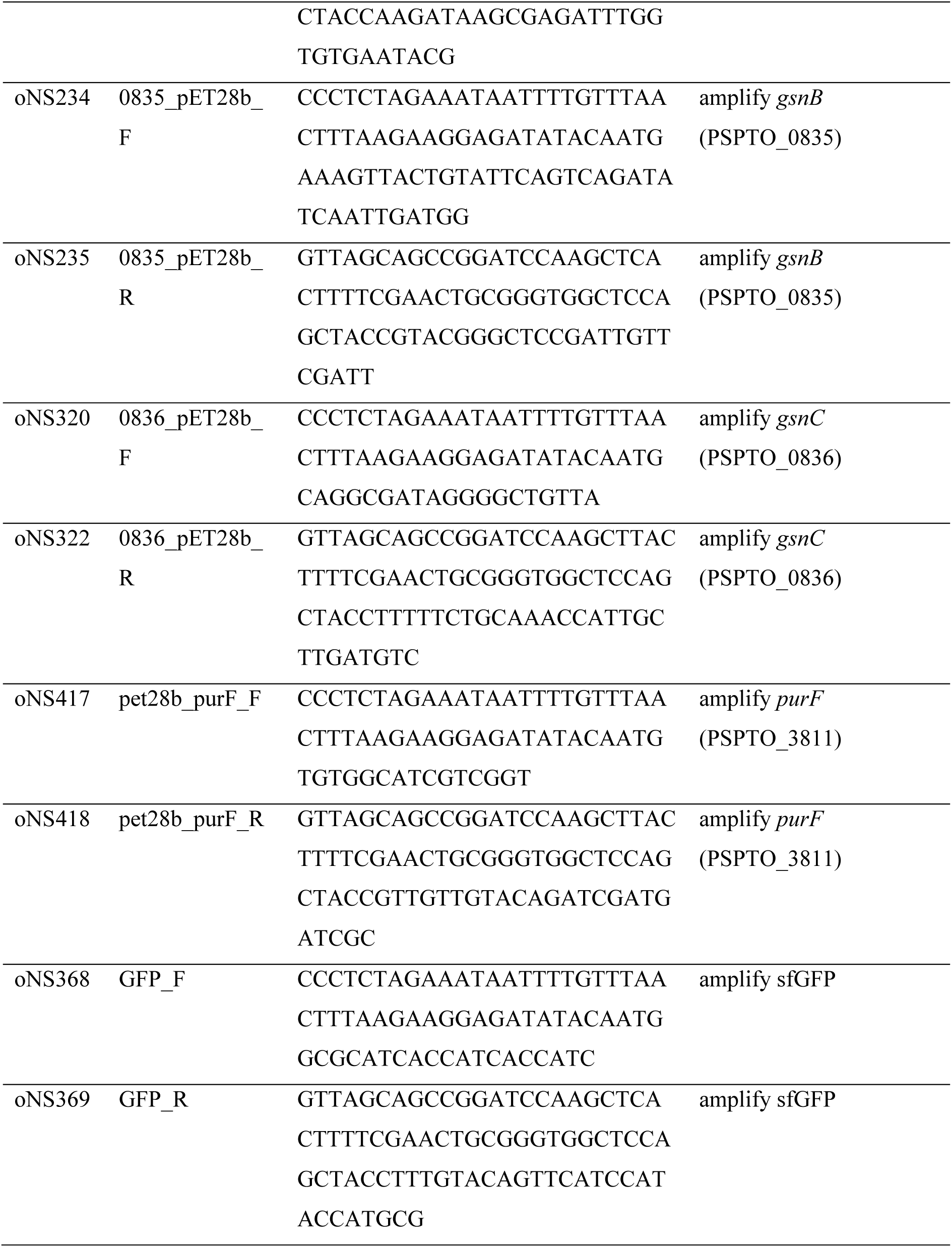
Primers used in this study.

**Table S6.** Cryo-EM data collection, refinement and validation statistics.

|  | Native WT<br>(EMDB-19182)<br>(PDB 8RI7) | *Δgsn*<br>(EMDB-19181)<br>(PDB 8RI6) | Synthetic galactosyrin<br>(EMDB-19183)<br>(PDB 8RI8) |
| --- | --- | --- | --- |
| **Data collection and processing** |  |  |  |
| Magnification (nominal) | 105k | 105k | 130k |
| Voltage (kV) | 300 | 300 | 300 |
| Detector | Gatan K3 | Gatan K3 | Falcon 4 |
| Energy filter | BioQuantum<br>20 eV slit | BioQuantum<br>20 eV slit | Selectris X<br>10 eV slit |
| Electron exposure (e<sup>-</sup>/Å<sup>2</sup>) | 40 | 40 | 40 |
| Defocus range (μm) | -0.5 to -2.1 | -0.5 to -2.1 | -0.5 to -2.1 |
| Pixel size (Å) | 0.829 | 0.831 | 0.921 with Super-resolution |
| Symmetry imposed | D2 | D2 | D2 |
| Images (no.) | 15,180 | 11,944 | 10,166 |
| Final particle images (no.) | 659,661 | 601,662 | 588,492 |
| Map resolution (Å) | 1.93 | 2.06 | 1.42 |
| FSC threshold | 0.143 | 0.143 | 0.143 |
| **Refinement** |  |  |  |
| Initial model used (PDB code) | De novo | De novo | De novo |
| Map sharpening *B* factor (Å<sup>2</sup>) | -56.4061 | -55.5941 | -43.7367 |
| Model composition |  |  |  |
| Non-hydrogen atoms | 33715 | 33662 | 33718 |
| Protein residues | 4064 | 4064 | 4064 |
| Ligands | 1 | 1 | 1 |
| *B* factors (Å<sup>2</sup>) |  |  |  |
| Protein | 5.44 | 14.73 | 13.79 |
| R.m.s. deviations |  |  |  |
| Bond lengths (Å) | 0.006 | 0.003 | 0.008 |
| Bond angles (°) | 0.834 | 0.626 | 1.009 |
| Validation |  |  |  |
| MolProbity score | 1.69 | 1.42 | 1.81 |
| Clashscore | 5.96 | 3.55 | 6.09 |
| Poor rotamers (%) | 1.29 | 1.03 | 1.90 |
| Ramachandran plot |  |  |  |
| Favored (%) | 96.00 | 96.10 | 96.10 |
| Allowed (%) | 3.58 | 3.88 | 3.61 |
| Disallowed (%) | 0.42 | 0.02 | 0.30 |

